# Statistical tests for bivariate spatial association across multi-omics data with disjoint coordinates

**DOI:** 10.64898/2026.06.19.732250

**Authors:** Stijn Hawinkel, Wangjun Hu, Britta Velten, Steven Maere

**Affiliations:** VIB Center for Plant Systems Biology, Technologiepark 71, 9052, Ghent, Belgium; Department of Plant Biotechnology and Bioinformatics, Ghent University, Technologiepark 71, 9052, Ghent, Belgium; Centre for Organismal Studies, Heidelberg University, Im Neuenheimer Feld 230, 69120, Germany; Interdisciplinary Center for Scientific Computing, Heidelberg University, 69120, Germany; Cluster of Excellence GreenRobust, Heidelberg University, 69120, Germany

## Abstract

Spatial biology has entered a new era of multimodal profiling, with multiple, high-dimensional spatial omics types being measured on consecutive tissue slices, or co-assayed on the same slice. Interest then lies in statistical testing for spatial association between the features of the different modalities, to gain insight in biological processes. One major challenge is the multitude of bivariate combinations, leading to high computational demands. Another difficulty is the difference in spatial resolution between technologies, implying no one-to-one matching between the measurement spots of the different modalities, even after alignment. As a result, common statistical measures such as joint distributions and correlations are not defined, and tests need to rely on spatial vicinity only. Moreover, we argue that many existing bivariate association tests address an inappropriate null hypothesis, or make inappropriate assumptions, both implying absence of spatial autocorrelation in any of the features and leading to misleading conclusions. As a remedy, we modify tests for the detection of spatially variable genes (Moran’s I, Gaussian processes and generalized additive models) to derive bivariate spatial association tests across modalities with non-overlapping coordinate sets, and provide variance estimators that do account for spatial autocorrelation. We develop inference methods for single sections as well as for replicated experiments with multiple sections, and compare their performance in nonparametric and parametric simulations. Finally, we apply the newly developed methods to two co-assayed spatial transcriptomics and metabolomics datasets from mouse and human. The full suite of tests is available from github.com/sthawinke/sbivar as the R-package *sbivar*.

**Author summary:** Spatial biology investigates molecules and cells in their spatial context in living tissues. For this purpose, recent technologies measure different classes of biomolecules on the same or adjacent tissue sections. Here we focus on methods to detect colocalisation of molecule pairs, which may provide clues on their biological function, e.g. involvement in the same pathways. Many existing tests for identifying such colocalized pairs rely on the two technologies being measured on the exact same locations, which is often not the case due to differences in resolution. Hence they first require location matching by granulating the technology with the best resolution to that of the worst, discarding information and failing to exploit the full spatial resolution of the data. Moreover, existing methods ignore spatial patterning or autocorrelation naturally present in spatial omics data, leading to many false findings. As a remedy, we present a suite of scalable methods that work directly on disjoint coordinate sets and do account for spatial autocorrelation, and confirm their validity in simulation studies. Next we apply the new methods to metabolite and gene expression measurements of mouse and human datasets, uncovering interesting colocalized gene-metabolite pairs. The methods presented are available in the *sbivar* R-package from github.com/sthawinke/sbivar.

## Introduction

Current biotechnology permits to study the molecular composition of biological tissue at cellular resolution, yielding insights in the spatial organisation of living tissues [1, 2]. Different spatial omics assays exist that each target a single class of biomolecules, e.g. RNA, proteins or metabolites. To obtain a more comprehensive view, different spatial omics technologies are increasingly being applied to the same tissue, either on consecutive slices [3, 4], or by measuring the different molecule types or modalities consecutively on the same slice (co-assaying) [5–9]. Although single molecules can be detected using hybridisation [10], here we focus on technologies that quantify local molecular composition on a predefined grid of spots, such as the 10x Genomics Visium platform for spatial transcriptomics [11] or matrix-assisted laser desorption/ionisation mass spectrometry imaging (MALDI-MSI) for spatial metabolomics [12]. To discover statistical associations between the different classes of biomolecules, a pivotal first step is to align the images of the different modalities profiled onto a common coordinate framework (CCF) to make corresponding regions of the original tissue overlap. This alignment problem is mostly solved manually, sometimes with the help of hematoxylin and eosin (H&E) stainings of the different images [3, 9, 13]. Given a CCF, the relative positions of the spots of the different modalities are known, allowing the two modalities to be analysed jointly. In this work, we focus on bivariate spatial association tests across modalities, as bivariate spatial association can provide biological insights, even though it does not prove physical interaction or functional coupling between molecules. Bivariate spatial association tests pose a unique analysis problem, as even after alignment there is no one-to-one mapping between the spots from different modalities, i.e. the coordinate sets are disjoint due to differences in resolution and measurement area between the technologies. As a result, common statistical quantities such as correlations and joint distributions are undefined. Current practice is therefore to aggregate the spots of the modality with the best resolution and pair them with spots of the other modality to remove the disjointness [3, 8, 9], but this results in information loss. An additional challenge is computational, as the number of bivariate tests greatly exceeds the number of univariate tests applied to each modality separately.

Finally, the null hypothesis should be chosen carefully. One option is the randomisation null hypothesis, which means that all observations are equally likely to occur at any location, implying an absence of spatial patterning or spatial autocorrelation (SAC) in any of the two variables involved in the test. The randomisation null distribution can be described analytically [14–16], or be approximated by permuting the observations over their measurement locations [17, 18]. Rejection of the randomisation null hypothesis indicates *either* bivariate spatial association, *or* SAC in one of both variables, *or* both. More formally, for two quantitative measurements *x* and *y* from different modalities, the randomisation null hypothesis entails all of the following:

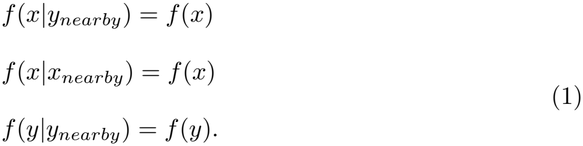

with *f* a density and *x*|*y*_*nearby*_ indicating *x* conditional on nearby values of *y*, including y-values measured at the same location as *x*. The result of such omnibus test is hard to interpret, and calling it a test for bivariate spatial association is at best misleading. We argue that it is preferable to have separate tests for univariate spatial patterning only, and for bivariate spatial association only. Many good tests for the former exist already, in particular for detecting spatially variable genes (SVGs) in transcriptomics, based on e.g. univariate Moran’s I (*MERINGUE* [17]), Gaussian processes (*SPARK* [19] and *SpatialDE2* [20]), generalized additive models (*spVC* [21]) and nonparametric tests for association between distance and covariance (*SPARK-X* [22]). For that reason, we focus on tests for bivariate association only, and call the associated null hypothesis the independence null hypothesis, entailing only the first condition in Eq. 1. Many existing methods attempt to test the independence null hypothesis, but make inappropriate assumptions, in practice testing the randomisation null hypothesis instead. Standard inference on Pearson correlation after spot matching [3, 8, 9] and some versions of bivariate Moran’s I [23, 24] explicitly assume i.i.d. observations. Tests based on permutations, e.g. *SpaGene* [18], *MERINGUE* [17] and *Liana+* [25], implicitly assume exchangeability of observations [26]. Yet both the i.i.d. and exchangeability assumptions are violated in presence of SAC in either variable, which can lead to erroneous inference.

Other existing methods do achieve valid inference on the independence null hypothesis by specifically accounting for SAC. The modified t-test for the Pearson correlation coefficient tests the independence null hypothesis by estimating SAC for every variable separately and decreasing the degrees of freedom in the t-test accordingly [27–29], but is only applicable to joint coordinate sets. Custom permutation schemes have been developed that leave the SAC intact and thus test the independence null hypothesis for any test statistic: the so-called “random-shift” tests [30, 31]. Similarly, [32] permute one of both variables and then reintroduce SAC with the help of the variogram of the observed variable to construct synthetic null datasets. Yet such tests can be computationally demanding and their p-values are discretized by the number of permutations or synthetic datasets built, which can result in low power after multiple testing correction. The SpaceBF method explicitly accounts for SAC in a Bayesian framework [33], but requires joint coordinates and relies on Markov chain Monte-Carlo sampling which is computationally demanding for large number of feature combinations.

In this work, we evaluate existing methods for detection of bivariate spatial association using nonparametric and parametric simulations, and develop scalable new statistical tests for the independence null hypothesis of bivariate association across disjoint coordinates, available in the R-package *sbivar* from github.com/sthawinke/sbivar. As demonstration, we apply the new tests to spatial multi-omics datasets from mouse [9] and human [34] tissues.

## Materials and methods

### Algorithms

Given are outcome matrices **X**_*n×p*_ and **Y**_*m×r*_ of the two modalities, with *n* and *m* the number of spots and *p* and *r* the number of features, and corresponding coordinate matrices **C**_*n×*2_ and **E**_*m×*2_ in a CCF. For ease of notation, we shift and scale the coordinate matrices to the unit square. We adapt three methods, bivariate Moran’s I, Gaussian processes (GPs) and generalized additive models (GAMs), to test for bivariate spatial association as follows.

### Bivariate Moran’s I under the independence null

Bivariate Moran’s I is a measure of bivariate spatial association [14, 15] that naturally generalizes to the disjoint coordinate case. Define a weight matrix **W**_*n×m*_ with positive elements *w*_*ij*_, usually decaying with distance between **c**_*i*_ and **e**_*j*_. We restrict the discussion to a single feature pair with observation vectors **x** and **y**, which are mean-centered and scaled such that 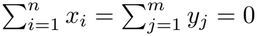 and 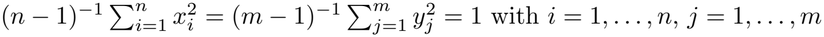 . Bivariate Moran’s I is defined as 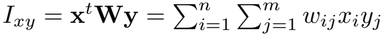 (see Fig 1a). Under the independence null hypothesis, *E*(*I*_*xy*_) = 0 (see Section 1.2.1 in S1 Text). The variance 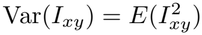 can be written as

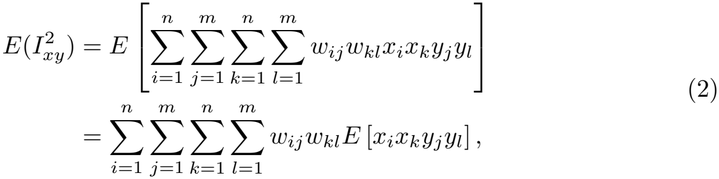

given that the grid locations, and thus the weights *w*_*ij*_, are fixed. Under the independence null hypothesis, *x* and *y* are independent, such that *E* [*x*_*i*_*x*_*k*_*y*_*j*_*y*_*l*_] = *E* [*x*_*i*_*x*_*k*_] *E* [*y*_*j*_*y*_*l*_] + Cov(*x*_*i*_*x*_*k*_, *y*_*j*_*y*_*l*_) = *E* [*x*_*i*_*x*_*k*_] *E* [*y*_*j*_*y*_*l*_], with the expectations *E* [*x*_*i*_*x*_*k*_] and *E* [*y*_*j*_*y*_*l*_] determined by SAC and the distances *d*_*ik*_ and *d*_*jl*_ only. Plugging in such estimates of the SAC 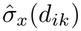 and 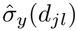, Eq. 2 can be rewritten to estimate the variance as 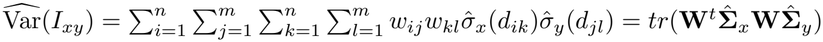,with *tr*(·) the trace function and 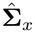 and 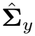 estimated autocorrelation matrices, ignoring the effects of centering *x* and *y* as *n* and *m* are large. Any autocorrelation estimator could be used, for computational speed we choose Matheron’s variogram estimator [35] followed by linear or exponential variogram model fitting using the *fit*.*variogram* function in the *gstat* R-package [36], fixing the sill to 1. Unrealistically assuming *x* and *y* are i.i.d. yields *tr*(**W**^*t*^**W**) as variance estimate, as given previously [23, 24]. Since SAC is usually positive, it inflates the variance of the Moran’s I estimator with respect to the i.i.d. case. If the coordinate sets are joint (**C** = **E**) and only pairs measured at the same location are assigned non-zero, equal weights (**W** = **I** with **I** the identity matrix), then *I*_*xy*_ is proportional to the Pearson correlation with variance estimator 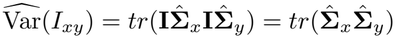 . This latter expression is proportional to the variance of the Pearson correlation estimate in the modified t-test (see Section 1.1.2 in <u>S1 T</u>ext), which is thus a special case of bivariate Moran’s I under the independence null with joint coordinates (see Table 1 for an overview). Statistical testing on *I*_*xy*_ proceeds by assuming that ^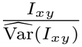^ is asymptotically standard normal, in line with theoretical results on univariate Moran’s I [37] and our empirical findings (Fig S1). *I*_*xy*_ is bounded in absolute value by 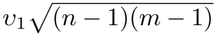, with *v*_1_ the first singular value of ***W*** (see Section 1.2.1 in <u>S1 T</u>ext). The choice of weight matrix determines which type of spatial association patterns bivariate Moran’s I can detect. To maintain power against many possible patterns but still provide a single p-value, we test several weight matrices, and combine the resulting p-values into one using the Cauchy combination test for dependent p-values [38]. Many weight matrices are possible, here we opt for weights proportional to 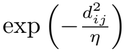 with *η ∈* [5 *×* 10^*−*6^, 2 *×* 10^*−*4^, 2 *×* 10^*−*2^] to detect interactions up to 20% of the largest dimension of the image (see Fig S2).

**Table 1.**
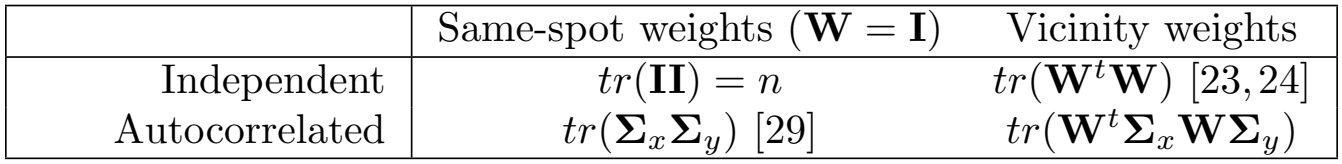
Variances of bivariate Moran’s I *I*_*xy*_ = **x**^*t*^**Wy** for normalized variables *x* and *y* under different weight matrices (columns) and dependence structures of observations (rows) for large *n* and *m*. The first column is only applicable to joint coordinate sets, in which case *I*_*xy*_ is proportional to Pearson correlation.

|  | Same-spot weights (**W** = **I**) | Vicinity weights |
|---|---|---|
| Independent | *tr*(**II**) = *n* | *tr*(**W**<sup>*t*</sup>**W**) [23, 24] |
| Autocorrelated | *tr*(**Σ**<sub>*x*</sub>**Σ**<sub>*y*</sub>) [29] | *tr*(**W**<sup>*t*</sup>**Σ**<sub>*x*</sub>**WΣ**<sub>*y*</sub>) |

**Fig 1.**
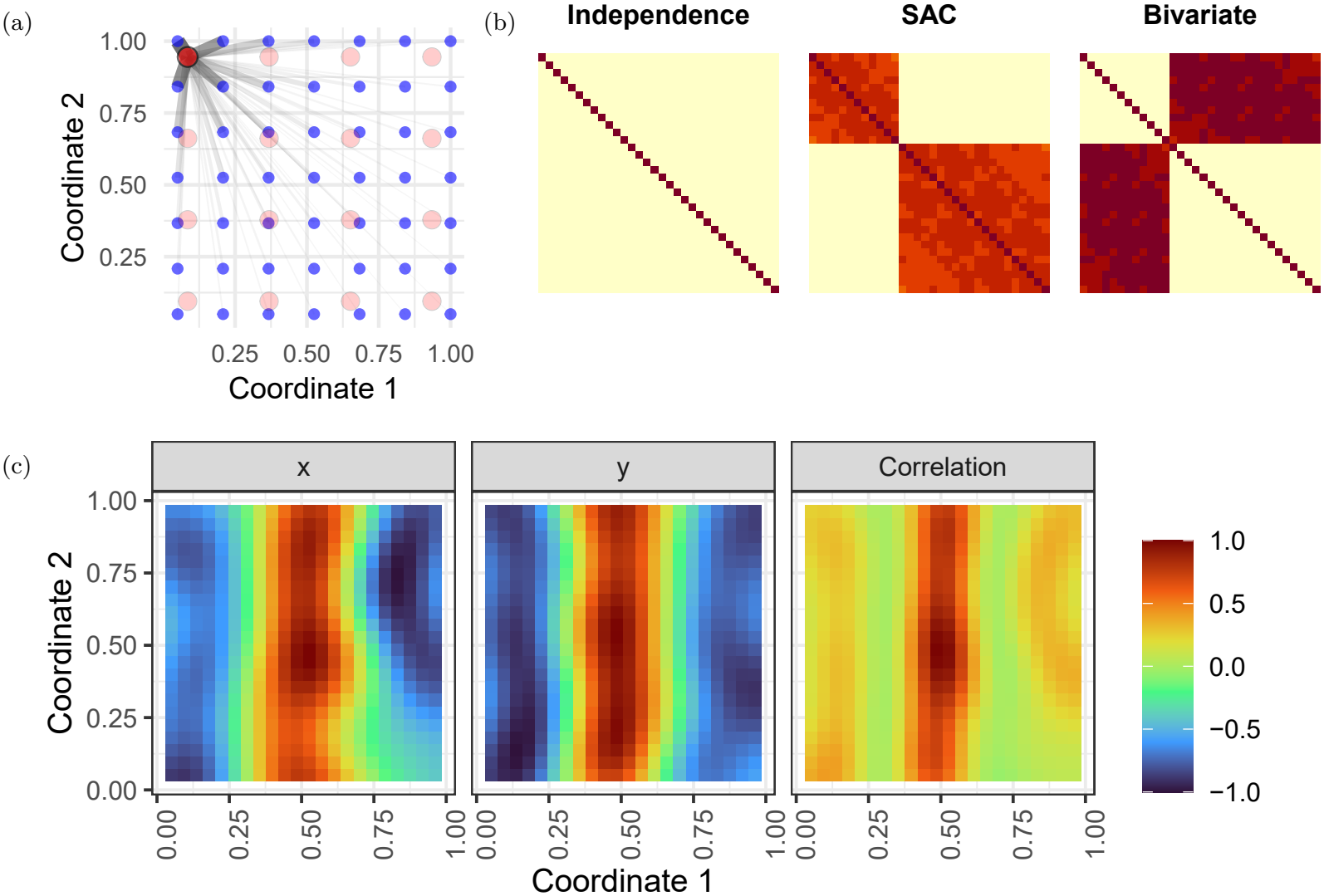
Illustration of the three new methods for detecting bivariate spatial association implemented in *sbivar* . (a) Visualisation of one term in the calculation of bivariate Moran’s I for disjoint coordinate sets: red and blue dots represent measurement spots from modalities with low and high resolution, the widths of the grey lines between them reflect the weights *w*_*ij*_. (b) The three components of the variance-covariance matrix of the bivariate Gaussian process in Eq. 4. The bivariate spatial association test revolves around the bivariate covariance matrix on the right being zero. (c) Estimated spline surfaces of GAMs for simulated outcomes of x and y variables forming a “streak” pattern (left and middle column), and contributions to the correlation between spline surfaces (right column), all scaled to the [-1,1] range for legibility.

### Bivariate Gaussian processes

A second method is based on Gaussian processes (GPs), and we call it “bivariate GPs” as it models the concatenation of x and y values, indicated by the vector **z** of length *n* + *m* = *h*, through a multivariate normal (MVN) distribution:

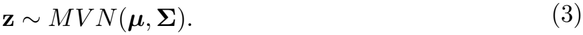

The mean vector ***µ*** accounts for differences in mean between *x* and *y*. The covariance matrix **Σ** is composed of the following three covariance matrices: i.i.d. noise in *x* and *y* (diag(***σ***^2^)), SAC within *x* and *y* (**Σ**^*SAC*^) and spatial covariance between x and y (**Σ**^*biv*^):

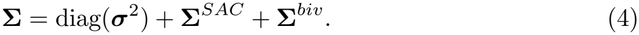

***σ***^2^ is a vector of noise variances, **Σ**^*SAC*^ a block diagonal matrix of matrices 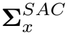 and 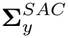 describing SAC within *x* and *y*, and **Σ**^*biv*^ models the spatial association of interest (see Fig 1b). Both SAC and spatial covariance are assumed to decay with distance, which we model through the isotropic Gaussian covariance function 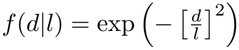, with *d* the distance between observations and *l* the length scale. Hence 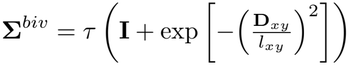 with *τ* the variance of the random effect engendering the bivariate association, *l*_*xy*_ its length scale and **D**_*xy*_ an *h × h* matrix with distances between **c** and **e** on the off diagonal blocks and *∞* for distances within **c** and **e** (see the right plot in Fig 1b). The null hypothesis of interest is that *τ* = 0, which can be tested using a score test statistic U [39]:

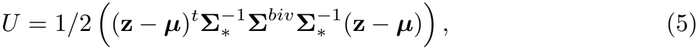

with **Σ**_*\**_ = diag(***σ***^2^) + **Σ**^*SAC*^. Under the null, *U* follows a scaled chi-squared distribution [20, 39] (see Section 1.2.2 in S1 Text for details). A score test is indicated as the Wald and likelihood ratio test statistics have degenerate null distributions when the null value is on the edge of the parameter space [40], as is the case here for *τ* . The score test has the additional advantage of not having to estimate all full models, i.e. all *pr* matrices **Σ**^*biv*^, but only *p* + *r* matrices 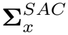 and 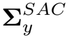of the null model. Yet the test as formulated can only detect positive associations, so we perform an additional one with 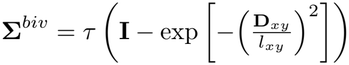, and take two times the smallest of both p-values as final p-value to perform a two-sided test. This leaves the question how to construct **Σ**^*biv*^ if *l*_*xy*_ is not estimated. Following SPARK [19] and SpatialDE2 [20], we test a series of values for *l*_*xy*_, defined as the exponential of an equally spaced series of length five ranging from the logarithm of the 0.5th to the logarithm of the 50th percentile of all observed distances between spots. The resulting p-values of the five two-sided tests are again combined with the Cauchy combination test [38].

### Generalized additive models (GAMs)

As a third method, GAMs can be employed to test for bivariate association. Outcomes *x* and *y* are modeled as a function of space through 2D thin plate smoothing splines *s*_*x*_ and *s*_*y*_:

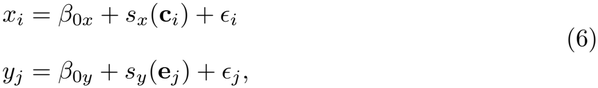

with *β*_0*x*_ and *β*_0*y*_ baselines. *ϵ*_*i*_ and *ϵ*_*j*_ are error terms with zero-mean normal distributions that can either be assumed i.i.d. or spatially autocorrelated to absorb residual autocorrelation not captured by the smoothing splines. The models in Eq. 6 are then fitted by the *gam* or *gamm* functions in the *mgcv* package [41], and the corresponding method referred to as “GAMs” or “GAMMs” (for generalised additive mixed models), respectively. Define a new set of coordinates as an evenly spaced grid **C**_*B*_ of size *B* within the concave hull spanning both **C** and **E**. Evaluating the thin plate splines in **C**_*B*_ yields ***ι*** = *s*_*x*_(**C**_*B*_) and ***ζ*** = *s*_*y*_(**C**_*B*_), for which we can calculate the covariance *q*_*xy*_ and correlation *ρ*_*xy*_ as

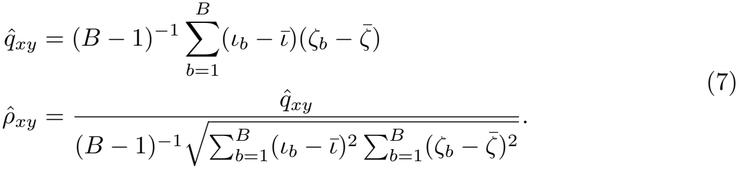

The evaluations of a single spline *s*_*x*_ at different points *b* are dependent as they rely on the same estimated spline parameters, so the regular formulae for the variance of 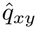 and 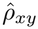 are invalid. Instead, we propagate the uncertainty on the spline coefficients as follows. Call **H**_*x*_ the spline basis of dimension *n × v* and ***β***_*x*_ the spline coefficients of length *v* with variance-covariance matrix ***Φ***_*x*_. Then ***ι*** = *s*_*x*_(**C**_*b*_) = **H**_*x*_***β***_*x*_ are the spline evaluations, with variance-covariance matrix 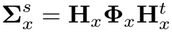 . Using the delta method [42], the variance of 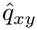 can be approximated as:

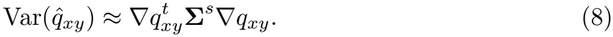

The gradient 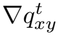 of length 2*B* equals 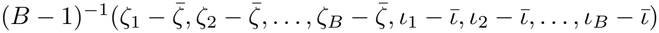. **Σ**^*s*^ is a block diagonal matrix of 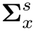 and 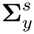 : the evaluations *ι*_*b*_ and *ζ*_*b*_ are independent, since the splines were estimated on different outcome vectors. Given this variance estimate 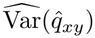, we can test if 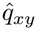is significantly different from 0 using a z-test. Splines allow for easy visualisation of the individual spatial patterns, and 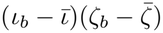 can be plotted as a function of space to discover areas contributing most to the overall spatial association (see Fig 1c). GAMs naturally generalize to other link functions and outcome distributions (see Section 1.2.3.1 in S1 Text). Note that the three methods presented detect departures in the first two moments, mean and covariance. Rejection of their corresponding null hypothesis implies that the first condition in Eq. 1, and hence the independence null hypothesis, does not hold.

### Multi-image analysis

Next we extend the methods described for single-image analysis to datasets with multiple images, e.g. tissue slices from different biological replicates [3, 9]. For the model based methods (bivariate GPs and GAMs), the ideal approach would accommodate all images in a single model, but this solution is computationally demanding and requires many additional assumptions on how e.g. spatial covariance parameters are allowed to vary across replicates. Here we take the pragmatic approach of extracting a single measure of association strength from each image, and using it as outcome in a linear model accounting for the design structure [43, 44]. This implies that variability is quantified across replicates rather than within a single image. Nevertheless, we investigated if it is beneficial to inverse weigh the extracted association measure, bivariate Moran’s I or the correlation of evaluated spline surfaces *ρ*_*xy*_, by their estimated variance in the linear model to improve the power. In addition, we tried scaling bivariate Moran’s I by its maximum absolute value for comparability. For the bivariate GPs, no measure of association strength is estimated, only a test statistic and corresponding p-value, so we do not use it for replicated analysis.

### Simulation studies

#### Nonparametric simulations

In order to assess type I error control, we perform the following nonparametric simulations based on the datasets by [8] and [9], referred to as the Godfrey and the Vicari data. Both datasets contain spatial transcriptomics and metabolomics measurements on the same tissue slices. The Godfrey data consists of human lung and breast cancer tissue, the Vicari data of three mouse brains, lesioned unilaterally with 6-hydroxydopamine (6-OHDA) to emulate Parkinson’s disease, measured in triplicate. The spots of the Godfrey data have been matched by the authors by aggregating the metabolomics spots to a lower resolution (Fig S26), whereas the coordinates of the Vicari data are disjoint and these we aligned with the MAGPIE package, pipeline version ‘noHE’ with affine transformation [13] (Fig S34, chosen landmarks are available in S1 Data). For the Vicari data, transcriptomics spots with fewer than 1,000 reads were filtered out, for the metabolomics modality and both modalities of the Godfrey data, spots with all zero values were omitted from the analysis. For both datasets, we repeatedly permute the coordinates of one of both modalities or both of them, and apply tests for bivariate spatial association to all pairs of the 30 most abundant features per modality. We included our implementation of bivariate Moran’s I (”Moran’s I (independence) [sbivar]”) and GAMs (”GAMs [sbivar]”) in this comparison, but not bivariate GPs or GAMMs for computational reasons. For fitting GAMs, we use the gamma distribution to model the metabolite data, after augmenting all values by a pseudocount of 10^*−*8^, and the negative binomial distribution for the transcriptome data. In both cases, we employ a log link function and use the spot sums (library sizes or total ion counts) as offsets. For the other methods, the observations were normalized to relative abundances by dividing by the spot sums, as in the original analyses. As existing analysis methods, for the Godfrey data with joint coordinate sets, we use bivariate Moran’s I with the variances supplied by [15] (”Moran’s I (randomisation)”) and by [24] in their *SpatialDM* method (”Moran’s I (SpatialDM)”), and with permutation null (”Moran’s I (permutation)”) and the random shift null distribution ( “Moran’s I (random shift)”, see Section 1.2.1.6 in S1 Text for details) [30, 31], and further Lee’s L (“Lee’s L (randomisation)”, *lee*.*test* in the *spdep* R-package) [16], bivariate Geary’s C with permutation null distribution (”Geary’s C (permutation)”, code based on *geary*.*bi* in the *bispdep* R-package) [45], SpaGene by [18] (”SpaGene”, *SpaGene LR* in the *SpaGene* R-package), Pearson correlation (“Pearson correlation”, *cor*.*test* in the *stats* R-package) and the modified t-test (”Modified t-test”, *modified*.*ttest* in the *SpatialPack* R-package) [27]. Where applicable, the weight matrices for the aforementioned methods where constructed as for bivariate Moran’s I above with *η* = 2 *×* 10^*−*4^. 300 permutations were run for the permutation-based methods. For the disjoint coordinate sets of the Vicari data, we use GAMs and bivariate Moran’s I with our independence null, with regular permutation and with random shift null distributions. In addition, false discovery rate (FDR) control at a nominal level of 0.05 is assessed after adjusting the p-values using the Benjamini-Hochberg procedure [46].

### Parametric simulations

Next, to also probe the power to detect bivariate associations, we perform the following parametric simulation study. Observations are generated on evenly spaced grids, with joint coordinate sets with n=m=400, and also with disjoint coordinate sets with n=400 and m=900, and boundaries 0 and 1 for **C** and 0.03 and 1.03 for **E** (Fig S3). The concatenation **z**^*t*^ = (**x**^*t*^, **y**^*t*^) of outcome variables *x* and *y* is drawn from a multivariate Gaussian distribution: **z** *∼ MV N* (***µ*, Σ**). We investigate the following null and alternative scenarios with spatial dependence structure introduced in the mean or covariance. The null scenarios are:

- “No spatial pattern”: ***µ*** = **0** and **Σ** = **I**
- “Independent GPs”: ***µ*** = **0** and **Σ** block diagonal with 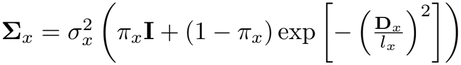and _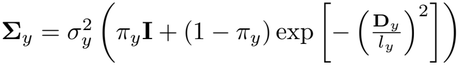_, with **D** and **D** distance matrices within **C** and **E**, respectively, *π*_*x*_ = *π*_*y*_ = 0.25 the nuggets, *l*_*x*_ = *l*_*y*_ = 0.15 the length scales and 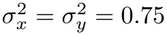 the variances.

For the following alternative scenarios, one fifth of the features are generated as in “No spatial pattern” above, the other four fifths as:

- “Dependent GPs”: ***µ*** = **0** with **Σ**_*x*_ and **Σ**_*y*_ as for independent GPs. **Σ**^*biv*^ is constructed as 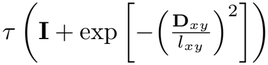as in the description of “bivariate GPs”, and added to the covariance matrix described for the “independent GPs” scenario to yield the final **Σ**, with *l*_*xy*_ = 0.15 and *τ* = 0.15.
- “Streak”: 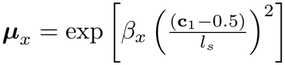 and 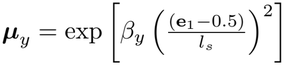, with **c** and **e**_1_ the first columns of the coordinate matrices, i.e. the x-coordinates, *β*_*x*_ and *β*_*y*_ drawn uniformly on [0.75, 1], *l*_*s*_ = 0.15 and **Σ** = **I**.
- “Zones”: 6 random points are drawn independently and uniformly within the area boundaries, and distances from the grid to these points are calculated as the *n ×* 6 matrix **D** . The mean is defined as ***µ*** = 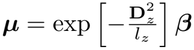, with the elements of ***β*** drawn uniformly on [0.75, 1], *l*_*z*_ = 0.05 and **Σ** = **I**.

For the final two alternative scenarios, one fifth of the features are generated as in “Independent GPs” above, the other four fifths as:

- “Streak + GPs”: ***µ*** as for “Streak” and **Σ** as for “Independent GPs”.
- “Zones + GPs”: ***µ*** as for “Zones” and **Σ** as for “Independent GPs”.

The designs are visualized in Fig S4, the resulting amounts of SAC as measured by the univariate Moran’s I in Fig S5. 50 Monte-Carlo simulations with ten features per modality are run per scenario. As now spot numbers are fewer, we also employ GAMMs (“GAMMs [sbivar]”) and bivariate GPs (”Bivariate GPs [sbivar]”) on top of the methods applied in the nonparametric simulation study. Moreover, as now observations are available on a regular grid, also the random shift test (*CC*.*test* in the *NTSS* R-package) with Kendall’s correlation coefficient and correction = “variance” (”Kendall’s correlation (random shift)”) [30] is included. An overview of all methods compared and their properties is given in Table 2. For disjoint coordinates, all methods requiring joint coordinate sets were applied after matching the observations in **E** to their nearest neighbours in **C** using the *nn*2 function in the *RANN* R-package [47], effectively reducing the number of observations of the second modality to 400, following [9]. Type I error and power are calculated as the proportion of raw p-values below the significance level of 0.05 for the null features respectively the features with true association. The FDR was calculated as the proportion of false findings among feature pairs with adjusted p-values below 0.05, and set to 0 if there were no such findings. In addition, we set up a simulation study with replicated designs, simulating each time nine replicates of the aforementioned spatial patterns on disjoint coordinate sets, with per replicate a 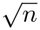 drawn from a negative binomial distribution with mean 25, and a 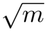 from a negative binomial distribution with mean 30, both with overdispersion parameter 0.2, and with a minimum of 100 for *n* and 400 for *m*. Pearson correlation was calculated after nearest-neighbour matching as above, bivariate Moran’s I *I*_*xy*_ and the correlation *ρ*_*xy*_ between GAMs’ spline surfaces were calculated directly on the disjoint coordinate sets and all are used as outcome values of linear models. GAMMs were not employed as their spline surfaces differ little from the regular GAM ones. We investigate whether inverse weighting by the variance of bivariate Moran’s I and GAM correlation estimates in the linear model, and scaling bivariate Moran’s I statistic by its maximum value, improve the inference.

**Table 2.** Methods to test for bivariate spatial association considered in this study, with the null distributions shown in parentheses; [sbivar] indicates methods proposed or improved in this work and available in the *sbivar* R-package. The other columns show the null hypothesis tested, the estimand, how the methods account for spatial autocorrelation (SAC) and whether they accept disjoint coordinates.

| Method | Null hypothesis | Estimand | SAC structure | Ingests disjoint coordinates |
|---|---|---|---|---|
| Moran’s I (randomization) | No SAC or bivariate spatial association | Moran’s I | None | No |
| Moran’s I (permutation) | No SAC or bivariate spatial association | Moran’s I | None | Yes |
| Moran’s I (*SpatialDM*) | No SAC or bivariate spatial association | Moran’s I | None | No |
| Geary’s C (permutation) | No SAC or bivariate spatial association | Geary’s C | None | No |
| *SpaGene* | No SAC or bivariate spatial association | Subnetwork connectivity | None | No |
| Lee’s L (randomisation) | No SAC or bivariate spatial association | Lee’s L | None | No |
| Pearson correlation | No SAC or bivariate spatial association | Pearson correlation | None | No |
| Random shift tests | No bivariate spatial association | Any statistic | SAC preserved in random shifts | Possibly |
| Modified t-test | No bivariate spatial association | Pearson correlation | Variogram | No |
| Bivariate Moran’s I (inde-pendence) [sbivar] | No bivariate spatial association | Moran’s I | Variogram | Yes |
| Bivariate GPs [sbivar] | No bivariate spatial association | Spatial covariance | SAC matrix | Yes |
| GAMs [sbivar] | No bivariate spatial association | Correlation of mean surfaces | Thin plate spline | Yes |
| GAMMs [sbivar] | No bivariate spatial association | Correlation of mean surfaces | Thin plate spline + SAC matrix | Yes |

Finally, we investigate the effect of alignment error on bivariate association inference as follows. We generate 10 features per modality with the spatial patterns described above on the aligned coordinates of the Vicari data, changing parameter values to *π*_*x*_ = *π*_*y*_ = 0.85, *l*_*x*_ = *l*_*y*_ = *l*_*xy*_ = 0.1, *l*_*s*_ = 0.05, *l*_*z*_ = 0.01 and *β* generated uniformly on [0.3, 0.5] to expose the effect of alignment error. Alignment errors are emulated by adding random zero-mean Gaussian variates with standard deviation increasing as [0.5, 1, 1.5, 2, 3] to the chosen landmarks’ coordinates. Next we run the remaining *MAGPIE* pipeline with the distorted landmarks, and analyze the outcomes generated on the original coordinates while providing the distorted coordinates (shown in Fig S6) to the analysis methods. We repeat the landmark distortion 10 times for every sample and standard deviation.

### Real data analysis

The same analyses described above were applied to the individual sections of the Godfrey and Vicari data, and to the lung cancer and breast cancer samples of the Godfrey data, and to the entire Vicari dataset in a multi-image analysis. All detected genes and all identified metabolites were included in the analysis. GAMMs were not employed for computational reasons. For the Vicari data, a random effect for mouse is included in the linear models to account for replicate measures on the same mouse. No random effects were used for the Godfrey data, as all but two sections come from different individuals. To correct for multiple testing, we applied the Benjamini-Hochberg multiplicity correction [46], and adjusted p-values smaller than 0.05 were considered significant.

## Results

### Simulation studies

#### Nonparametric simulations

Fig 2 reveals that all methods control the type I error when the coordinates of both modalities are permuted, as this corresponds to the randomisation null hypothesis. Yet only Moran’s I (randomisation), Moran’s I (random shift), Moran’s I (independence), the modified t-test, Pearson correlation and GAMs also control it when only one modality is permuted, having approximately uniform or larger than uniform p-value distributions (Figs S8-S9). All other methods suffer from inflated type I errors when only one of both modalities is permuted. This is likely because of the SAC remaining in the non-permuted modality, as the type I error is highest when the modality with highest SAC (the metabolome, see Figs S10-S11) is left untouched. Moreover, features with high SAC are more likely to incur false positives for the methods with inflated type I errors (see Fig S12). Moran’s I (randomisation) is found very conservative, presumably suffering from the heteroskedasticity in the data. The FDR is inflated with respect to the nominal level for the same methods as for the type I error, although the violation is not as severe (Fig S13). In a multi-image analysis of the same permuted datasets, per cancer type for the Godfrey data and on all mouse samples for the Vicari data, we found Pearson correlation, bivariate Moran’s I as well as GAMs to control the type I error and FDR at the nominal level (Figs S14-S15), as now the variability of the association measures is correctly quantified across the replicates. An exception are bivariate Moran’s I and GAMs with inverse variance weighting, which have inflated type I errors and FDRs, hence inverse weighting does not seem to be a good strategy when heterogeneity between replicates is a major source of variation [48, 49].

**Fig 2.**
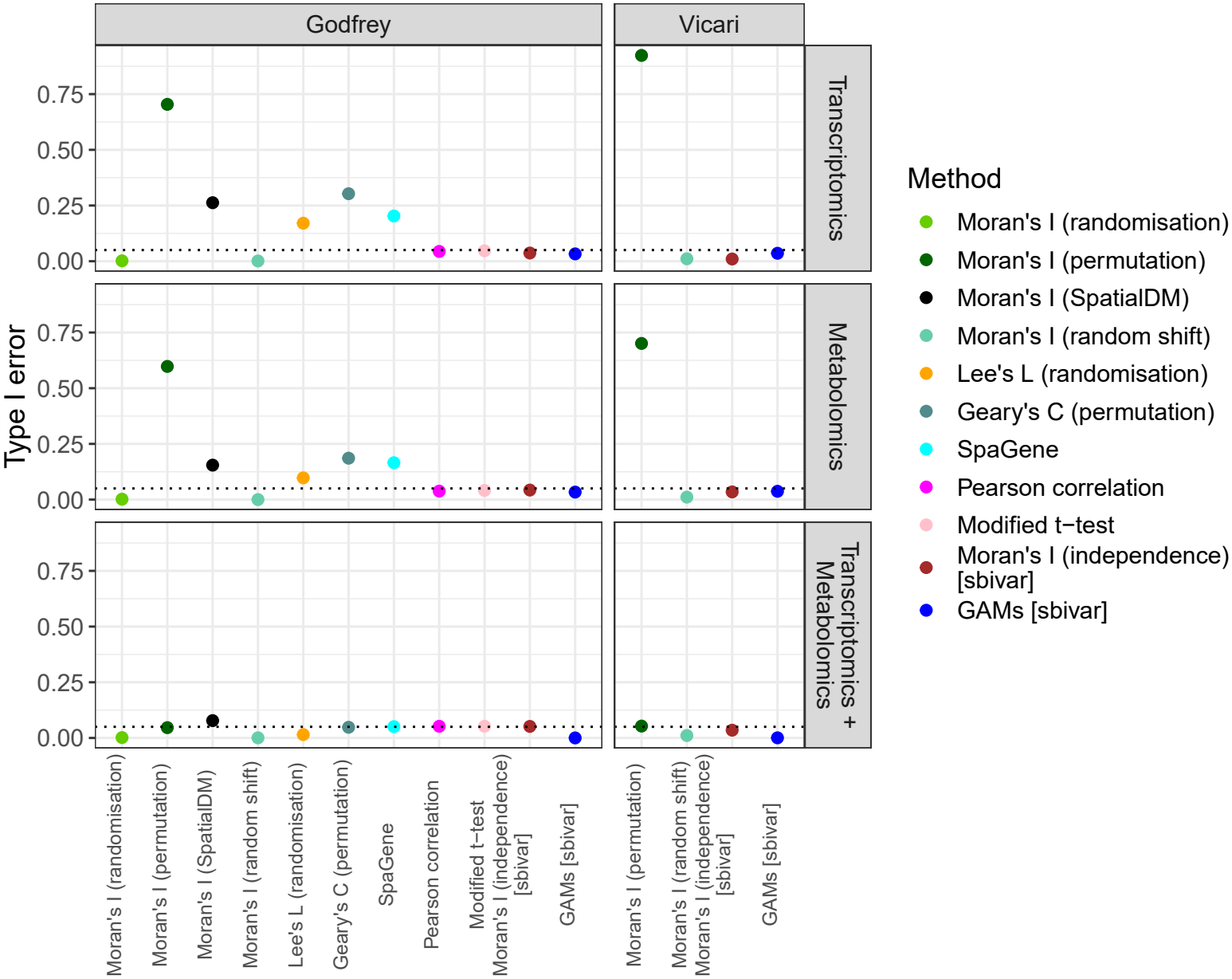
Type I error (y-axis) for different methods with null distributions in parentheses (x-axis, colour) for one or two permuted modalities (rows) averaged over all samples of the Godfrey and Vicari data (columns). The dotted horizontal line indicates the significance level of 0.05, [sbivar] indicates methods introduced in this work.

### Parametric simulations

The simulation results in Fig 3 and Figs S16-S17 show that all methods considered control the type I error of the independence null hypothesis in absence of SAC. Yet many methods have inflated type I errors in presence of SAC, also including Pearson correlation unlike in the nonparametric simulation. This is because Pearson correlation only becomes liberal when both variables exhibit SAC (see Fig S18) [50]. Also GAMs become liberal in presence of SAC, presumably for the same reason, and even GAMMs suffer from slightly inflated type I errors in some scenarios. All methods failing to control type I error yield most false findings when both features exhibit SAC (Fig S18), suggesting that the results of the nonparametric simulation study still underestimate the type I error inflation on real data, as there is SAC left in only one of both modalities. Unlike in the nonparametric scenarios, also Moran’s I (randomisation) suffers from inflated type I error, presumably as the outcomes are now generated in homoskedastic fashion. The results for Moran’s I (permutation) and Moran’s I (SpatialDM) are very similar for joint coordinate sets, having inflated type I errors in all scenarios with background GP noise, confirming that *SpatialDM* presents an analytical version of the randomisation null hypothesis of no SAC or bivariate association. Fig S19 reveals that among the methods controlling the type I error, the modified t-test and bivariate GPs also control the FDR at the nominal level. Yet bivariate Moran’s I (independence) has a slightly inflated FDR with disjoint coordinates, and Kendall’s correlation (random shift), GAMs and GAMMs have excessively large FDRs, likely due to slow convergence to asymptotic null distributions.

**Fig 3.**
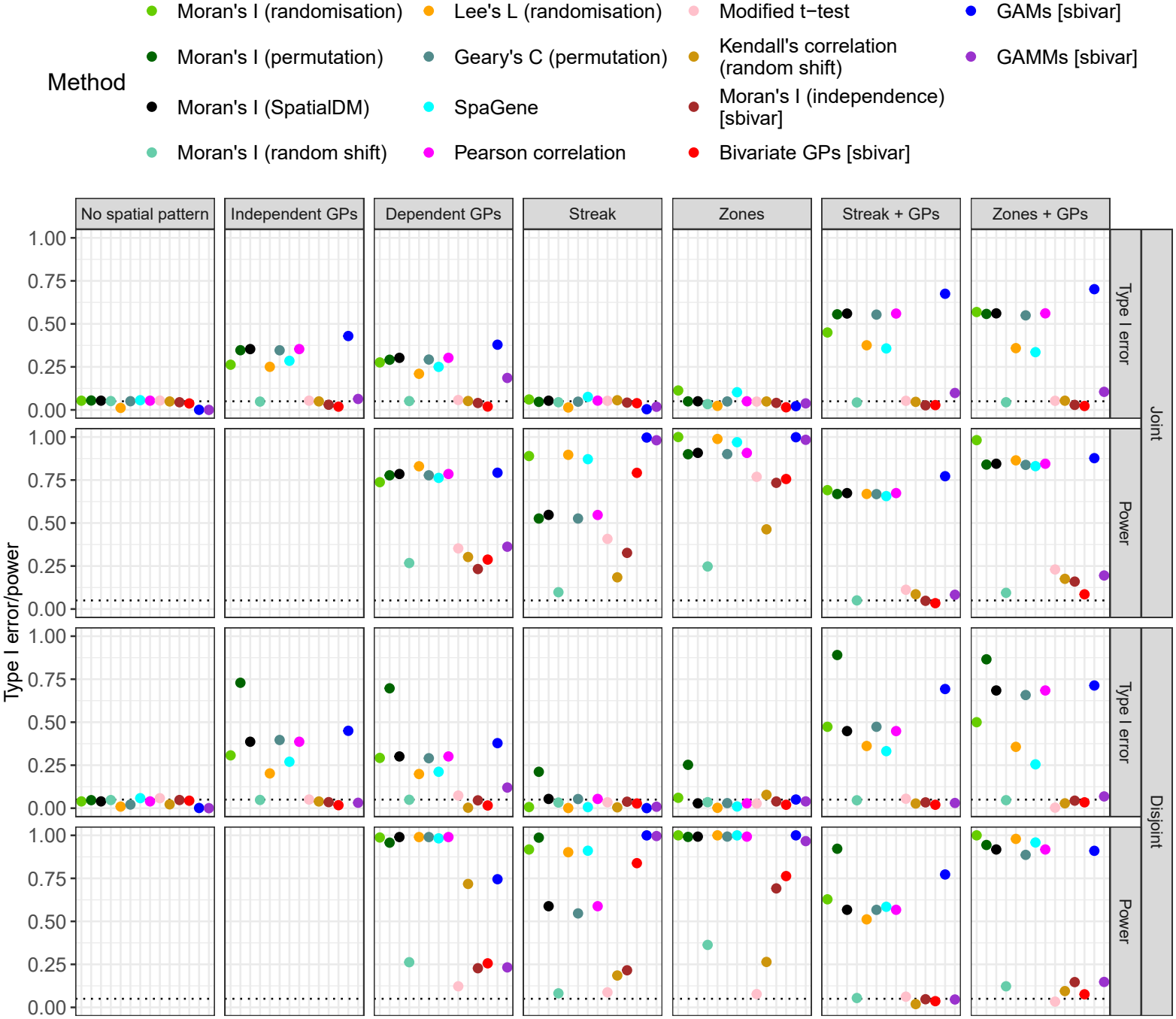
Type I error and power (y-axis, subrows) for different methods with null distributions in parentheses (colour, x-axis) for data with different spatial patterns (columns) with joint and disjoint coordinate sets (rows). The horizontal dotted line indicates the significance level of 5%, [sbivar] indicates methods introduced in this work.

Among the methods controlling type I error, the new methods in *sbivar*, bivariate Moran’s I (independence), bivariate GPs and GAMMs and the modified t-test have good power, whereas bivariate Moran’s I and Kendall’s correlation (random shift) have low power (Fig 3), likely because not all bivariate association is removed in the random shifts. Moreover, the power to detect shared zones or streaks is much lower in presence of background spatial autocorrelation. With disjoint coordinates, methods applicable to the original disjoint coordinates (bivariate Moran’s I (independence), bivariate GPs and GAMMs) mostly outperform methods relying on sample matching (modified t-test, Kendall’s correlation coefficient) in terms of power. In an additional simulation investigating the effect of anisotropy and nonstationarity in the SAC structure (see Fig S7 for setup), we found that the type I error is modestly inflated by anisotropy for the modified t-test, by nonstationarity for GAMs and GAMMs, and for the combination of anisotropy parameter *γ* = 0.5 and nonstationarity for bivariate GPs (Fig S20). Among methods controlling type I error, the power of Kendall’s correlation (random shift) and Moran’s I (independence) is mostly affected by anisotropy and nonstationarity (Fig S21). The calculation times of bivariate GPs and especially GAMMs are highest and increase quickly with the number of measurement spots, whereas GAMs, modified t-test and Moran’s I (independence) as implemented in *sbivar* are faster and their computation times increase more slowly with the number of spots (see Fig S22). GP fitting can be sped up considerably with GPU acceleration [20], but since most of the computation time is used in calculating the test statistic and its variance, such acceleration would not reduce the total computing time by much. The simulations with alignment errors reveal that the power of many methods decreases with increasing distortion of the alignment, but type I error control is upheld (Fig S23).

The results for multi-image analysis in Fig S24 reveal that GAMs, bivariate Moran’s I and Pearson correlation control the type I error under the null scenarios, as the increased estimator variability due to SAC is absorbed into the between-image variance in the linear model. Inverse weighting by the variance of 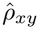for GAMs or *I*_*xy*_ for bivariate Moran’s I does not consistently improve the power and can even lead to FDR inflation, whereas scaling by the maximum value of *I*_*xy*_ does yield an increase in power for bivariate Moran’s I. Pearson correlation has low power overall. Without variance weighting, GAMs are slowest and Pearson correlation is the fastest method for multi-image analysis (Fig S25).

### Real data analysis

The original analysis of the Godfrey data (see Fig S26) was based on raw correlations between relative gene expressions and relative metabolite abundances, thresholded at an absolute value of 0.4. In comparison, our testing based approach discovers more feature pairs, the most significant pairs found by GAMs but not in the original analysis are shown in Fig 4 and Fig S27, all exhibiting negative association patterns. The original failure to detect these associations is likely due to high thresholding of the correlation coefficient, combined with Pearson correlation’s sensitivity to outliers from spots with low library sizes or total ion counts. Conversely, pairwise associations found in the original analysis but not by GAMs are shown in Figs S28-S29. The most significant findings of bivariate Moran’s I are shown in Fig S30. A striking result by GAMs in lung cancer sample LC 091, recurring to a lesser extent in breast cancer samples BC 515 Section 1 and BC 515 Section 2, is the positive association of a number of Kallikrein transcripts (*KLK10, KLK11, KLK12* and *KLK13* ) with PI 34:1 (Fig S31 and Table S1, the lack of significance in the other samples may be due to low expression of the Kallikrein genes). Kallikreins are a subgroup of serine proteases that participate in extracellular proteolysis, signaling cascades and remodeling of the tumor microenvironment [51]. Growing evidence suggests that many kallikreins are implicated in tumor inhibition [51–53], although it has been claimed that they promote breast cancer progression too [54]. *KLK10* and *KLK11* activate the PI3K/AKT signaling pathway [55, 56], whereas *KLK13* expression is reduced upon suppression of this pathway [57]. PI 34:1 is a member of the phosphatidylinositols, and serves as precursor for phosphoinositides that generate PIP_3_, the central lipid second messenger activating PI3K/AKT signaling [58]. Hence the shared involvement of the *KLK* transcripts and PI 34:1 in the PI3K/AKT signaling pathway may explain their co-occurrence in the tumor region of sample LC 091 (see Fig S32). The p-value distributions of replicated analyses per cancer type are shown in Fig S33, suggesting some feature pairs to have consistent association across replicates, although the sample size is too small to tell real from spurious findings, resulting in no significances after multiplicity correction.

**Fig 4.**
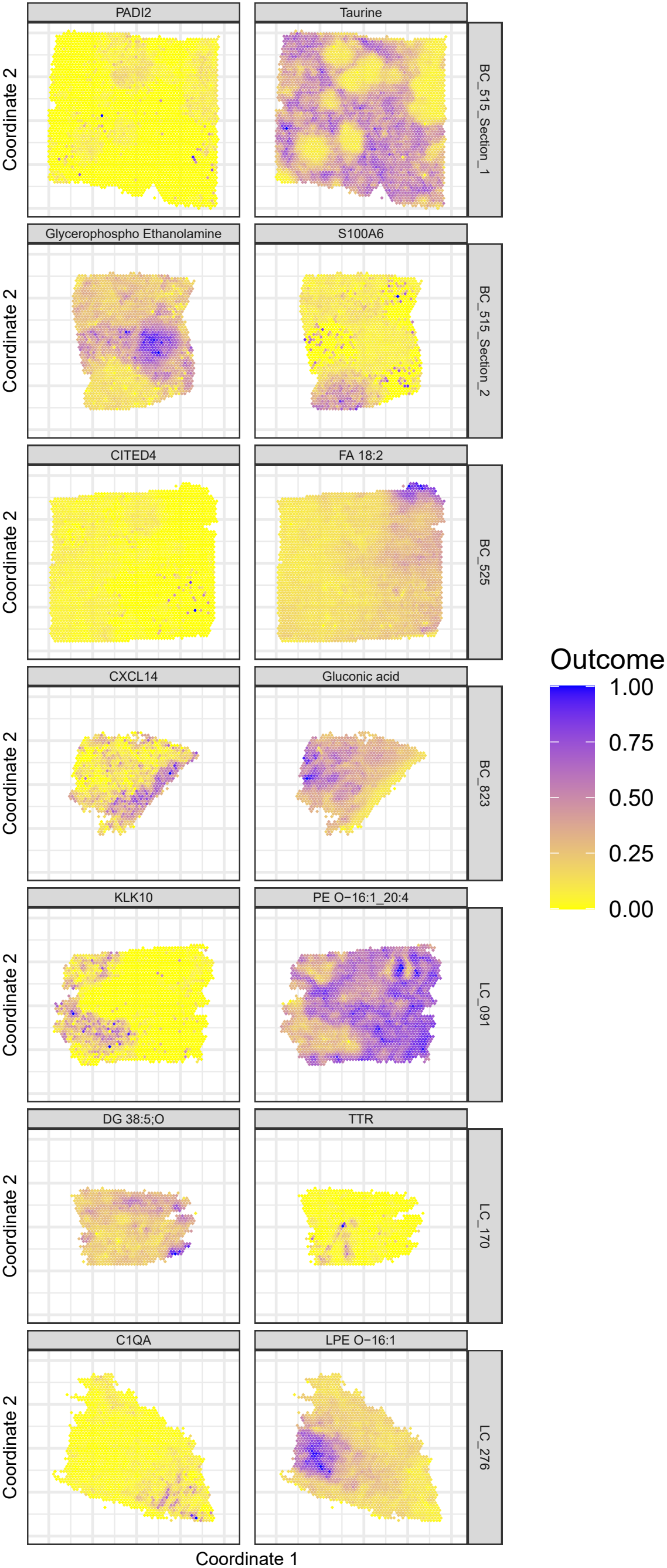
Gene-metabolite pairs most significant according to GAMs (all with negative correlations), but not found in the original analysis, in different sections of the Godfrey dataset (rows). The pairs in samples BC 515 Section 2, BC 823 and LC 170 were not found significant by bivariate Moran’s I. Relative abundances are shown in colour, scaled to the [0,1] range per image for legibility.

The authors of the Vicari data (Fig S35) focused mainly on the metabolite dopamine in their original analysis, finding it to be highly abundant on the side of the brain not lesioned by 6-OHDA, and tested for colocalisation of dopamine with genes after spot matching [9]. Some of those pairs were not confirmed by our analysis based on GAMs on the disjoint coordinate sets (Figs S36-S37). We find associations for many other pairs, the most significant positive ones per image according to GAMs are shown in Fig 5, the most significant negative ones in Fig S38, and results for Moran’s I (independence) in Fig S39. An interesting positive association found both by GAMs and bivariate Moran’s I in striatum samples V11L12-038 A1 and V11L12-038 B1 is the one between *CSRP1* transcripts and palmitoyl-L-carnitine (Fig S40). *CSRP1* belongs to the LIM-domain adaptor proteins, which interact with actin filaments to drive cellular shape changes [59]. Hence they are highly expressed in astrocytes [60, 61], which are known to rapidly restructure their cell shape [62]. Palmitoyl-L-carnitine is a long-chain acylcarnitine that serves as an intermediate to transport long-chain fatty acids into mitochondria for oxidation [63, 64]. Since lipids are known to be transferred from neurons to astrocytes for breakdown [64], palmitoyl-L-carnitine can be expected to be enriched in astrocytes and depleted in neurons, which explains its spatial association with *CSRP1* transcripts at the tissue level. A joint analysis of all replicates of the striata and substantia nigra found no significant pairs. Still, the p-value distributions do suggests some features to be associated (Fig S41), so insufficient sample sizes could explain the lack of significance after multiplicity correction.

**Fig 5.**
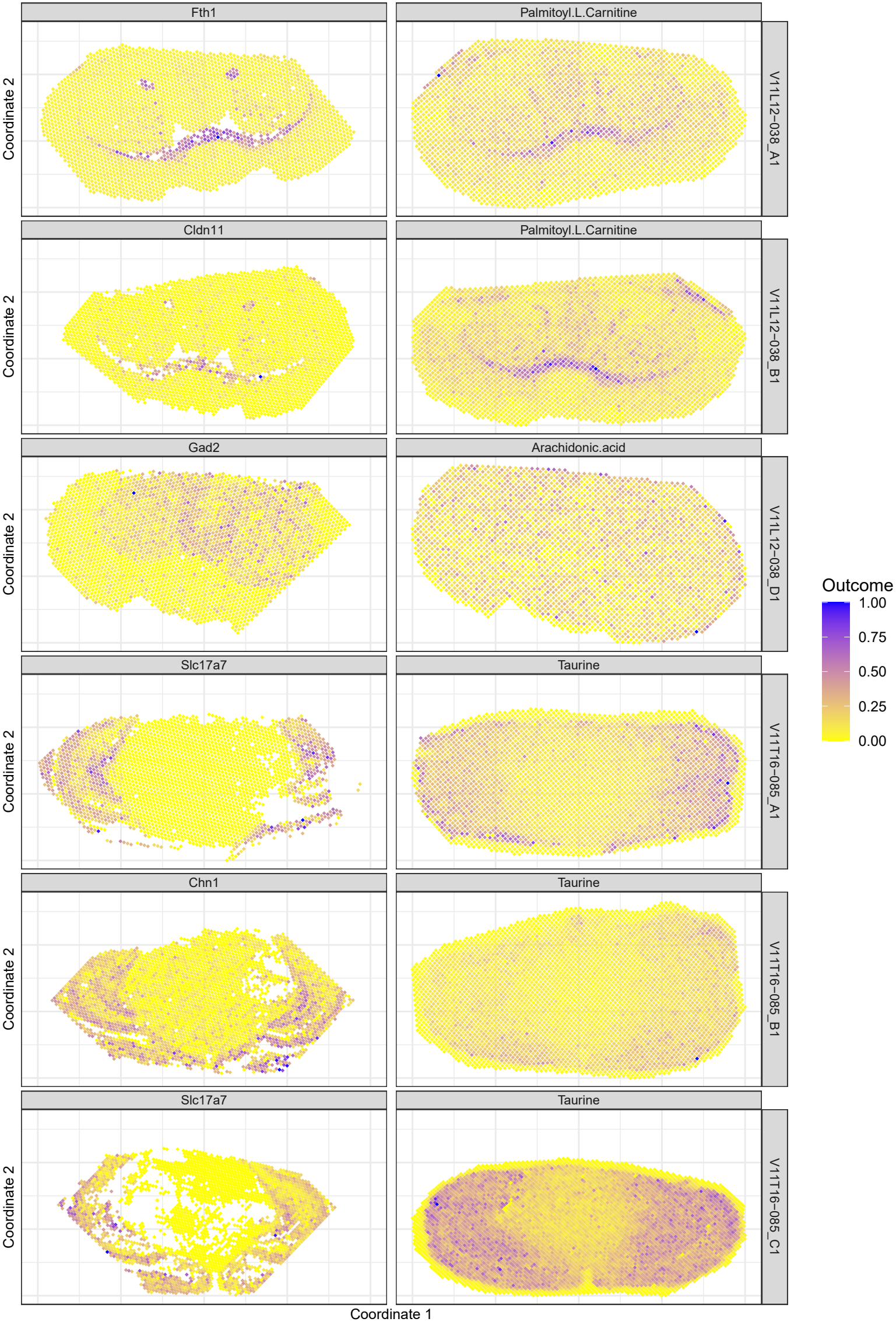
Gene-metabolite pairs most significantly positively correlated according to GAMs in different samples of the Vicari2024 dataset with significant positive correlations (rows). The first two pairs are also significant according to bivariate Moran’s I. Relative abundances are shown in colour, scaled to the [0,1] range per image for legibility.

## Discussion

It has long been known that statistical tests should account for dependence between observations, e.g. autocorrelation in spatial or temporal data, to achieve valid statistical inference. Yet most published tests for bivariate association in spatial omics data ignore this principle. Pearson correlation is a workhorse measure often applied to test for bivariate spatial association, but standard formulae for its variance are invalid in presence of spatial autocorrelation (SAC). For other association measures, such as bivariate Moran’s I, bivariate Geary’s C and Lee’s L, exact null distributions have been derived under the randomisation null hypothesis, which also implies absence of spatial autocorrelation in any variable. Also naive permutation procedures, which indiscriminately scramble observations over measurement locations, test this omnibus null hypothesis, falsely suggesting bivariate association where only univariate spatial patterns are being picked up. The reason is that permutation tests assume exchangeability of observations, which is not fulfilled in spatial omics data because of SAC.

Instead, bivariate association tests are needed that allow for SAC in both variables, while being sensitive only to true bivariate spatial association. The modified t-test accounts for the variance inflation of the Pearson correlation estimator under SAC, but relies on one-to-one spot matching and ignores neighbouring spots, which can lead to low power. Bivariate Moran’s I is an intuitive, nonparametric measure of spatial association that takes neighbourhood information into account and naturally generalizes to disjoint coordinate sets. Here we derive an analytical expression for its variance accounting for SAC, which controls type I error despite relying on a plug-in estimate of the SAC structure, even for moderate spot numbers of a few hundreds. Moreover, it achieves good power against a range of bivariate spatial patterns. Another method we describe relies on bivariate Gaussian processes (GPs), which model covariance between observations as a function of distance, and as such naturally allow for disjoint coordinate sets. This method is similar in spirit to generalised least squares (GLS) and autoregressive models, but avoids spot matching and arbitrarily assigning features to the role of outcome value or regressor, as required for the latter models. Yet bivariate GPs are mathematically and computationally demanding, scaling poorly with the number of spots despite past optimisation efforts. Moreover, they provide no quantitative measure of association strength, only a p-value. A third method we introduced uses generalized additive models (GAMs) to describe univariate spatial patterns using splines as a function of space. They overcome the disjointness by evaluating the spline surfaces of the two features tested in the same set of points to find their correlation, and calculate its variance by propagating the uncertainty on the spline coefficients. The resulting correlation estimate is easy to interpret and visualize. Yet as smoothing splines often fail to fully capture fine-grained spatial structure [65], GAMs can still become liberal in presence of short-range SAC because the variance of the spline coefficients is underestimated. As a solution, a GAMM allowing for residual SAC can be used for confirmatory purposes, albeit at a major computational cost and even then with poor FDR control due to slow convergence of the variance estimator. Hence all valid inference methods work in similar fashion: first univariate spatial structures are estimated, and only then can bivariate association tests be performed. Bivariate Moran’s I and GPs look for neighbourhood similarity, but the range of the spatial interaction is not estimated from the data, necessitating repeated testing at a prespecified set of ranges to achieve good power against all potential spatial patterns. GAMs, on the other hand, do not require a range specification as they look for a point-wise correspondence in spatial patterns, often yielding more human-visible patterns. Because of its nonparametric nature, bivariate Moran’s I struggles with heteroskedasticity as encountered in many omics data types, as is known for univariate Moran’s I [66, 67], which emphasises the need for adequate, custom normalization prior to such nonparametric tests. Bivariate GPs and GAMs, on the other hand, can cope with heteroskedasticity when augmented with appropriate outcome distributions. The three tests presented are amenable to other choices of variogram function, covariance kernel, normalization, spline type and outcome distribution, e.g. to account for anisotropy or nonstationarity, but are valid only as long as these these are appropriate for the data at hand.

As spatial omics technologies mature, replicated datasets consisting of multiple images of comparable tissue will become increasingly available. Since the principled approach of analysing all images through a single model may be computationally prohibitive, we propose to plug association measures calculated from single images into a linear model accounting for the experimental design as a pragmatic alternative. Since this quantifies variability across images and thus absorbs additional variability due to SAC, quantifying the variance of the image-wise estimates is unnecessary in this case. Yet it is important to render the estimates comparable before feeding them to the linear model, i.e. scaling bivariate Moran’s I to its maximum value, which depends on the measurement grid.

We demonstrated that many existing tests fail to control type I error in absence of bivariate spatial association. Yet comparing methods in term of power is harder as infinitely many types of bivariate spatial association exist, and additional, independent benchmarks are needed. Based on the results in this study, we recommend applying both bivariate Moran’s I and GAMs, inspect the results visually and if possible confirm findings using GAMMs (see Table S2 in S1 Text for an overview of the methods’ strengths and weaknesses). We have wrapped a suite of valid tests in the R-package *sbivar* available from github.com/sthawinke/sbivar.

## Supporting information

**S1 Text Supplementary** Method details, exhaustive simulation results and real data analysis.

**S1 Data Vicari alignment data** Landmarks for alignment of the Vicari data chosen using *MAGPIE* [13], and the coordinates of the resulting alignment.

## Acknowledgments

We thank Ilia Kats and Marco Vicari for useful discussions.

## References

1. Williams CG, Lee HJ, Asatsuma T, Vento-Tormo R, Haque A. An introduction to spatial transcriptomics for biomedical research. Genome Med. 2022;14(1):68. doi:10.1186/s13073-022-01075-1.

2. Du J, Yang YC, An ZJ, Zhang MH, Fu XH, Huang ZF, et al. Advances in spatial transcriptomics and related data analysis strategies. J Transl Med. 2023;21(1):330. doi:10.1186/s12967-023-04150-2.

3. Ravi VM, Will P, Kueckelhaus J, Sun N, Joseph K, Salié H, et al. Spatially resolved multi-omics deciphers bidirectional tumor-host interdependence in glioblastoma. Cancer Cell. 2022;40(6):639–655 e13. doi:10.1016/j.ccell.2022.05.009.

4. Mir CM, Pisterzi P, Poorter ID, Rilou M, van Kranenburg M, Heijs B, et al. Spatial multi-omics in whole skeletal muscle reveals complex tissue architecture. Communications Biology. 2024;7(1):1272. doi:10.1038/s42003-024-06949-1.

5. Long Y, Ang KS, Sethi R, Liao S, Heng Y, van Olst L, et al. Deciphering spatial domains from spatial multi-omics with SpatialGlue. Nat Methods. 2024:110. doi:10.1038/s41592-024-02316-4.

6. Zhang D, Deng Y, Kukanja P, Agirre E, Bartosovic M, Dong M, et al. Spatial epigenome–transcriptome co-profiling of mammalian tissues. Nature. 2023;616(7955):113–122. doi:10.1038/s41586-023-05795-1.

7. Ben-Chetrit N, Niu X, Swett AD, Sotelo J, Jiao MS, Stewart CM, et al. Integration of whole transcriptome spatial profiling with protein markers. Nat Biotechnol. 2023;41(6):788–793. doi:10.1038/s41587-022-01536-3.

8. Godfrey TM, Shanneik Y, Zhang W, Tran T, Verbeeck N, Patterson NH, et al. Integrating Ambient Ionization Mass Spectrometry Imaging and Spatial Transcriptomics on the Same Cancer Tissues to Identify RNA–Metabolite Correlations. Angew Chem Int Ed. 2025. doi:10.1002/anie.202502028.

9. Vicari M, Mirzazadeh R, Nilsson A, Shariatgorji R, Bjärterot P, Larsson L, et al. Spatial multimodal analysis of transcriptomes and metabolomes in tissues. Nat Biotechnol. 2024;42(7):1046–1050. doi:10.1038/s41587-023-01937-y.

10. Eng CHL, Lawson M, Zhu Q, Dries R, Koulena N, Takei Y, et al.Transcriptome-scale super-resolved imaging in tissues by RNA seqFISH+. Nature. 2019 4;568(7751):235–239. doi:10.1038/s41586-019-1049-y.

11. 10x Genomics. Visium Spatial Gene Expression;2020. Accessed 2026-07-01. Available from: https://www.10xgenomics.com/products/spatial-gene-expression.

12. Moore JL, Charkoftaki G. A Guide to MALDI Imaging Mass Spectrometry for Tissues. Journal of Proteome Research. 2023;22(11):3401–17. doi:10.1021/acs.jproteome.3c00167.

13. Williams EC, Franzén L, Lindvall MO, Hamm G, Oag S, Majumder MM, et al. Spatially resolved integrative analysis of transcriptomic and metabolomic changes in tissue injury studies. Nat Commun. 2026;17(1):205. doi:10.1038/s41467-025-68003-w.

14. Wartenberg D. Multivariate Spatial Correlation: A Method for Exploratory Geographical Analysis. Geographical Analysis. 1985;17(4):263–283. doi:10.1111/j.1538-4632.1985.tb00849.x.

15. Czaplewski RL. Expected value and variance of Moran’s bivariate spatial autocorrelation statistic for a permutation test. vol. 309. US Department of Agriculture, Forest Service, Rocky Mountain Forest and Range Experiment Station;1993.

16. Lee SI. Developing a bivariate spatial association measure: An integration of Pearson’s r and Moran’s I. Journal of Geographical Systems. 2001;3(4):369–385. doi:10.1007/s101090100064.

17. Miller BF, Bambah-Mukku D, Dulac C, Zhuang X, Fan J. Characterizing spatial gene expression heterogeneity in spatially resolved single-cell transcriptomic data with nonuniform cellular densities. Genome Res. 2021;31(10):1843–1855. doi:10.1101/gr.271288.120.

18. Liu Q, Hsu CY, Shyr Y. Scalable and model-free detection of spatial patterns and colocalization. Genome Res. 2022;32(9):1736–1745. doi:10.1101/gr.276851.122.

19. Sun S, Zhu J, Zhou X. Statistical analysis of spatial expression patterns for spatially resolved transcriptomic studies. Nat Methods. 2020 2;17(2):193–200. doi:10.1038/s41592-019-0701-7.

20. Kats I, Vento-Tormo R, Stegle O. SpatialDE2: Fast and localized variance component analysis of spatial transcriptomics. Biorxiv. 2021:2021–2010. doi:10.1101/2021.10.27.466045.

21. Yu S, Li WV. spVC for the detection and interpretation of spatial gene expression variation. Genome Biol. 2024;25(1):103. doi:10.1186/s13059-024-03245-3.

22. Zhu J, Sun S, Zhou X. SPARK-X: Non-parametric modeling enables scalable and robust detection of spatial expression patterns for large spatial transcriptomic studies. Genome Biol. 2021;22(1):184. doi:10.1186/s13059-021-02404-0.

23. DeTomaso D, Yosef N. Hotspot identifies informative gene modules across modalities of single-cell genomics. Cell Syst. 2021;12(5):446–456. doi:10.1016/j.cels.2021.04.005.

24. Li Z, Wang T, Liu P, Huang Y. SpatialDM for rapid identification of spatially co-expressed ligand-receptor and revealing cell-cell communication patterns. Nat Commun. 2023;14(1):3995. doi:10.1038/s41467-023-39608-w.

25. Dimitrov D, Schäfer PSL, Farr E, Rodriguez-Mier P, Lobentanzer S, Badia-I-Mompel P, et al. LIANA+ provides an all-in-one framework for cell-cell communication inference. Nat Cell Biol. 2024;26(9):1613–1622. doi:10.1038/s41556-024-01469-w.

26. Xu K, Cheng Q. Test of conditional independence in factor models via Hilbert–Schmidt independence criterion. J Multivar Anal. 2024;199:105241. doi:10.1016/j.jmva.2023.105241.

27. Clifford P, Richardson S, Hemon D. Assessing the Significance of the Correlation between Two Spatial Processes. Biometrics. 1989;45(1):123–134. doi:10.2307/2532039.

28. Dutilleul P, Clifford P, Richardson S, Hemon D. Modifying the t test for assessing the correlation between two spatial processes. Biometrics. 1993:305–314. doi:10.2307/2532625.

29. Dutilleul P, Pelletier B, Alpargu G. Modified F tests for assessing the multiple correlation between one spatial process and several others. J Stat Plan Inference. 2008;138(5):1402–1415. doi:10.1016/j.jspi.2007.06.022.

30. Mrkvička T, Dvořák J, González JA, Mateu J. Revisiting the random shift approach for testing in spatial statistics. Spat Stat. 2021;42:100430. Towards Spatial Data Science. doi:10.1016/j.spasta.2020.100430.

31. Ridder GI, Hardy OJ, Ovaskainen O. Generating spatially realistic environmental null models with the shift-and-rotate approach helps evaluate false positives in species distribution modelling. Methods in Ecology and Evolution. 2024;15(12):2331–2342. doi:10.1111/2041-210X.14443.

32. Viladomat J, Mazumder R, McInturff A, McCauley DJ, Hastie T. Assessing the significance of global and local correlations under spatial autocorrelation: A nonparametric approach. Biometrics. 2014;70(2):409–418. doi:10.1111/biom.12139.

33. Seal S, Neelon B. SpaceBF: Spatial coexpression analysis using Bayesian Fused approaches in spatial omics datasets. Gigascience. 2026;15. doi:10.1093/gigascience/giag006.

34. Hu J, Wang SG, Hou Y, Chen Z, Liu L, Li R, et al. Multi-omic profiling of clear cell renal cell carcinoma identifies metabolic reprogramming associated with disease progression. Nat Genet. 2024;56(3):442–457. doi:10.1038/s41588-024-01662-5.

35. Matheron G. Principles of geostatistics. Economic Geology. 1963;58(8):1246–1266. doi:10.2113/gsecongeo.58.8.1246.

36. Pebesma EJ. Multivariable geostatistics in S: The gstat package. Computers & Geosciences. 2004;30(7):683–691. doi:10.1016/j.cageo.2004.03.012.

37. Kelejian HH, Prucha IR. On the asymptotic distribution of the Moran I test statistic with applications. J Econom. 2001;104(2):219–257. doi:10.1016/S0304-4076(01)00064-1.

38. Liu Y, Xie J. Cauchy combination test: A powerful test with analytic p-value calculation under arbitrary dependency structures. J Am Stat Assoc. 2020;115(529):393–402. doi:10.1080/01621459.2018.1554485.

39. Zhang D, Lin X. Hypothesis testing in semiparametric additive mixed models. Biostatistics. 2003;4(1):57–74. doi:10.1093/biostatistics/4.1.57.

40. Self SG, Liang KY. Asymptotic Properties of Maximum Likelihood Estimators and Likelihood Ratio Tests Under Nonstandard Conditions. J Am Stat Assoc. 1987;82(398):605–610. doi:10.2307/2289471.

41. Wood SN. Fast stable restricted maximum likelihood and marginal likelihood estimation of semiparametric generalized linear models. Journal of the Royal Statistical Society (B). 2011;73(1):3–36. doi:10.1111/j.1467-9868.2010.00749.x.

42. Gauss CF. Theoria Combinationis Observationum Erroribus Minimis Obnoxia. Göttingen: Dieterich. 1823.

43. Canete NP, Iyengar SS, Ormerod JT, Baharlou H, Harman AN, Patrick E. spicyR: Spatial analysis of in situ cytometry data in R. Bioinformatics. 2022;38(11):3099–3105. doi:10.1093/bioinformatics/btac268.

44. Hawinkel S, Yang X, Poelmans W, Motte H, Beeckman T, Maere S. Smoppix: Unified nonparametric analysis of single-molecule spatial omics data using probabilistic indices. Genome Biol. 2026. doi:10.1186/s13059-026-03976-5.

45. Cliff AD, Ord JK. Spatial processes: Models & applications. vol. 44. Pion London;1981.

46. Benjamini Y, Hochberg Y. Controlling the False Discovery Rate: A Practical and Powerful Approach to Multiple Testing. JRSS Series B. 1995 1;57(1):289–300. doi:10.1111/j.2517-6161.1995.tb02031.x.

47. Jefferis G, Kemp SE, Arya S, Mount D. RANN: Fast Nearest Neighbour Search (Wraps ANN Library) Using L2 Metric; 2024. R package version 2.6.2. doi:10.32614/CRAN.package.RANN.

48. Murphy KM, Topel RH. Estimation and Inference in Two-Step Econometric Models. J Bus Econ Stat. 1985;3(4):370–379. doi:10.1198/073500102753410417.

49. Beckmann CF, Jenkinson M, Smith SM. General multilevel linear modeling for group analysis in FMRI. Neuroimage. 2003;20(2):1052–1063. doi:10.1016/S1053-8119(03)00435-X.

50. Lennon JJ. Red-Shifts and Red Herrings in Geographical Ecology. Ecography. 2000;23(1):101–113. doi:10.1111/j.1600-0587.2000.tb00265.x.

51. He X, Meng F, Qin L, Liu Z, Zhu X, Yu Z, et al. KLK11 suppresses cellular proliferation via inhibition of Wnt/β-catenin signaling pathway in esophageal squamous cell carcinoma. American Journal of Cancer Research. 2019;9(10):2264.

52. Zhao R, Wang S, Liu J, Xu C, Zhang S, Shao Y, et al. KLK11 acts as a tumor-inhibitor in laryngeal squamous cell carcinoma through the inactivation of Akt/Wnt/β-catenin signaling. J Bioenerg Biomembr. 2021;53(1):85–96. doi:10.1007/s10863-020-09870-4.

53. Ding Y, Wang Z, Chen C, Li D, Wang W, Jia Y, et al. miR-1304 targets KLK11 to regulate gastric cancer cell proliferation through the mTOR signaling pathway. Carcinogenesis. 2024;45(1 -2):45–56. doi:10.1093/carcin/bgad077.

54. Sano A, Sangai T, Maeda H, Nakamura M, Hasebe T, Ochiai A. Kallikrein 11 expressed in human breast cancer cells releases insulin-like growth factor through degradation of IGFBP-3. Int J Oncol. 2007. doi:10.3892/ijo.30.6.1493.

55. Wei H, Dong C, Shen Z. Kallikrein-related peptidase (KLK10) cessation blunts colorectal cancer cell growth and glucose metabolism by regulating the PI3K/Akt/mTOR pathway. Neoplasma. 2020;67(4):889–897. doi:10.4149/neo2020190814n758.

56. Zhang Y, Xu Z, Sun Y, Chi P, Lu X. Knockdown of KLK11 reverses oxaliplatin resistance by inhibiting proliferation and activating apoptosis via suppressing the PI3K/AKT signal pathway in colorectal cancer cell. OncoTargets and therapy. 2018;11:809–821. doi:10.2147/OTT.S151867.

57. Paliouras M, Diamandis EP. Intracellular signaling pathways regulate hormone-dependent kallikrein gene expression. Tumour Biol. 2008;29(2):63–75. doi:10.1159/000135686.

58. Liu R, Chen Y, Liu G, Li C, Song Y, Cao Z, et al. PI3K/AKT pathway as a key link modulates the multidrug resistance of cancers. Cell Death Dis. 2020;11(9):797. doi:10.1038/s41419-020-02998-6.

59. Weiskirchen R, Günther K. The CRP/MLP/TLP family of LIM domain proteins: Acting by connecting. Bioessays. 2003;25(2):152–162. doi:10.1002/bies.10226.

60. Uhlén M, Fagerberg L, Hallström BM, Lindskog C, Oksvold P, Mardinoglu A, et al. Proteomics. Tissue-based map of the human proteome. Science (New York, NY). 2015;347(6220):1260419. doi:10.1126/science.1260419.

61. Zeisel A, Muñoz-Manchado AB, Codeluppi S, Lönnerberg P, Manno GL, Juréus A, et al. Cell types in the mouse cortex and hippocampus revealed by single-cell RNA-seq. Science. 2015;347(6226):1138–1142. doi:10.1126/science.aaa1934.

62. Bernardinelli Y, Muller D, Nikonenko I. Astrocyte-synapse structural plasticity. Neural Plast. 2014;2014(1):232105. doi:10.1155/2014/232105.

63. Schönfeld P, Reiser G. How the brain fights fatty acids’ toxicity. Neurochem Int. 2021;148:105050. doi:10.1016/j.neuint.2021.105050.

64. Ioannou MS, Jackson J, Sheu SH, Chang CL, Weigel AV, Liu H, et al. Neuron-astrocyte metabolic coupling protects against activity-induced fatty acid toxicity. Cell. 2019;177(6):1522–1535. doi:10.1016/j.cell.2019.04.001.

65. Hawinkel S, De Meyer S, Maere S. Spatial Regression Models for Field Trials: A Comparative Study and New Ideas. Frontiers in Plant Science. 2022;13. doi:10.3389/fpls.2022.858711.

66. Waldhör T. The spatial autocorrelation coefficient Moran’s I under heteroscedasticity. Stat Med. 1996;15(7 -9):887–892. doi:10.1002/(sici)1097-0258(19960415)15:7/9¡887::aid-sim257¿3.0.co;2-e.

67. Zhang T, Lin G. On Moran’s I coefficient under heterogeneity. Comput Stat Data Anal. 2016;95:83–94. doi:10.1016/j.csda.2015.09.010.

