## Supplementary Material for "Statistical tests for bivariate spatial association across multi-omics data with disjoint coordinates"

**4 Recommendations** **48**

**5 Software versions** **50**

### 1 Algorithms

#### 1.1 Existing algorithms

##### 1.1.1 Bivariate Moran’s I

Bivariate Moran’s I is defined for two vectors $\mathbf{x}$ and $\mathbf{y}$ as $I_{xy} = \sum_{i=1}^{n}\sum_{j=1}^{m} w_{ij}x_i y_j = \mathbf{x}^t\mathbf{W}\mathbf{y}$. When testing only for spatial association between spots, under the randomisation null hypothesis for joint coordinate sets with $n$ spots, $I_{xy}$ has an expected value of $E(I_{xy}) = \frac{\text{Cor}(x,y)}{(n-1)}$ [1]. If zero correlation within the same spots is also considered part of the null hypothesis, as we do throughout the article, than the expectation equals 0. Several versions of the variance of bivariate Moran’s I exist for the case of joint coordinate sets. Czaplewski [1] provides the variance under the randomisation null hypothesis. Li et al. [2] provides a variance assuming no SAC in either outcome variable in their *SpatialDM* method, as do DeTomaso and Yosef [3]. Miller et al. [4] employ bivariate Moran's I in their *MERINGUE* method, using permutations to determine significance, as does Wartenberg [5].

###### 1.1.2 The modified t-test

Pearson correlation after spot matching or aggregation is commonly used to test for spatial association in multi-omics studies [6, 7, 8], although it is known that such test must account for SAC to be valid [9]. The most common correction is the modified t-test, correcting the degrees of freedom of the t-test and thus the variance estimate of the Pearson correlation estimate $\hat{\rho}_{xy}$ based on SAC in the two variables [10, 11, 12]. The variance of $\hat{\rho}_{xy}$ is given by $(n-1)^{-2}tr(\mathbf{\Sigma}_x\mathbf{\Sigma}_y)$, with $\mathbf{\Sigma}_x$ and $\mathbf{\Sigma}_y$ the spatial autocorrelation matrices of $x$ and $y$, respectively, and $tr(.)$ the trace function [12, 10].

###### 1.1.3 Other measures of spatial association

SpaGene by [13] defines its own measure of spatial colocalisation based on k-nearest neighbour networks, and determines significance through permutations. Seal and Neelon [14] notice problems with autocorrelation in simulations, and present their method SpaceBF to account for SAC in a Bayesian framework. Yet we found this method too slow to be applied in our simulations or on real spatial omics data. Lee [15] presents Lee's L as measure of spatial association for joint coordinate sets. Bivand and Wong [16] generalised Geary's C for the bivariate case of joint coordinate sets. To determine significance for any test statistic while accounting for SAC, custom permutation schemes have been developed that test the independence null hypothesis: the so-called "random-shift" tests [17, 18]. These involve shifting both images with respect to each other, which retains SAC, calculating the association measure based on the regions that still overlap, and correcting their variance for the reduction in overlap area (see Section 1.2.1.6 for the application to bivariate Moran's I).

#### 1.2 New algorithms

###### 1.2.1 Bivariate Moran's I under the independence null hypothesis

**1.2.1.1 Expectation** Under the independence null hypothesis, implying no correlation within the same measurement spots, for both joint and disjoint coordinate sets, bivariate Moran's I has expectation zero:

$$\begin{aligned} E(I_{xy}) &= E\left(\sum_{i=1}^{n}\sum_{j=1}^{m} w_{ij}x_iy_j\right) \\ &= \sum_{i=1}^{n}\sum_{j=1}^{m} w_{ij}E\left(x_iy_j\right) = 0 \end{aligned} \tag{i}$$

**1.2.1.2 Maximum absolute value** The lower and upper bounds of bivariate Moran's I statistic are dependent on the weight matrix and usually not -1 and 1. Instead, they depend on the largest singular value of the weight matrix as follows. The problem is to find $\mathbf{x}$ and $\mathbf{y}$ that maximize $I_{xy} = \mathbf{x}^t\mathbf{W}\mathbf{y}$ for fixed $\mathbf{W}$ and subject to $\mathbf{x}^t\mathbf{x} = n-1$ and $\mathbf{y}^t\mathbf{y} = m-1$. Given the singular value decomposition $\mathbf{U}\Upsilon\mathbf{V}^t$ of $\mathbf{W}$, with $r = \min(n, m)$ and $\Upsilon$ the diagonal matrix of singular values, this amounts to optimizing $\mathbf{x}^t\mathbf{W}\mathbf{y} = \mathbf{x}^t\mathbf{U}\Upsilon\mathbf{V}^t\mathbf{y}$. The columns of $\mathbf{U}$ and $\mathbf{V}$ being orthonormal, and $\upsilon_1 \geq \upsilon_2 \geq \ldots \geq \upsilon_r$, this measure is optimized when $\mathbf{x}$ aligns perfectly with the first column of $\mathbf{U}$ ($\mathbf{u}_1$), and $\mathbf{y}$ with the first column of $\mathbf{V}$ ($\mathbf{v}_1$), achieving $\sqrt{(n-1)(m-1)}\upsilon_1$ as maximum value. By symmetry, the minimum is achieved when $\mathbf{x} = -\mathbf{u}_1$ and $\mathbf{y} = \mathbf{v}_1$ (or $\mathbf{x} = \mathbf{u}_1$ and $\mathbf{y} = -\mathbf{v}_1$), which is $-\sqrt{(n-1)(m-1)}\upsilon_1$. It may be useful to scale $I_{xy}$ values by their maximum absolute value $\sqrt{(n-1)(m-1)}\upsilon_1$ to render them comparable across sections with different measurements grids and corresponding weight matrices.

**1.2.1.3 Variance** The expectation and maximum absolute value are the same under the randomisation and independence null hypotheses, but the variance differs as it is affected by SAC, as derived in Eq. 2 in the main paper. We estimate the SAC matrices nonparametrically using Matheron's variogram estimator [19] following

Dutilleul et al. [12]. Next we fit linear and exponential functions through the estimate variogram using the *fit.variogram* function in the *gstat* R-package [20], fixing the sill to 1, and retain the fits ($\hat{\sigma}_x$ and $\hat{\sigma}_y$) with the smallest mean squared error loss. Then we plug the covariance as a function of distance $\hat{\sigma}_x(d_{ik})$ and $\hat{\sigma}_y(d_{jl})$ into Eq. 2 in the main paper, to yield $\widehat{\mathrm{Var}}(I_{xy}) = \sum_{i=1}^{n}\sum_{j=1}^{m}\sum_{k=1}^{n}\sum_{l=1}^{m} w_{ij}w_{kl}\hat{\sigma}_x(d_{ik})\hat{\sigma}_y(d_{jl}) = tr(\mathbf{W}^t\hat{\mathbf{\Sigma}}_x\mathbf{W}\hat{\mathbf{\Sigma}}_y)$. The formula for joint coordinate sets is the same; we do not distinguish between association within the same coordinates and between nearby coordinates, the former ones simply get the highest weight.

**1.2.1.4 Empirical distribution** For significance testing on $I_{xy}$ we the assume that $\frac{I_{xy}}{\sqrt{\widehat{\mathrm{Var}}(I_{xy})}}$ is asymptotically standard normal. This is justified by theoretical results on univariate Moran's I [21], but we also verify it empirically in a small simulation study in the 'none' scenario from the parametric simulations with disjoint coordinate sets, revealing approximate normality of the statistic (Fig S1).

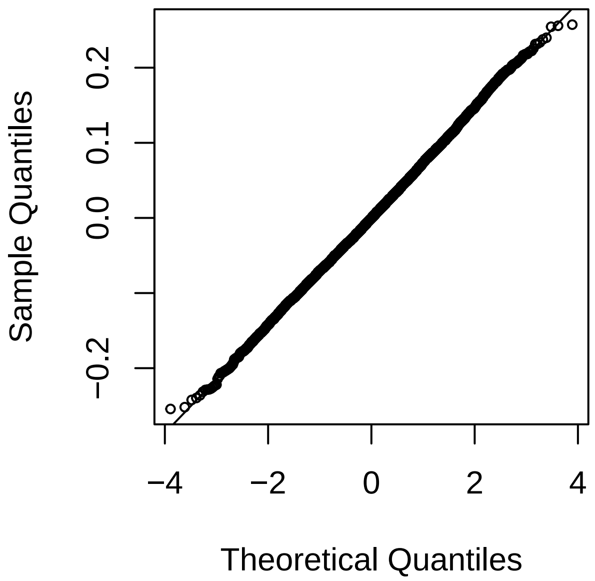

Fig S1: Normal QQ-plot of 10,000 bivariate Moran's I statistics generated under the 'none' scenario with $\eta = 0.0001$.

**1.2.1.5 The weight function** The choice of weight matrix $\mathbf{W}$ determines the power of tests based on bivariate Moran's I. The three different weight functions tested by default, for coordinate matrices shifted and scaled to the unit square with corners $(0,0)^t$, $(0,1)^t$, $(1,1)^t$ and $(1,0)^t$, are illustrated in Fig S2. The p-values from the different weight matrices are combined into one p-value through the Cauchy combination test [13].

**1.2.1.6 The random-shift null distribution** As an alternative to our analytical derivation of the variance and normality assumption, we adapt the random shift tests by Mrkvička et al. [17] and Ridder et al. [18] to bivariate Moran's I, and approximate its null distribution under the independence null hypothesis as follows. First the coordinate matrices are moved and scaled to the unit square. The weight matrix and observed bivariate Moran's I are calculated as before. Next a point is drawn uniformly on a disc of radius 0.5 centered in the origin, defining a vector $\mathbf{v}$ from the origin to this point. Define a shifted window with corners $(0,0)^t + \mathbf{v}$, $(0,1)^t + \mathbf{v}$, $(1,1)^t + \mathbf{v}$ and $(1,0)^t + \mathbf{v}$, and find the subsets $\mathbf{C}_-$ and $\mathbf{E}_-$ of sizes $n_-$ and $m_-$ within the shifted window. Shift $\mathbf{C}_-$ in the reverse direction to obtain $\mathbf{C}^*_- = \mathbf{C}_- - \mathbf{v}$. Calculate bivariate Moran's I

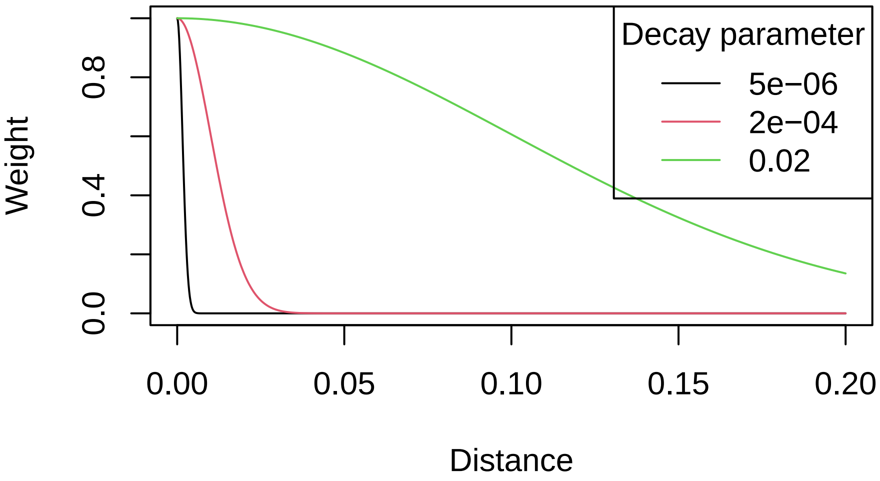

Fig S2: Illustration of weighting functions used for constructing the weight matrix used for a number of association metrics, including bivariate Moran's I.

$I_{xy-}$ with coordinates $\mathbf{C}^*_-$ and $\mathbf{E}_-$ and corresponding observations, and scale it to $I_{xy-}\sqrt{n_- m_-}$ to account for the variance inflation due to subsetting. Repeat this procedure $B$ times by drawing more instances $\mathbf{v}$ on the disc to build a null distribution for the scaled bivariate Moran's I statistic $I_{xy}\sqrt{nm}$. Following Phipson and Smyth [22], then calculate $Q = \frac{1+\sum_{b=1}^{B} I\left(I_{xy}\sqrt{nm} > I^b_{xy-}\sqrt{n^b_- m^b_-}\right)}{B+1}$ with $I(.)$ the indicator function, and the p-values as $2 \times \min(Q, 1-Q)$.

##### 1.2.2 Bivariate Gaussian processes (GPs)

**1.2.2.1 Derivation of the score test** The GP score test statistic $U$ was defined in Eq. 5 in the main text. It tests the null hypothesis that $\tau$, the variance of the random effect engendering the bivariate association, equals 0. Under the null, $U$ asymptotically follows a mixture of chi-squared distributions, which can be approximated using the Satterwaithe method by a scaled chi-squared distribution $\kappa\chi^2_\nu$ [23, 24]. Call $e = 0.5tr\left(\mathbf{P}^{-1}\mathbf{\Sigma}^{biv}\right)$ and the Fisher information matrix $I_{\tau\tau} = 0.5tr\left((\mathbf{P}^{-1}\mathbf{\Sigma}^{biv})^2\right)$ with $\mathbf{A}$ the design matrix, $\mathbf{P} = \mathbf{\Sigma}_*^{-1} - \mathbf{\Sigma}_*^{-1}\mathbf{A}(\mathbf{A}^t\mathbf{\Sigma}_*^{-1}\mathbf{A})^{-1}\mathbf{A}^t\mathbf{\Sigma}_*^{-1}$, then $\kappa = I_{\tau\tau}/(2e)$ and $\nu = 2e^2/I_{\tau\tau}$ [24, 23]. Zhang et al. [23] recommend to use the efficient information $\tilde{I}_{\tau\tau} = I_{\tau\tau} - I_{\tau\boldsymbol{\theta}}I^{-1}_{\boldsymbol{\theta\theta}}I^t_{\tau\boldsymbol{\theta}}$ rather than $I_{\tau\tau}$ with $\boldsymbol{\theta}$ the vector of (co)variance parameters of length $g$ defining $\mathbf{\Sigma}_*$. For the typical geostatistics parametrisation, used for instance in the *gls* function in the *nlme* R-package [25] this is $\boldsymbol{\theta}^t = (\sigma^2_x, \sigma^2_y, \pi_x, \pi_y, l_x, l_y)$ with $\sigma^2_x$ the variance, $\pi_x$ the nugget and $l_x$ the length scale of the x-variable, and analogously for the y-variable. Here we use the Gaussian covariance function $\text{Cov}(x_i, x_j) = \sigma^2_x\left(\pi_x I(d_{ij} = 0) + (1-\pi_x)\exp\left[-(\frac{d_{ij}}{l_x})^2\right]\right)$ with $d_{ij}$ the distance between observations $i$ and $j$. Further $I_{\tau\boldsymbol{\theta}} = 0.5tr(\mathbf{\Sigma}_*^{-1}\mathbf{\Sigma}^{biv}\mathbf{\Sigma}_*^{-1}\frac{\partial\mathbf{\Sigma}_*}{\partial\boldsymbol{\theta}})$ and $I_{\boldsymbol{\theta\theta}} = 0.5tr(\mathbf{\Sigma}_*^{-1}\frac{\partial\mathbf{\Sigma}_*}{\partial\boldsymbol{\theta}}\mathbf{\Sigma}_*^{-1}\frac{\partial\mathbf{\Sigma}_*}{\partial\boldsymbol{\theta}})$ [23]. Unlike in the simple univariate case in e.g. *SpatialDE2* by Kats et al. [24] where $\boldsymbol{\theta} = \sigma^2_x$, here $\frac{\partial\mathbf{\Sigma}_*}{\partial\boldsymbol{\theta}}$ is a $h \times h \times g$ tensor. The traces are here taken over the first two dimensions only, meaning the trace is applied over matrix slices of dimensions $h \times h$, such that $I_{\boldsymbol{\theta\theta}}$ is a $g \times g$ matrix and $I_{\tau\boldsymbol{\theta}}$ a vector of length $g$. Also the matrix multiplications $\mathbf{\Sigma}_*^{-1}\frac{\partial\mathbf{\Sigma}_*}{\partial\boldsymbol{\theta}}$ are done one matrix at a time to yield tensors again. We now find the derivatives $\frac{\partial\mathbf{\Sigma}_*}{\partial\boldsymbol{\theta}}$ for on-diagonal elements as

$$
\begin{aligned}
\frac{\partial \Sigma_{*ii}}{\partial \sigma_x} &= 2\sigma_x \\
\frac{\partial \Sigma_{*ii}}{\partial \pi_x} &= 0 \\
\frac{\partial \Sigma_{*ii}}{\partial l_x} &= 0,
\end{aligned} \tag{ii}
$$

and for off-diagonal elements

$$
\begin{aligned}
\frac{\partial \Sigma_{*ij}}{\partial \sigma_x} &= 2\sigma_x \exp\left[-(d_{ij}/l_x)^2\right](1-\pi_x) \\
\frac{\partial \Sigma_{*ij}}{\partial \pi_x} &= -\sigma_x^2 \exp\left[-(d_{ij}/l_x)^2\right] \\
\frac{\partial \Sigma_{*ij}}{\partial l_x} &= 2\sigma_x^2(1-\pi_x)\exp\left[-(d_{ij}/l_x)^2\right]\frac{d_{ij}^2}{l_x^3}.
\end{aligned} \tag{iii}
$$

Switching covariance kernels implies revisiting these expressions. The test is designed for univariate GPs estimated by restricted maximum likelihood (REML), e.g. with the *gls* function in the *nlme* package [25], which separates mean from covariance estimation. For speedup, one could instead use the *gpytorch* python package to fit the univariate GPs, on CPU as well as with GPU acceleration [26]. It is then crucial not to estimate the mean freely as default in *gpytorch*, but instead fix it to the empirical mean to approximate the REML outcome. Yet for the final computation time the GPU acceleration is of little importance, as calculating the test statistic $U$ and its variance takes up most of the computation time. Hence GPU acceleration is not included in the *sbivar* package.

**1.2.2.2 The score test for other outcome distributions** The score test can be generalized to other outcome distributions and link functions, although this is currently not implemented in the *sbivar* package. Building on Zhang et al. [23] and Kats et al. [24], Eq. 5 then becomes

$$
U = 1/2\left((\mathbf{z}^* - \boldsymbol{\mu})^t \boldsymbol{\Delta}\mathbf{K}^{-1}\boldsymbol{\Sigma}^{biv}\mathbf{K}^{-1}\boldsymbol{\Delta}(\mathbf{z}^* - \boldsymbol{\mu})\right). \tag{iv}
$$

For count distributions where the variance equals the mean, this becomes $\boldsymbol{\Delta} = \text{diag}(g'(\boldsymbol{\mu})) = \text{diag}(\boldsymbol{\mu})^{-1}$ and $\mathbf{K} = g'(\boldsymbol{\mu})\boldsymbol{\Sigma}^* g'(\boldsymbol{\mu})^t = \text{diag}(\boldsymbol{\mu})^{-1}$ and $g$ the link function. $\mathbf{z}^* = \frac{z-\boldsymbol{\mu}}{\boldsymbol{\mu}}$ is the working vector under the null.

##### 1.2.3 Generalized additive models (GAMs)

**1.2.3.1 Other outcome distributions on a fixed grid** We extend the bivariate test based on GAMs proposed in the main text to other outcome distributions and link functions. The models for the mean can then be written with the link functions $g$ and $h$ as

$$
\begin{aligned}
g(E(x_i)) &= \beta_{0x} + s_x(\mathbf{c}_i) \\
h(E(y_j)) &= \beta_{0y} + s_y(\mathbf{e}_j).
\end{aligned} \tag{v}
$$

Call $\iota_i = s_x(\mathbf{c}_i)$ and $\zeta_j = s_y(\mathbf{e}_j)$. The covariance estimate is then $q_{xy} = (B-1)^{-1}\sum_{b=1}^{B}(\iota_b - \bar{\iota})(\zeta_b - \bar{\zeta})$, with gradient $\nabla q_{xy}^t = \left(\frac{\partial q_{xy}}{\partial \iota_1}, \frac{\partial q_{xy}}{\partial \iota_2}, \ldots, \frac{\partial q_{xy}}{\partial \iota_B}, \frac{\partial q_{xy}}{\partial \zeta_1}, \frac{\partial q_{xy}}{\partial \zeta_2}, \ldots, \frac{\partial q_{xy}}{\partial \zeta_B}\right)$. The variance-covariance matrix of $\boldsymbol{\iota}$ is given by $\nabla\hat{\boldsymbol{\iota}}^t \boldsymbol{\Phi}_x \nabla\hat{\boldsymbol{\iota}}$ with $\nabla\hat{\boldsymbol{\iota}}^t = \left(\frac{\partial\hat{\boldsymbol{\iota}}}{\partial\beta_1}, \frac{\partial\hat{\boldsymbol{\iota}}}{\partial\beta_2}, \ldots, \frac{\partial\hat{\boldsymbol{\iota}}}{\partial\beta_h}\right)$, in which $\frac{\partial\hat{\boldsymbol{\iota}}}{\partial\beta_1} = (g^{-1})'(\beta_0 + \mathbf{H}_x\boldsymbol{\beta}_x + g_x(\mathbf{c}_i))\mathbf{H}_{1x}$ and $\boldsymbol{\Phi}_x$ the variance-covariance matrix of the thin plate spline coefficients $\boldsymbol{\beta}_x$. The variance of the correlation coefficient can be approximated by

$$
\text{Var}(\hat{q}_{xy}) \approx \nabla q_{xy}^t \nabla\hat{\mathbf{z}}^t \boldsymbol{\Phi}\nabla\hat{\mathbf{z}}\nabla q_{xy}, \tag{vi}
$$

with $\mathbf{\Phi}$ a block diagonal matrix of $\mathbf{\Phi}_x$ and $\mathbf{\Phi}_y$, and $\hat{\mathbf{z}}^t = (\hat{\boldsymbol{\iota}}^t, \hat{\boldsymbol{\zeta}}^t)$. Eq. vi simplifies to Eq. 8 in the main text for the identity link.

#### 2 Simulation studies

##### 2.1 Setup

Fig S3 shows the grids at which outcomes are generated in the parametric simulation studies, Fig S4 illustrates the spatial outcome patterns generated and Fig S5 the resulting univariate Moran's I. The coordinates of the *MAGPIE* alignments with jittered landmarks of the Vicari data are shown in Fig S6. Monte-Carlo variance on the proportions type I error, power and FDR for 50 replications with at least 10 independent feature pairs is at most $\frac{0.5\times0.5}{50\times10} = 0.0005$.

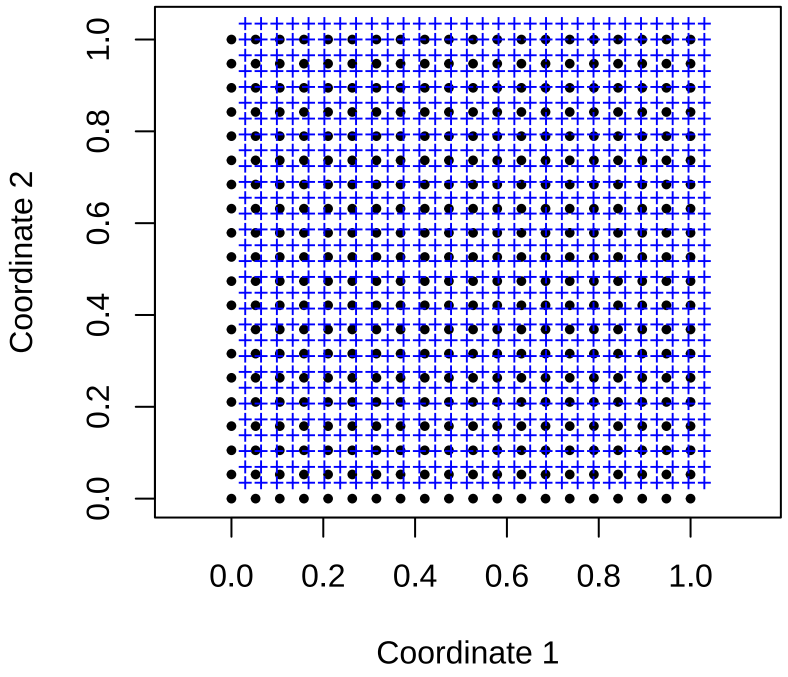

Fig S3: Spot grids for the parametric simulation study. The black squares are the locations of the spots of the first modality (X) and of the second (Y) for joint coordinate sets, the blue crosses the locations of the second modality (Y) for disjoint coordinate sets.

In an additional parametric simulation featuring anisotropy and nonstationarity, we generate outcomes according to the "independentGPs" and "dependentGPs" scenarios, but construct the autocorrelation matrices as follows. $\mathbf{\Sigma}_x = \sigma_x^2 \left( \pi_x \mathbf{I} + (1-\pi_x) \exp \left[ -\left( \frac{\mathbf{D}_{xc}}{l_x + \mathbf{c}_1 \mathbf{c}_1^t \xi} \right)^2 - \left( \frac{\mathbf{D}_{xe}}{l_x \gamma} \right)^2 \right] \right)$ with $\mathbf{D}_{xc}$ the distances in the first dimension and $\mathbf{D}_{xe}$ the distances in the second, $\gamma \in (0.5, 1, 2)$ the anisotropy parameter, $\xi \in (0, 0.1, 0.2, 0.3)$ the nonstationarity parameter and $l_x = l_y = 0.15$ as before. The parameter combination $\gamma = 1$ and $\xi = 0$ recovers the isotropic, stationary scenario used throughout the parametric simulations in the main text. The bivariate spatial covariance between $x$ and $y$ is constructed as before. The design is illustrated for all combinations of such parameters in Fig S7.

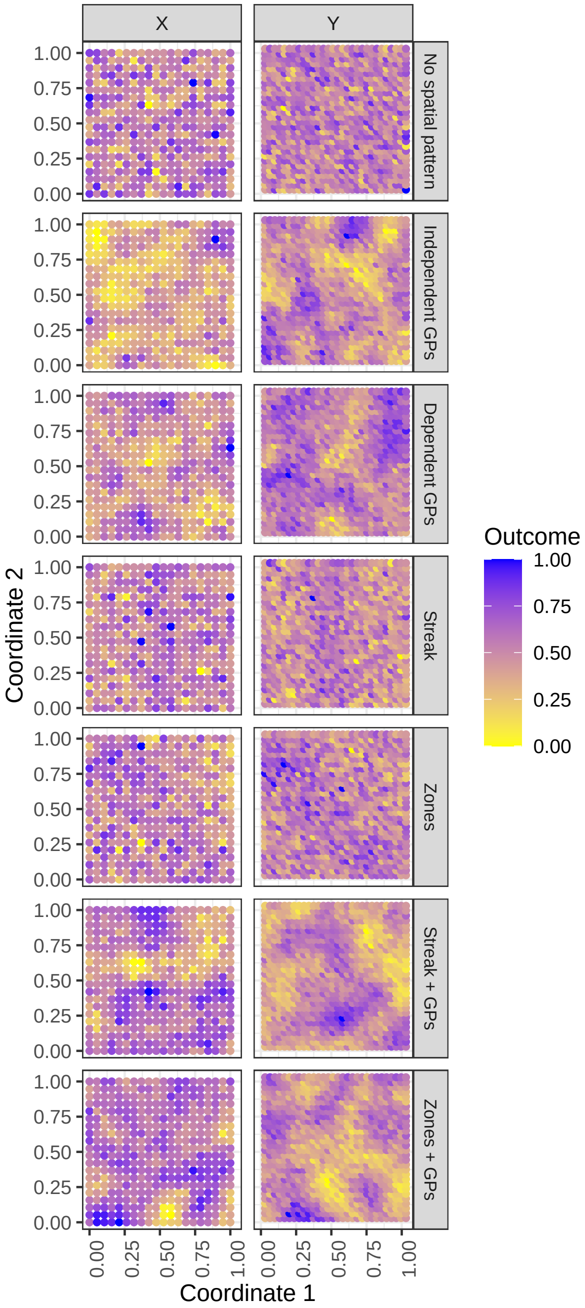

Fig S4: Visualisation of the different designs (rows) and outcomes (columns) for the parametric simulation study with disjoint coordinate sets. Outcomes are scaled to the [0,1] range for legibility.

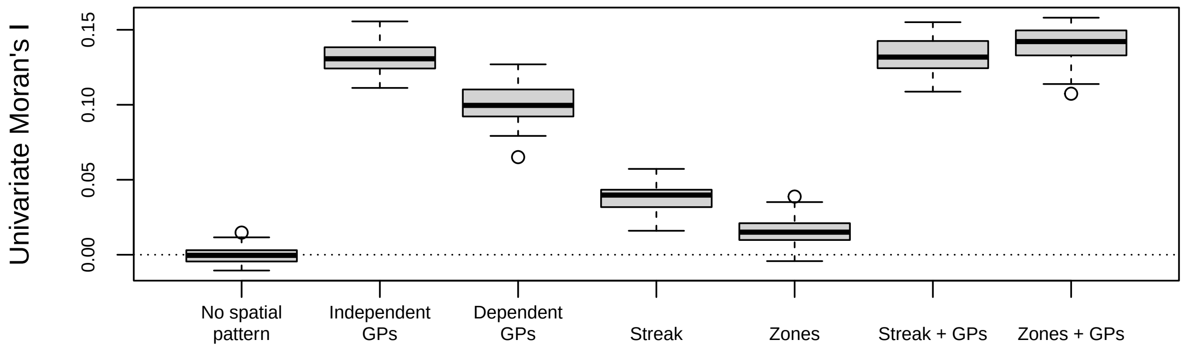

Fig S5: Boxplots of univariate Moran's I values of the $x$-variable (y-axis) from the designs in Fig S4 (x-axis), over 50 Monte-Carlo instances.

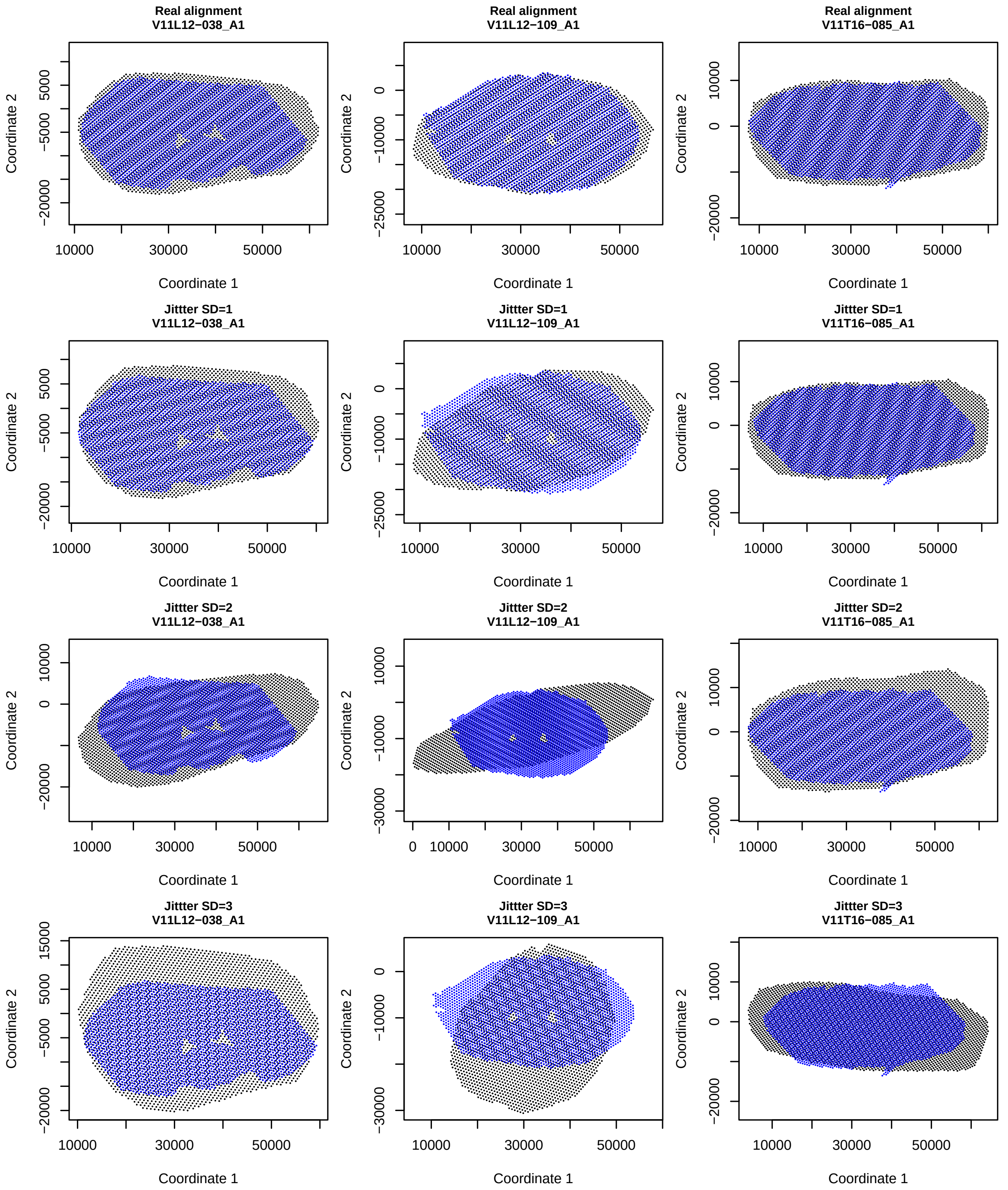

Fig S6: Real alignments (top plots) and examples of alignment after jittering landmarks with different standard deviations (SD) (other plots) of the first section of the three mice in the Vicari data (plot titles). Metabolic spots are black and transcriptomic spots are blue.

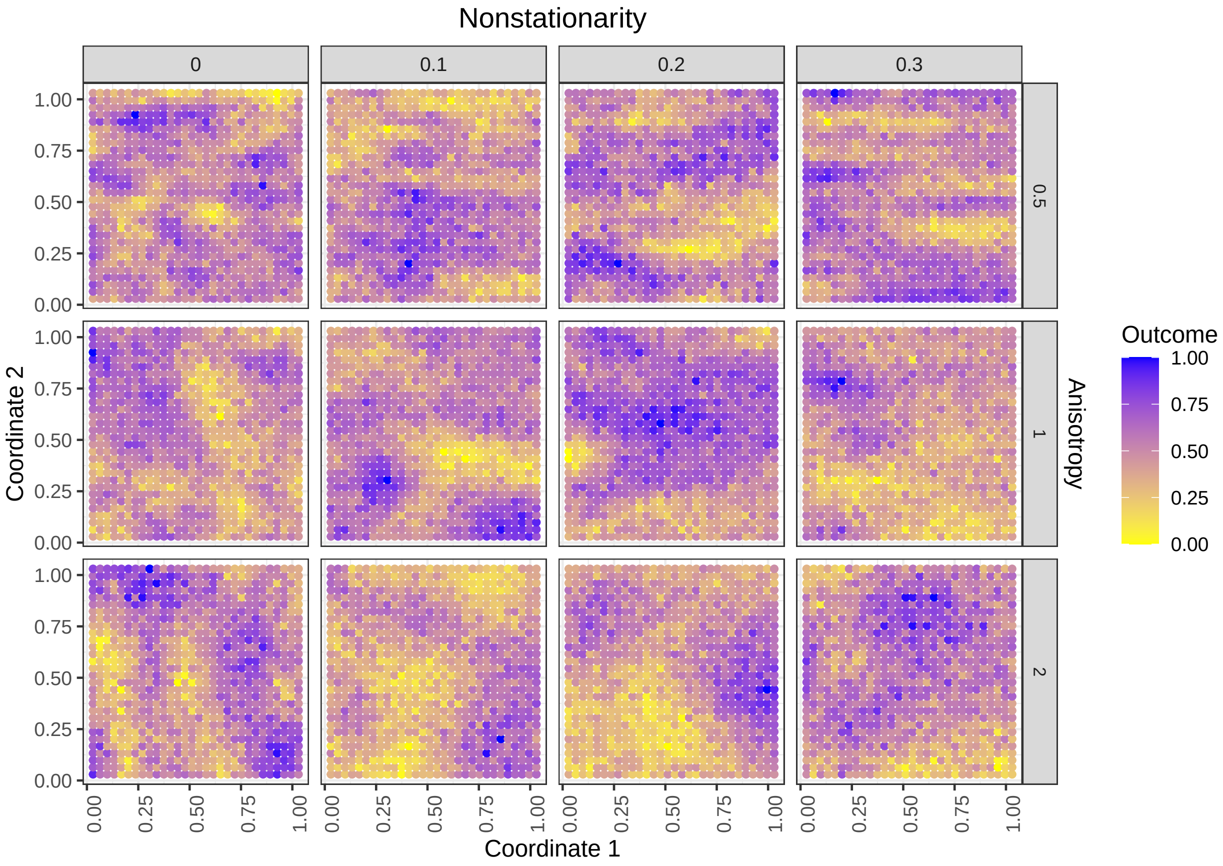

Fig S7: Visualisation of the different designs for the parametric simulation study with disjoint coordinate sets under anisotropy and nonstationarity. Rows show the anisotropy parameter $\gamma$, columns the nonstationarity parameter $\xi$. Outcomes of the y-variable are shown, scaled to the [0,1] range for legibility.

#### 2.2 Results

##### 2.2.1 Nonparametric simulations

**2.2.1.1 Single-image analysis** Fig 2 in the main text summarizes the results of the nonparametric simulations on single images, here we show more exhaustive results. Monte-Carlo variance of the estimated proportions, type I error and FDR, for 50 replicates is at most $\frac{0.5\times0.5}{50} = 0.005$. Figs S8-S9 demonstrate that bivariate Moran's I (random shift), bivariate Moran's I (randomisation) and GAMs when both modalities are permuted, often have p-value distributions that are larger than uniform, whereas many existing methods have smaller-than-uniform p-value distributions. Next we investigate which features are prone to incur false findings. As measure of SAC, we calculate univariate Moran's I values $\mathbf{X}^t\mathbf{W}\mathbf{X}$ for all features of the original data, constructing the weight matrix as for the bivariate case, with Gaussian weights and decay parameter $\eta = 10^{-3}$, and setting the diagonal to zero. Boxplots of the univariate Moran's I values thus calculated are shown in Figs S10-S11, showing considerable SAC in almost all images, with the metabolome exhibiting the strongest SAC. We model the probability that a feature is detected as significantly associated with a permuted feature from the other modality through generalized linear mixed models using the *glmer* function in the *lme4* package [27] with a binomial outcome distribution and logit link. Univariate Moran's I of the non-permuted feature was included as fixed effect; random intercepts and random slopes for univariate Moran's I were included for the sections. Separate models were fitted per method, permuted modality and dataset. Modeled probabilities of detection as a function of univariate Moran's I are plotted in Fig S12, revealing that for the methods found liberal in the simulations, features with high SAC in the non-permuted modality are more likely to incur false findings.

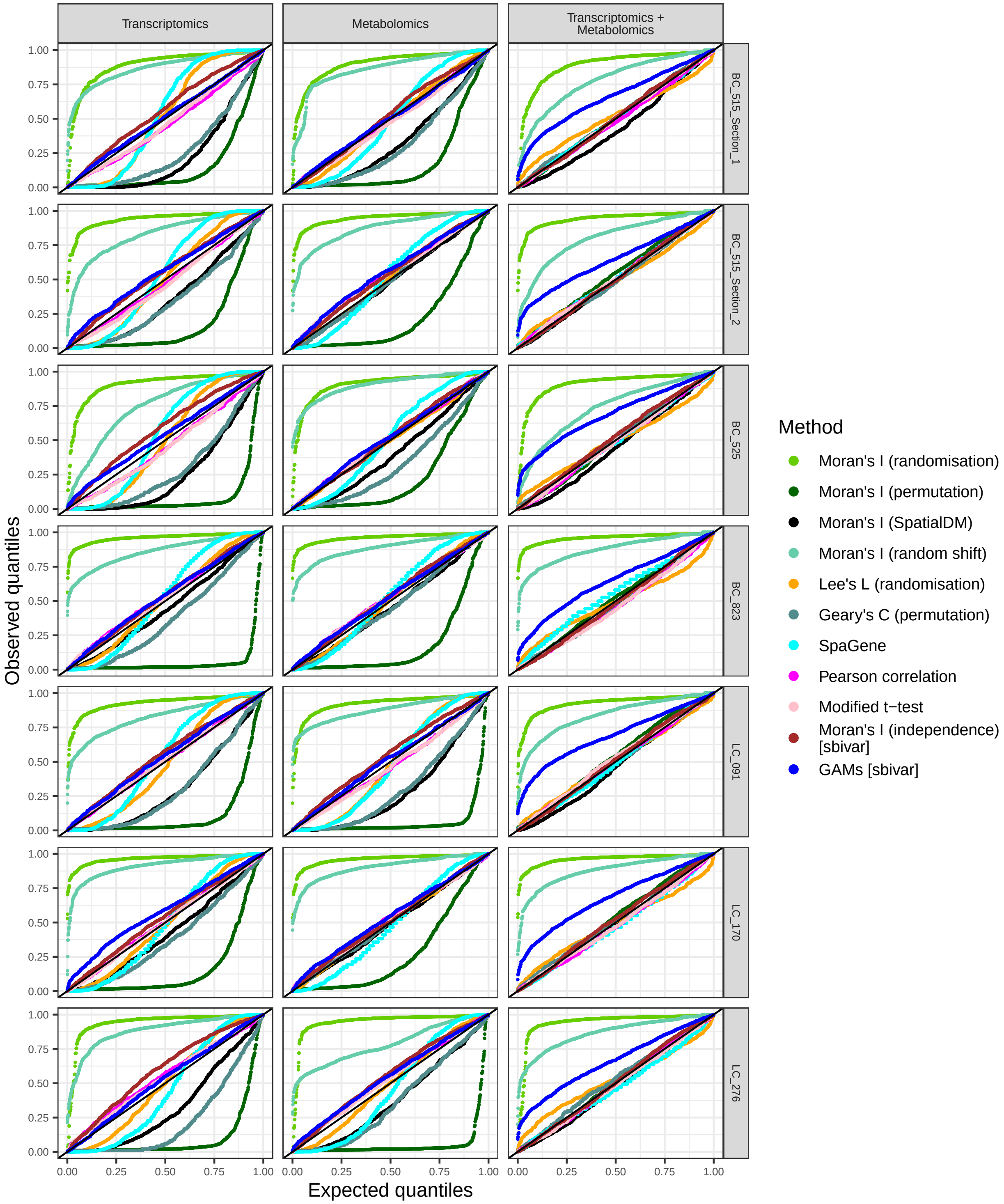

Fig S8: QQ-plots of observed p-values (y-axis) versus standard uniform quantiles (x-axis) for nonparametric simulations based on the Godfrey data, for different methods (colour) when permuting observations of one or two modalities (columns) of the sections (rows). The solid black line has slope 1 and intercept 0.

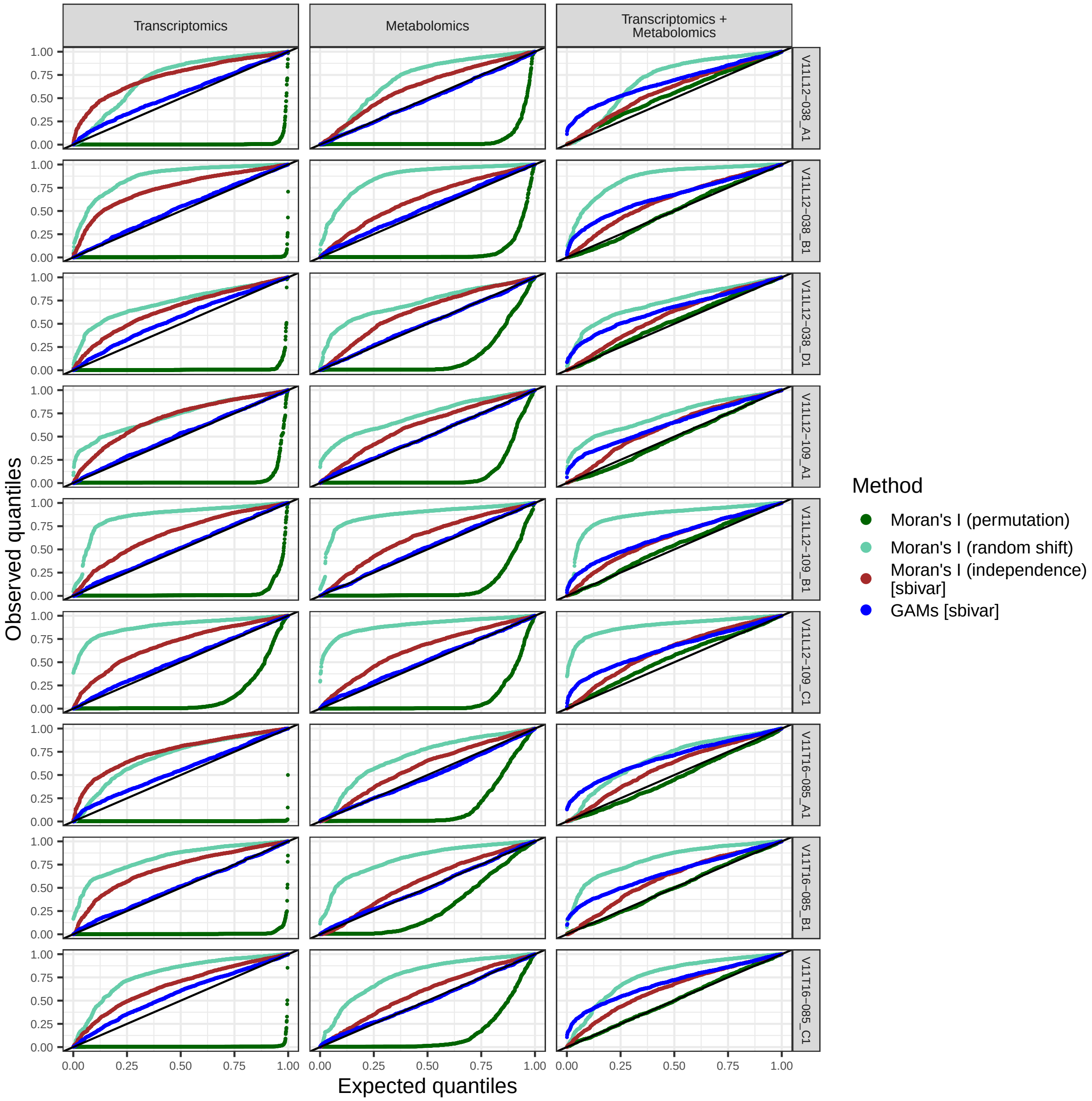

Fig S9: QQ-plots of observed p-values (y-axis) versus standard uniform quantiles (x-axis) for nonparametric simulations based on the Vicari data, for different methods (colour) when permuting observations of one or two modalities (columns) of the sections (rows). The solid black line has slope 1 and intercept 0.

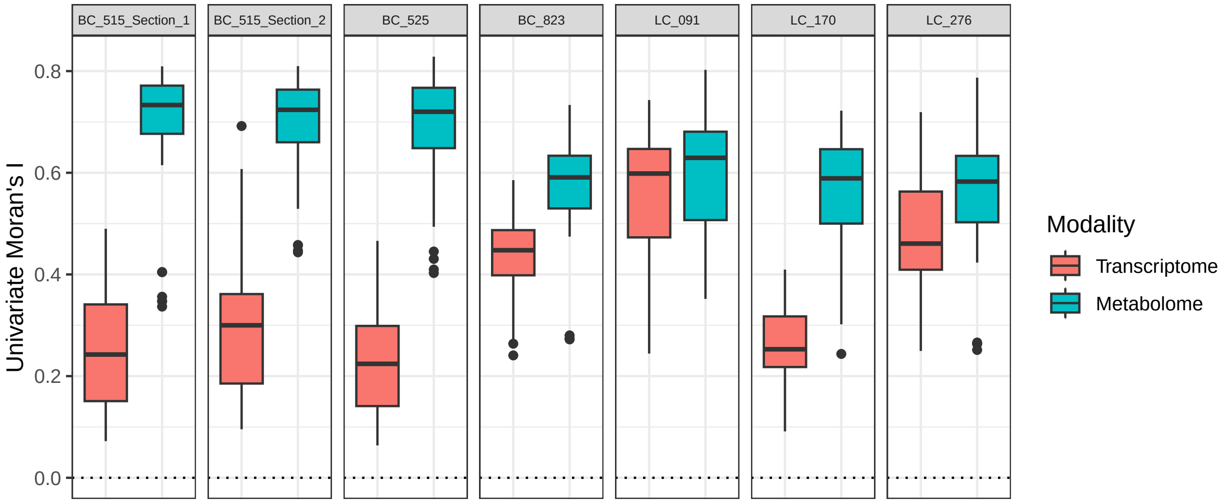

Fig S10: Boxplots of univariate Moran's I (y-axis) of the 30 most abundant features per section (columns) in the Godfrey data. The dotted horizontal line indicates 0.

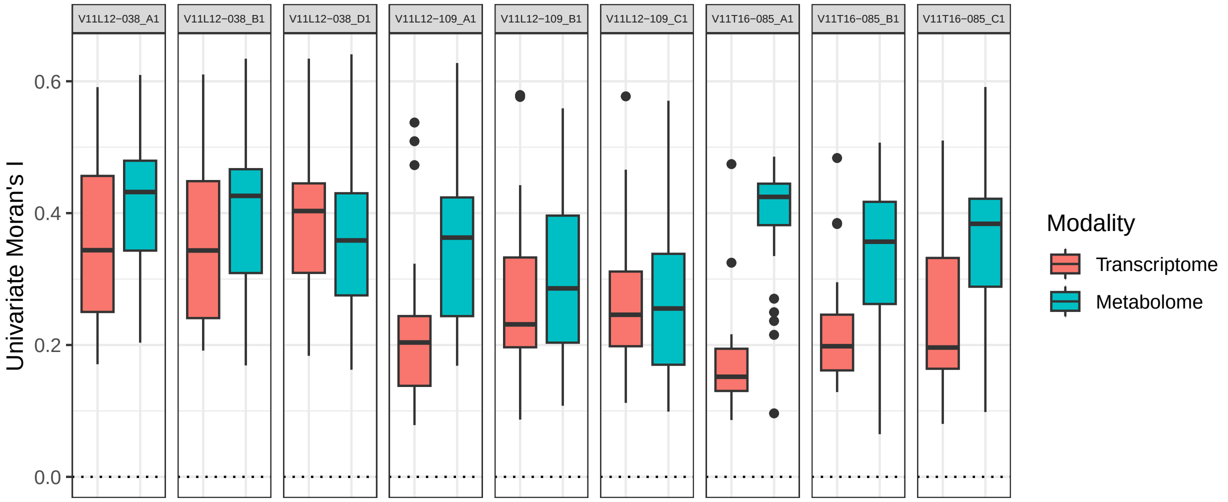

Fig S11: Boxplots of univariate Moran's I (y-axis) of the 30 most abundant features per section (columns) in the Vicari data. The dotted horizontal line indicates 0.

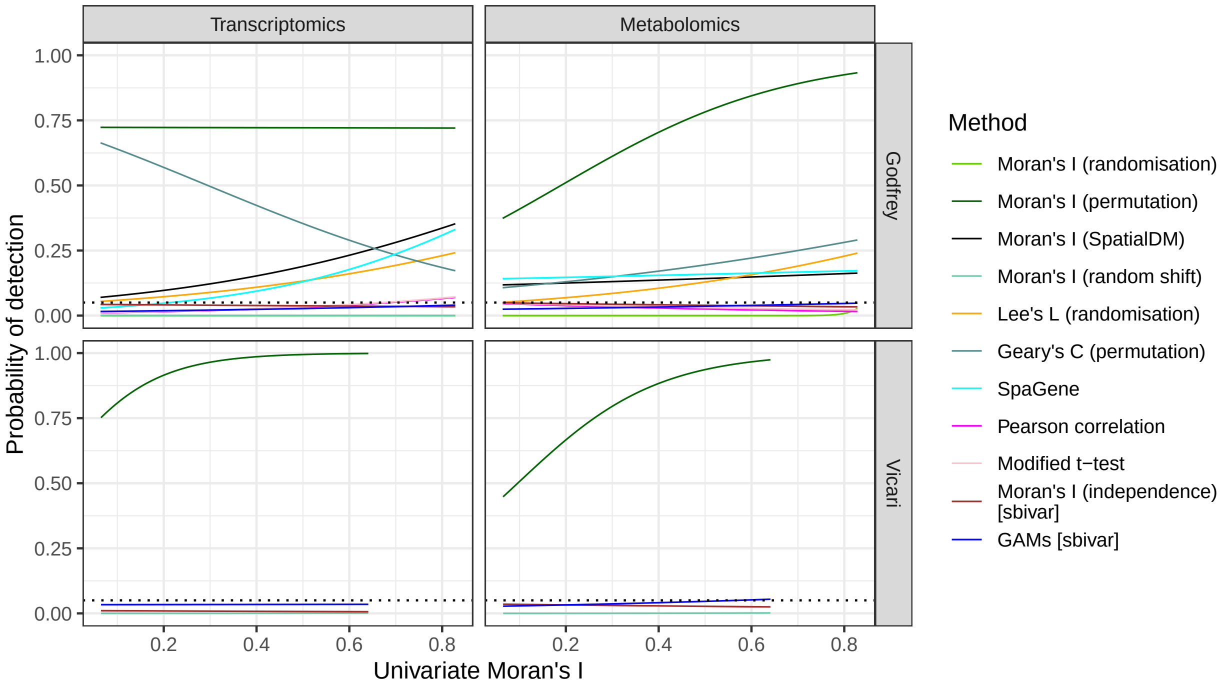

Fig S12: Predicted probability of detection as significant finding (y-axis) as a function of univariate Moran's I (x-axis) for different methods (colour), non-permuted modalities (columns) and dataset (rows) in the nonparametric simulation, modeled by a generalized linear mixed model with binomial outcome distribution (see text for details). The predictions are only shown within the range of observed univariate Moran's I values and for methods that made any findings. The horizontal dotted line indicates the significance level of 0.05.

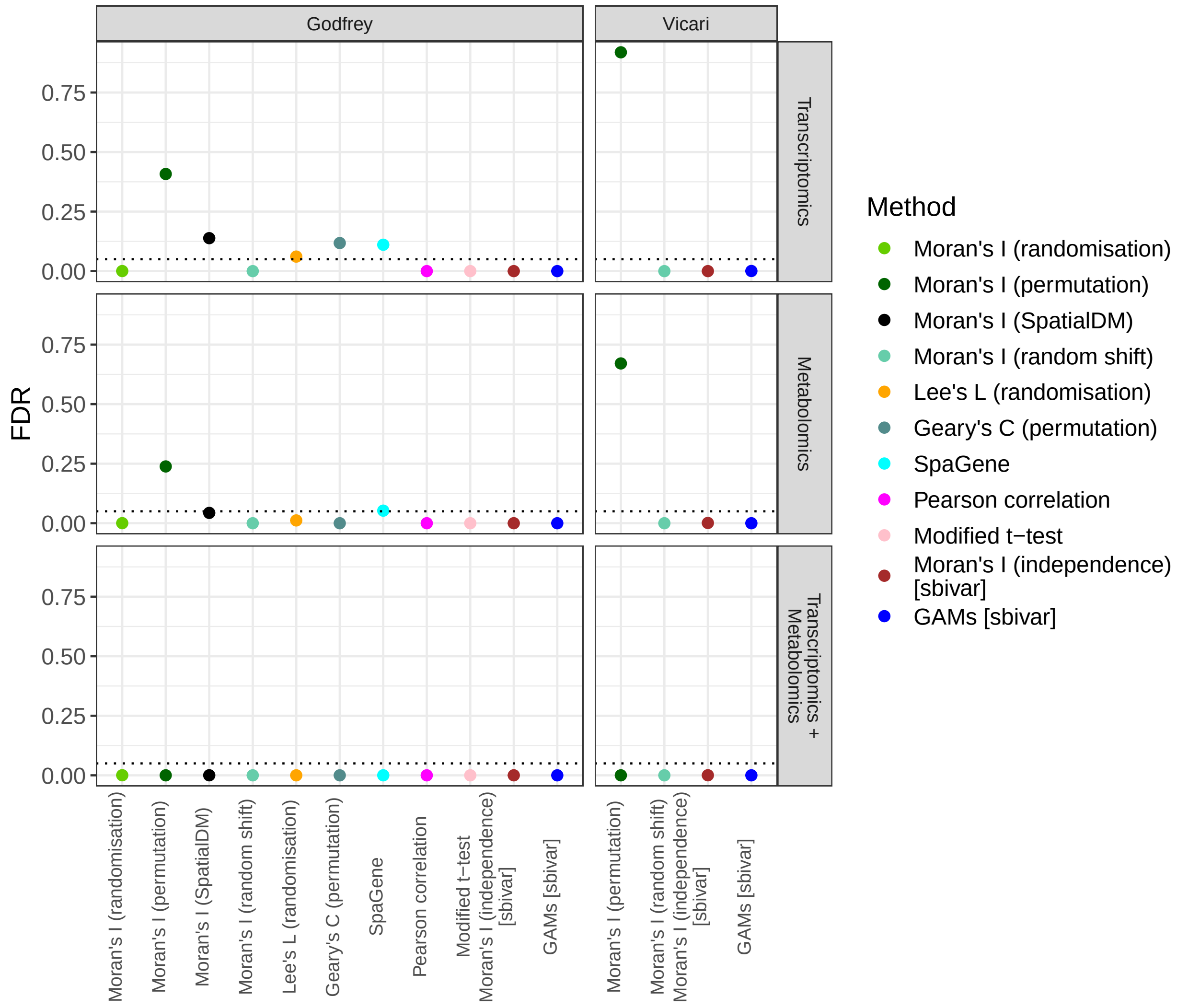

Fig S13: FDR (y-axis) for different methods with null distributions in parentheses (colour) for one or two permuted modalities (rows) averaged over all samples of the Godfrey and Vicari data (columns). The dotted horizontal line indicates the nominal FDR of 0.05, [sbivar] indicates methods introduced in this work.

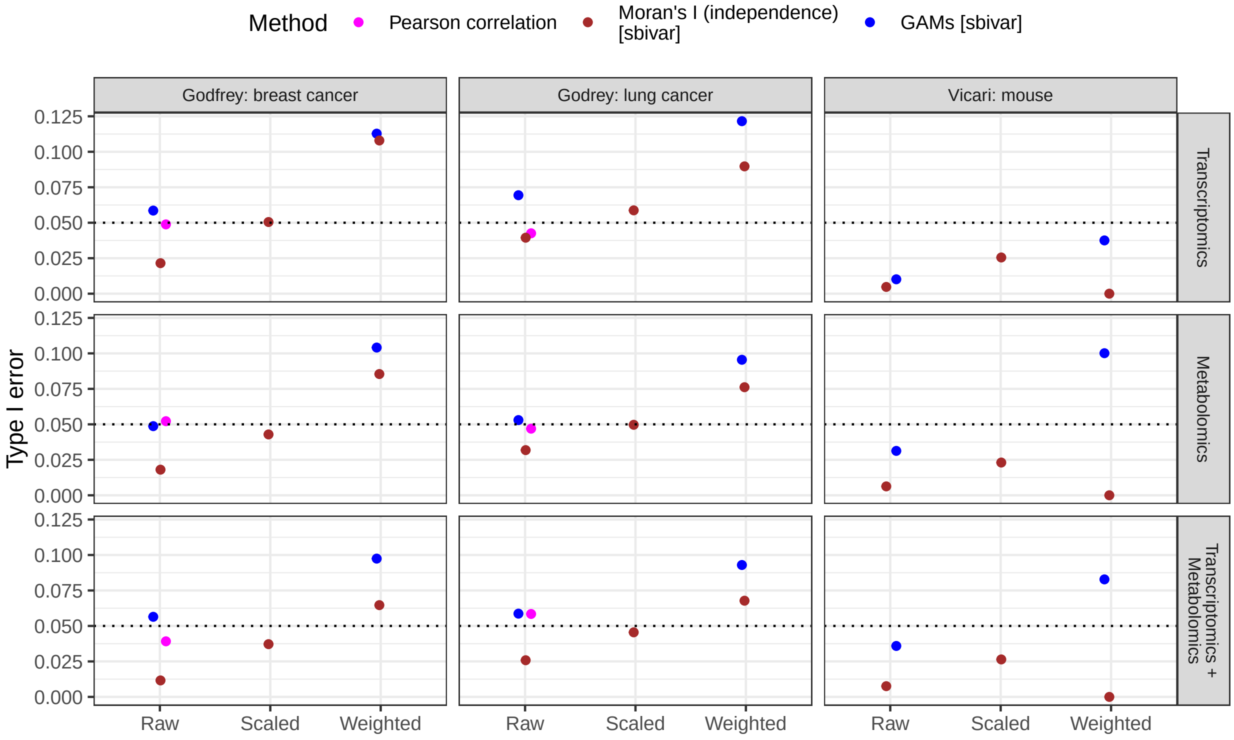

Fig S14: Type I error (y-axis) of different methods (colour) and weighting schemes (x-axis) for multi-image analysis of the Godfrey and Vicari data (columns) after scrambling one or two modalities (rows).

###### 2.2.1.2 Multi-image analysis

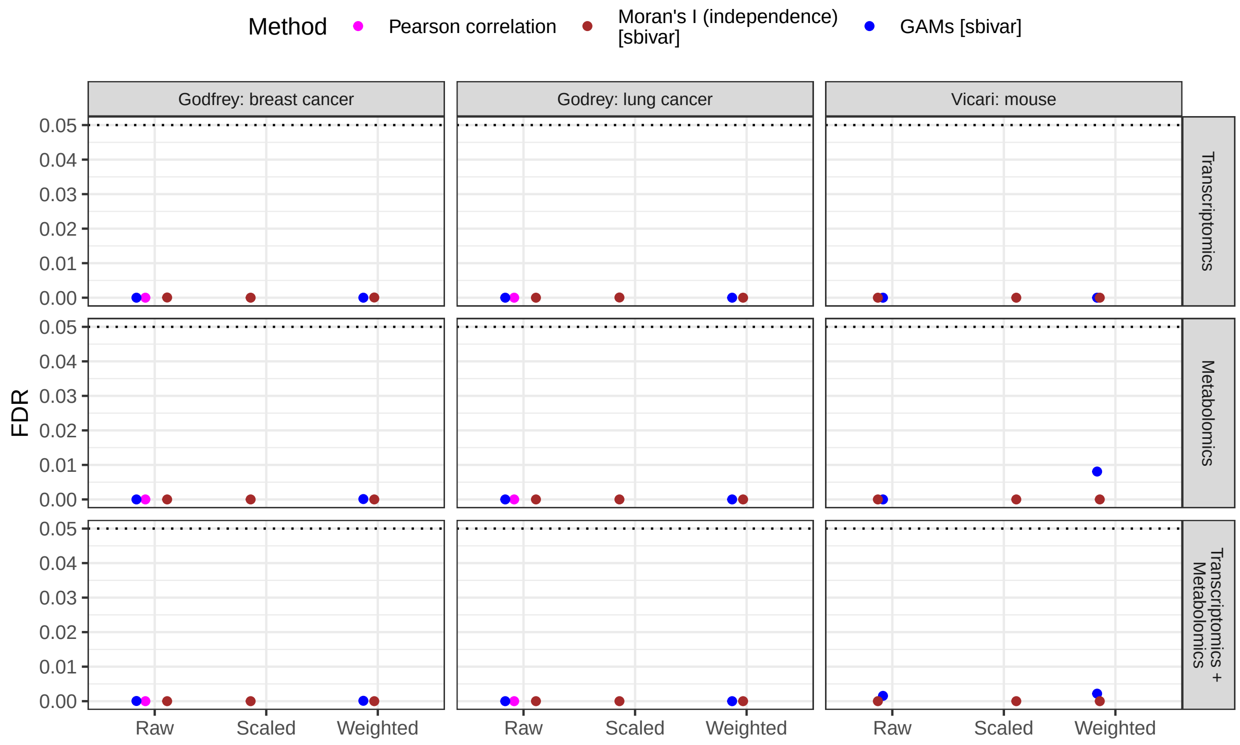

Fig S15: FDR (y-axis) of different methods (colour) and weighting schemes (x-axis) for multi-image analysis of the Godfrey and Vicari data (columns) after scrambling one or two modalities (rows). The horizontal dashed line indicates the nominal FDR of 0.05.

##### 2.2.2 Parametric simulations

**2.2.2.1 Single-image analysis** Fig S16 demonstrates empirically that our variance estimator $tr(\mathbf{W}^t\mathbf{\Sigma}_x\mathbf{W}\mathbf{\Sigma}_y)$ is accurate, whereas the variance estimator assuming i.i.d. observations ($tr(\mathbf{W}^t\mathbf{W})$ as in *SpatialDM* by Li et al. [2] and by DeTomaso et al. [3]) underestimates the variance in presence of SAC. Fig S17 shows qq-plots of the p-values for all methods the null scenarios. Fig S18 demonstrates how Pearson correlation and other methods suffer from type I error inflation only when there is SAC in both variables. Fig S22 shows average time consumption for the different methods on simulated datasets in the "no spatial pattern" scenario with joint coordinate sets for the number of spots increasing as 400, 625, 900, 1225, 1600, 2025, over 20 replicates, on an Intel Core i5-11400H 2.70GHz processor.

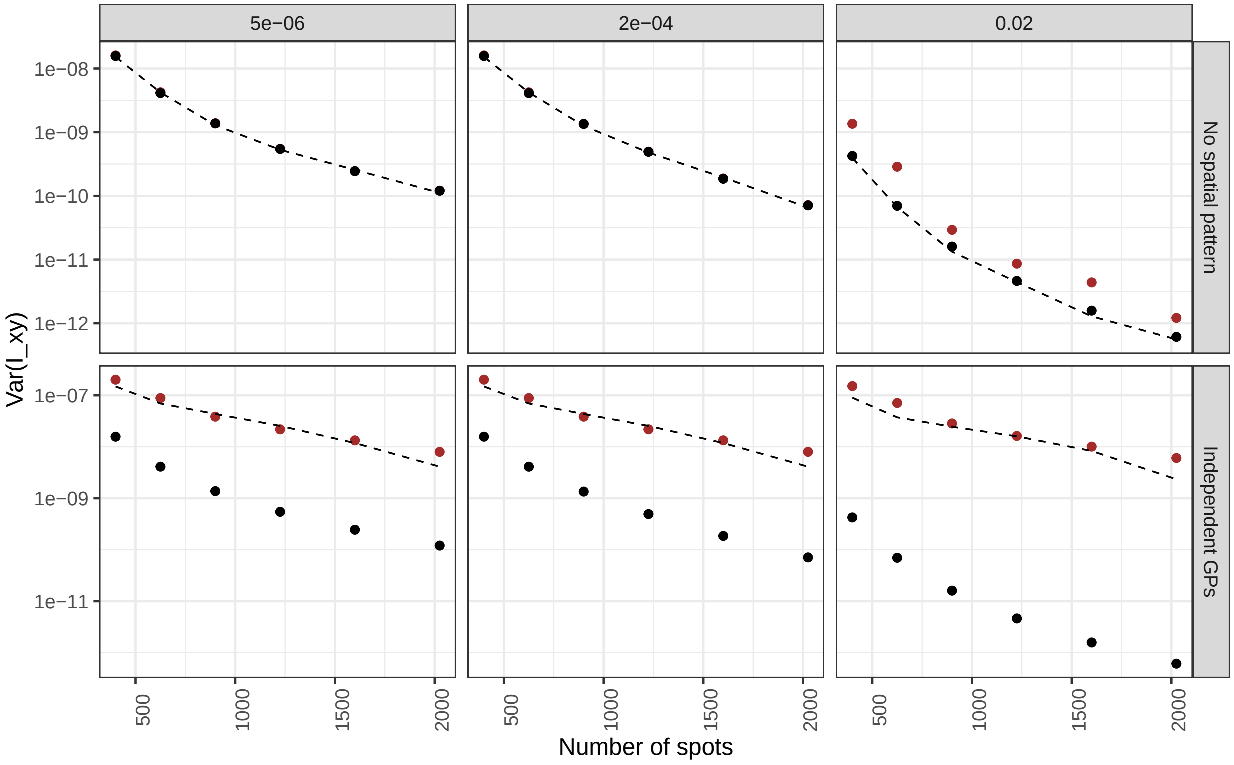

Fig S16: Empirical variances of bivariate Moran's I (y-axis, dashed black line) as a function of number of spots (x-axis), decay parameter of the weight matrix (colums) and spatial pattern under the independence null (rows). The black dots indicate the estimated variance under the randomisation null, the brown dots are *sbivar*'s estimates under the independence null. The y-axis is on the log scale.

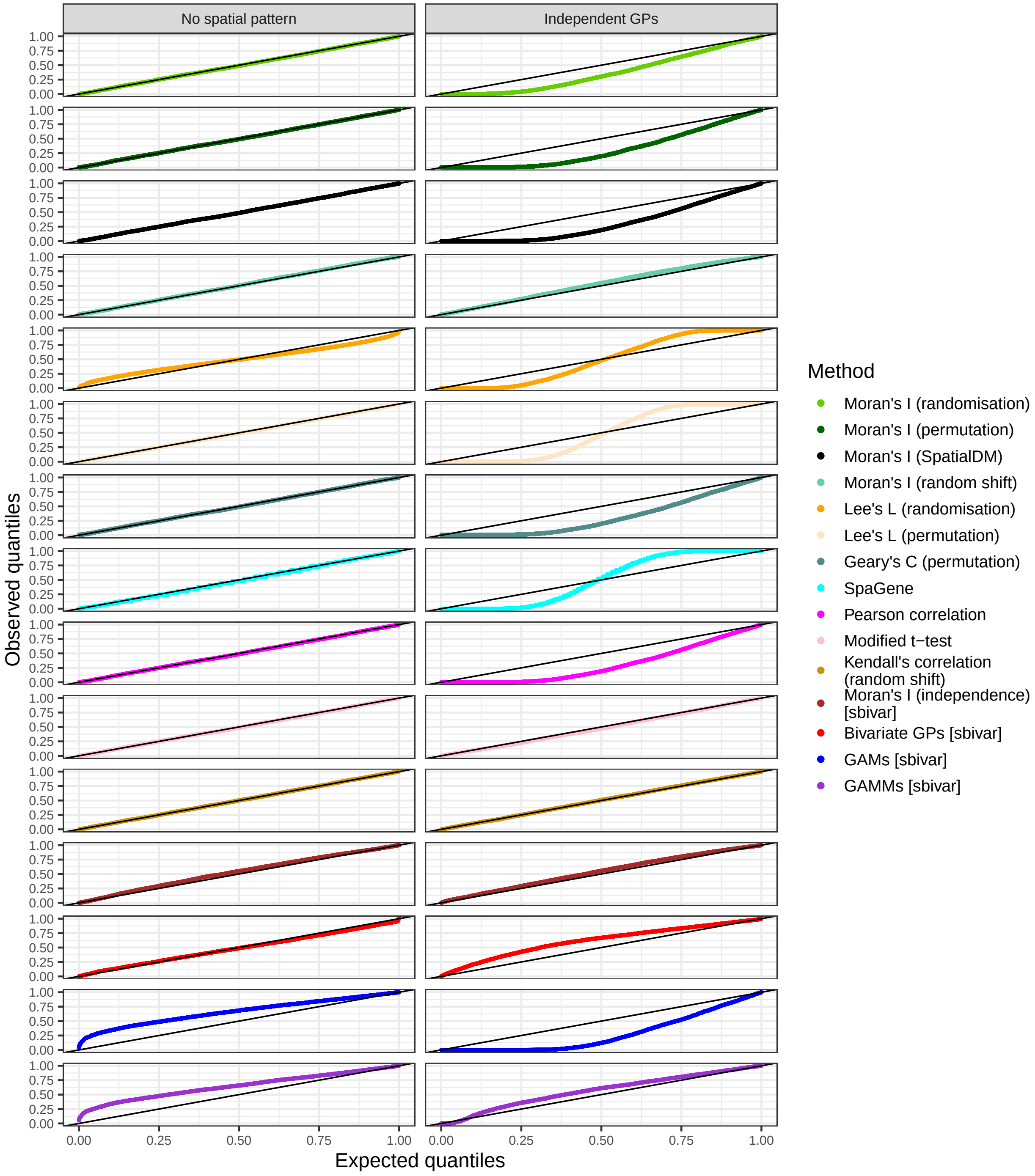

Fig S17: QQ-plots of observed p-values (y-axis) versus standard uniform quantiles (x-axis) for parametric simulations under the two null scenarios (columns), for different methods (colour). The solid black line has slope 1 and intercept 0.

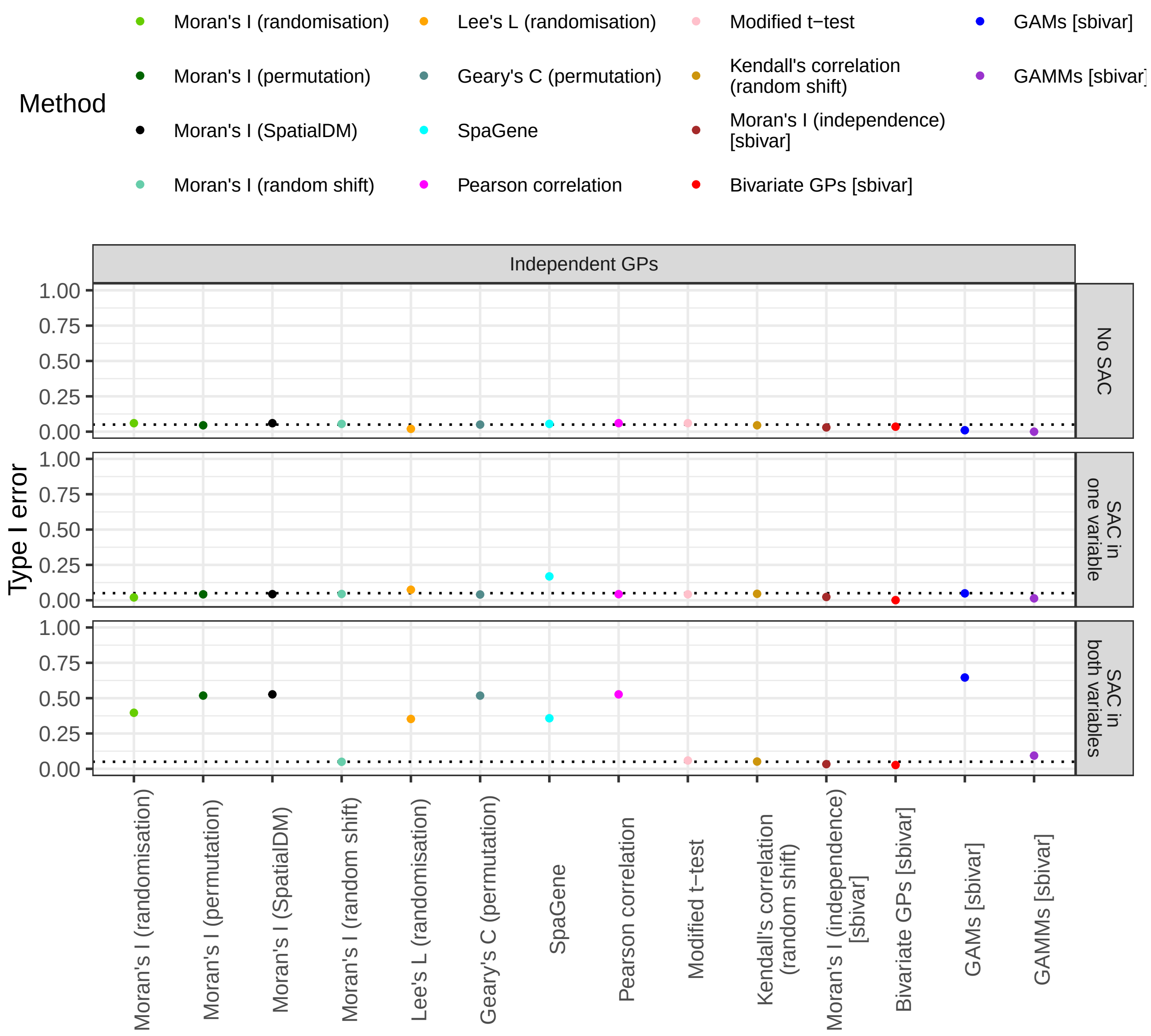

Fig S18: Type I error (y-axis) for different analysis methods (x-axis, colour) as a function of whether there is no SAC, SAC in one or in both variables (rows), for outcomes generated as independent GPs. The horizontal dotted line indicates the significance level of 5%.

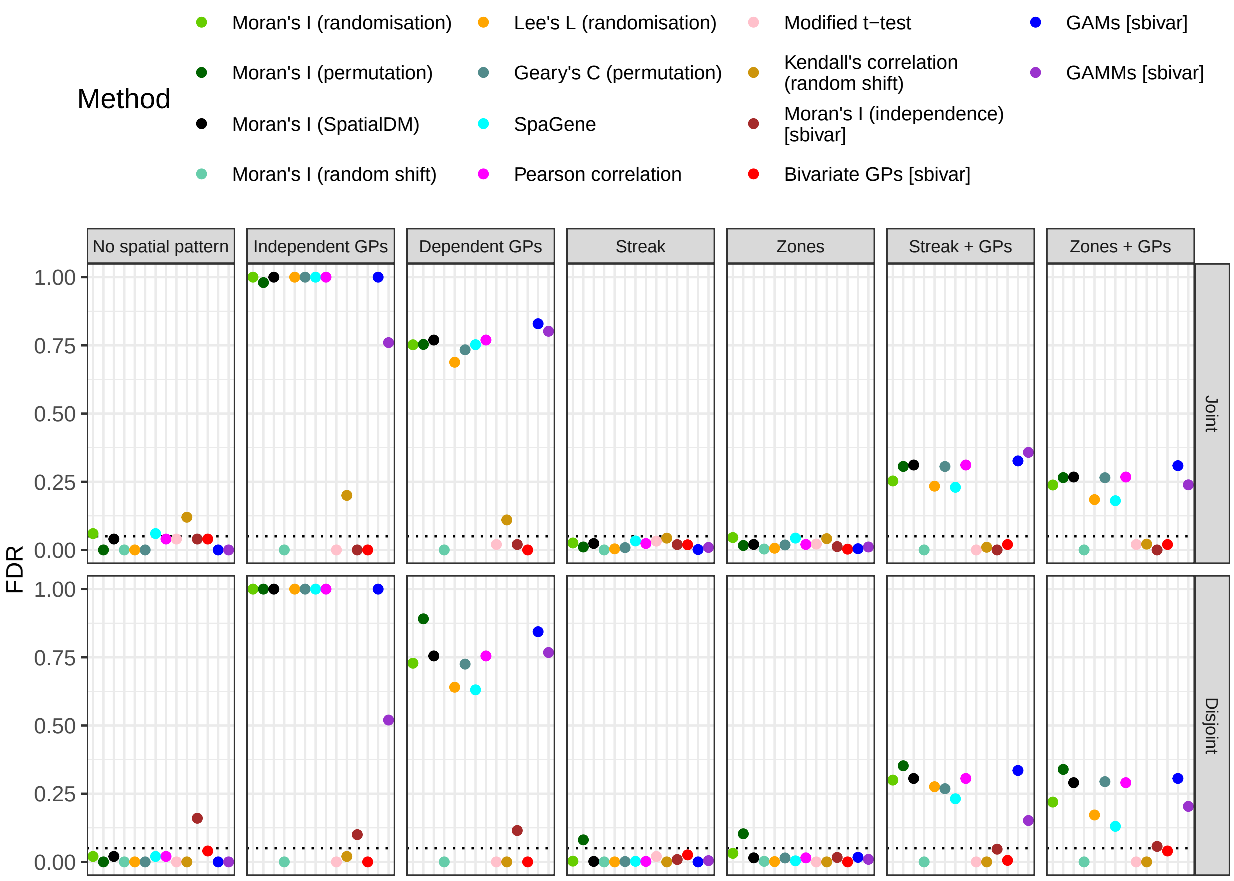

Fig S19: FDR (y-axis) for different methods with null distributions in parentheses (colour, x-axis) for data with different spatial patterns (columns) in parametric simulations with joint and disjoint coordinate sets (rows). The horizontal dotted line indicates the nominal FDR of 5%, [sbivar] indicates methods introduced in this work.

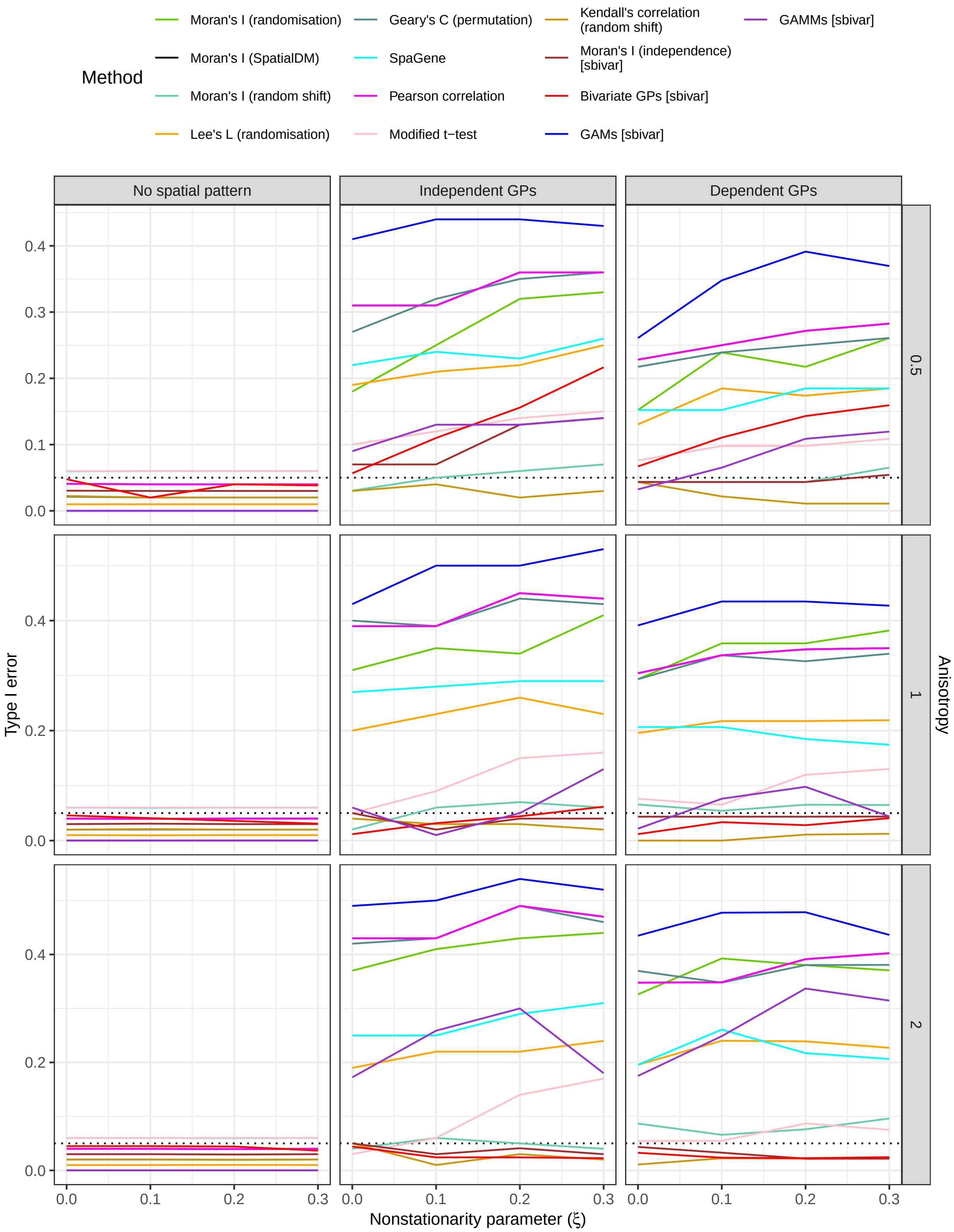

Fig S20: Type I error (y-axis) for different methods with null distributions in parentheses (colour) for data with different spatial patterns (columns), nonstationarity parameter $\xi$ (x-axis) and anisotropy parameter $\gamma$ (rows) in the parametric simulation with anisotropy and nonstationarity. The horizontal dotted line indicates the significance level of 5%, [sbivar] indicates methods introduced in this work.

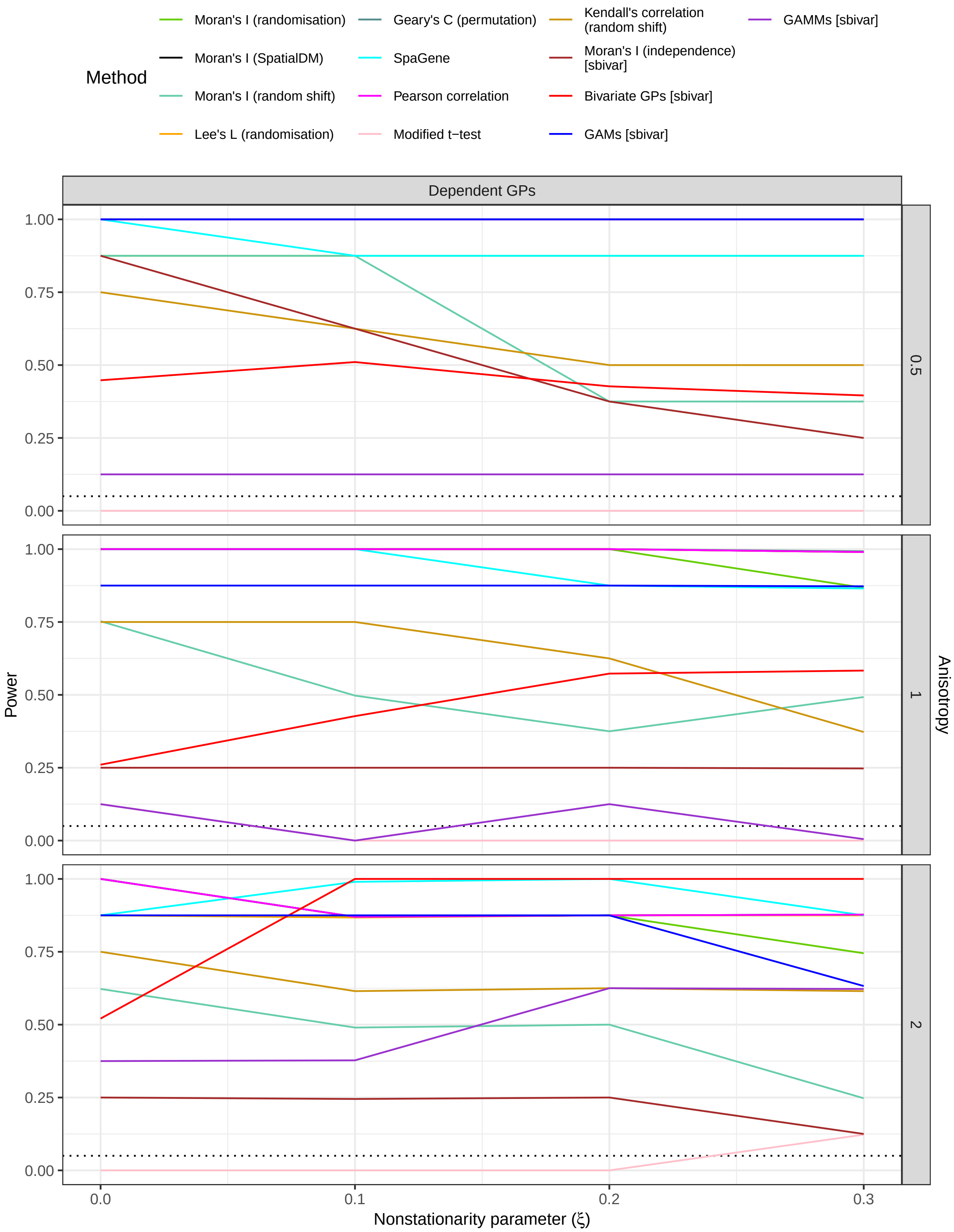

Fig S21: Power (y-axis) for different methods with null distributions in parentheses (colour) for data generated as dependent GPs, nonstationarity parameter $\xi$ (x-axis) and anisotropy parameter $\gamma$ (rows) in the parametric simulation with anisotropy and nonstationarity. The horizontal dotted line indicates the significance level of 5%, [sbivar] indicates methods introduced in this work.

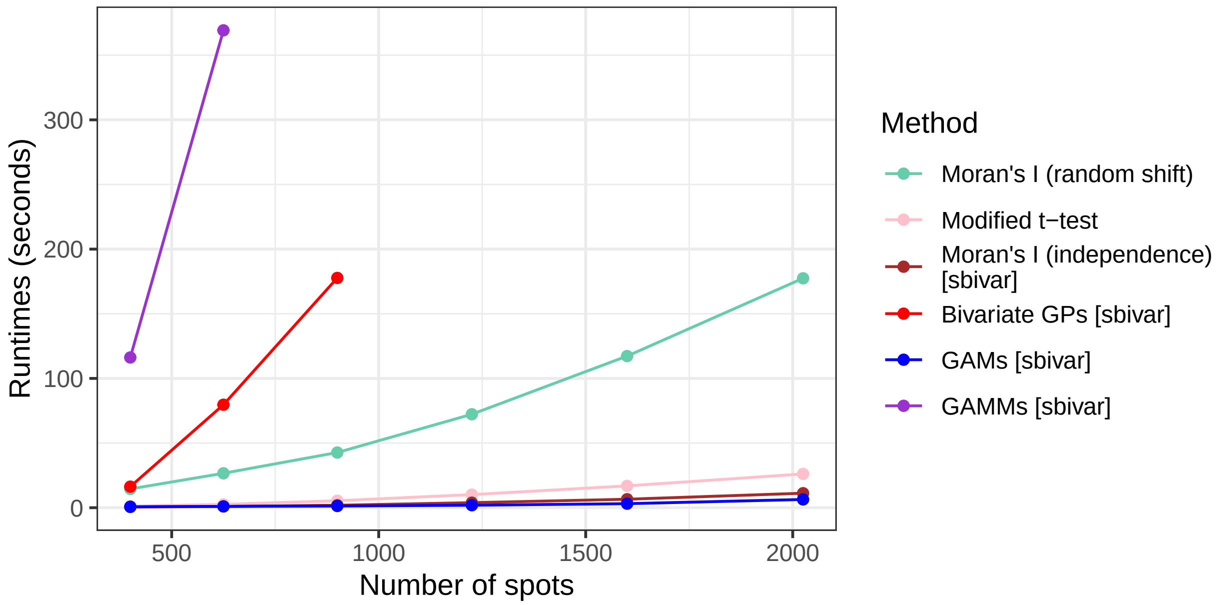

Fig S22: Runtimes (y-axis, in seconds) for different methods that control type I error rate (colour) and increasing number of spots (x-axis) with joint coordinate sets in the ‘No spatial pattern’ scenario. Bivariate GPs and GAMMs were not run for the higher spot numbers.

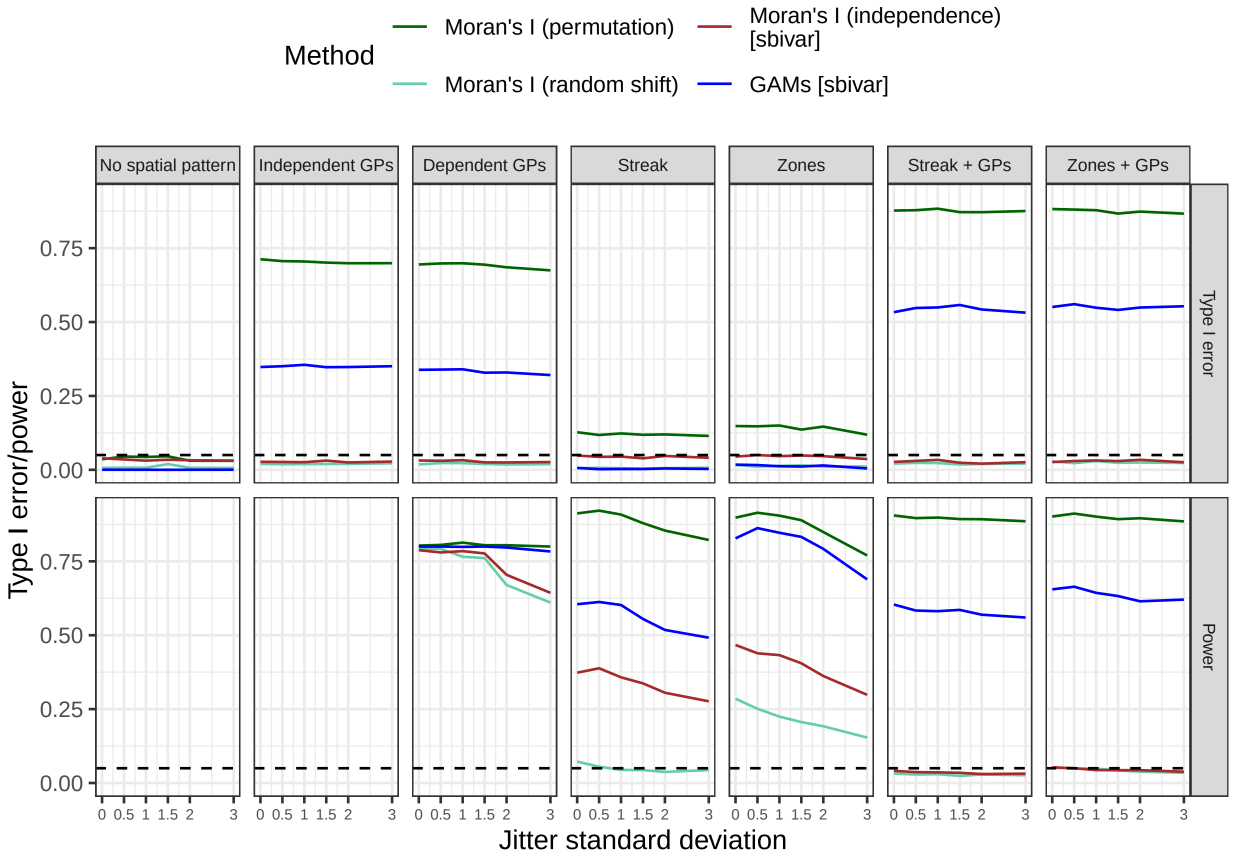

Fig S23: Type I error and power (y-axis, rows) averaged over all nine Vicari data template sections as a function of standard deviation by which the landmarks were jittered (x-axis) for different methods (colour) and spatial patterns (columns). A standard deviation of 0 corresponds to an analysis using the true, unjittered coordinates.

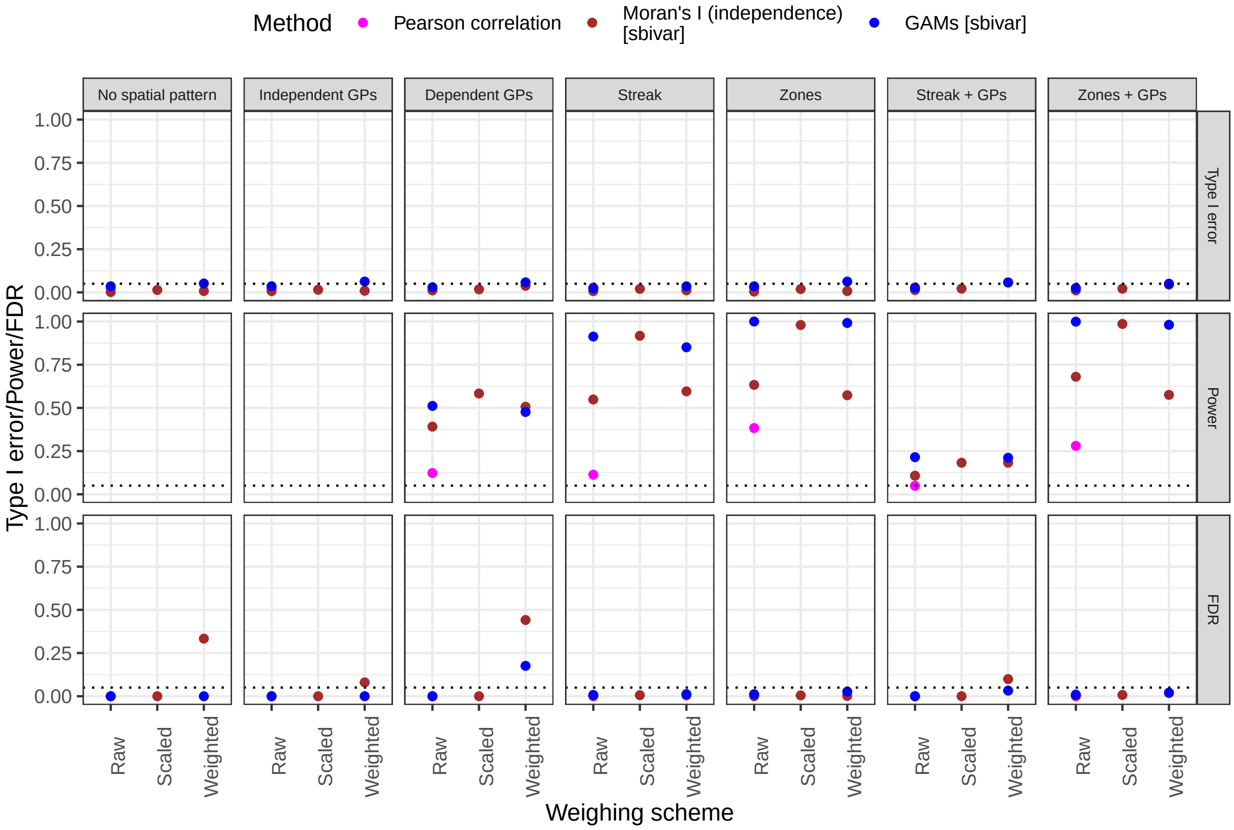

Fig S24: Type I error, power and FDR (y-axis, rows) for different methods (colours) and different spatial patterns (columns) in multi-image parametric simulations, as a function of weighing scheme used in the linear model (x-axis).

**2.2.2.2 Multi-image analysis** Fig S24 shows type I error and power for different methods under replication, proving the added value of scaling the Moran's I statistic by its maximum absolute value, but suggesting that inverse weighting by variance does not improve the power. Fig S25 shows a timing comparison on an Intel Core i5-11400H 2.70GHz processor.

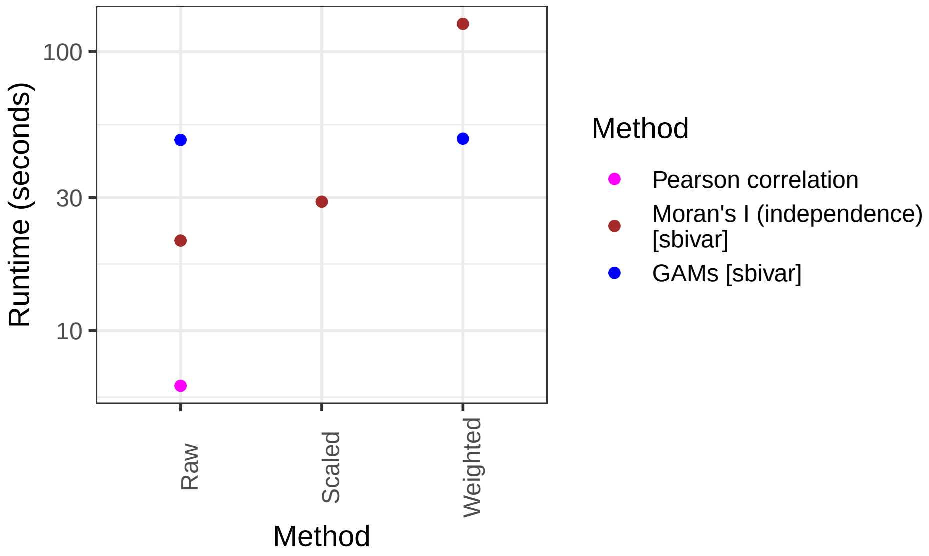

Fig S25: Average runtimes (y-axis, in seconds) for different methods (colour) and scaling schemes (x-axis) applied to multiple images with disjoint coordinate sets in the ‘No spatial pattern’ scenario of parametric simulations over 10 Monte-carlo instances with 20 features per modality.

### 3 Case studies

#### 3.1 Godfrey et al. (2025)

The dataset by Godfrey et al. [7] contains seven sections from human lung and breast cancer tissue coprofiled using desorption electrospray ionisation mass spectrometry imaging (DESI-MSI) for metabolites, followed by H&E staining and finally Visium for transcripts. The data were pre-aligned the help of the H&E stain and spot-matched by aggregating the metabolomics data with a Gaussian weighting scheme. Fig S26 reveals that the library sizes and total ion counts exhibit spatial patterning across the sections.

##### 3.1.1 Single-image analysis

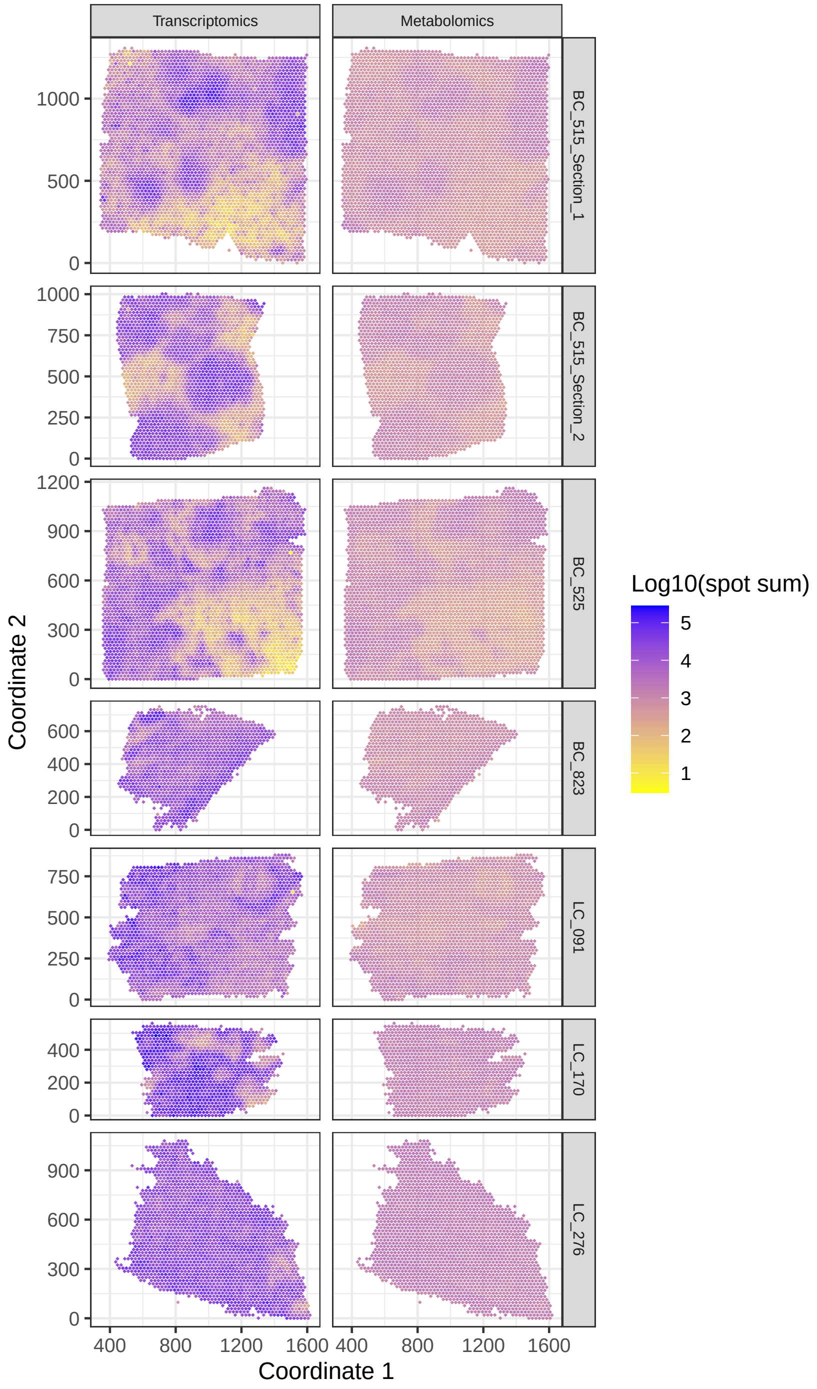

Fig S26: Log10 spot-wise sums (colour) of the different sections (rows) of the Godfrey data.

**3.1.1.1 Comparison to authors' findings** We can confirm all highest correlations discussed by the authors in the main text of their original publication. Differences in findings with the full supplementary results are shown in Fig 4 in the main text and Fig S27 here for findings only by our methods, and Figs S28-S29 for findings in the original analysis but not by our methods.

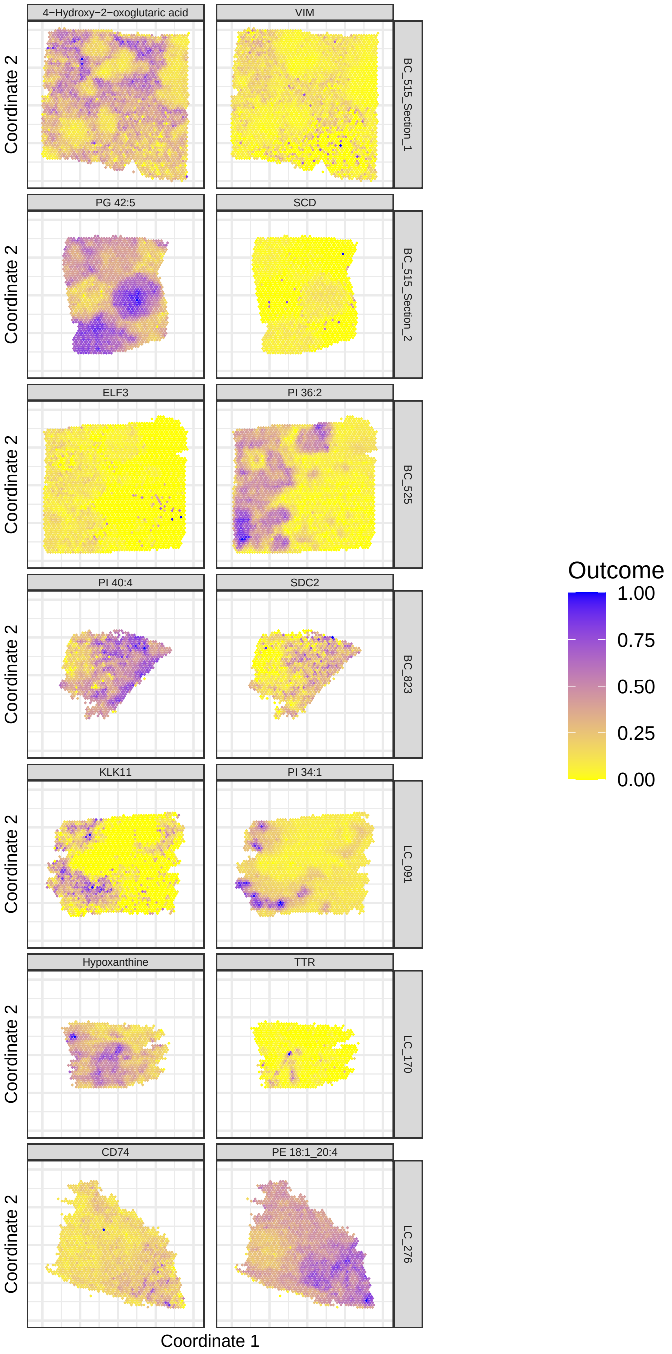

Fig S27: Gene-metabolite pairs found most significantly positively correlated by GAMs in different sections of the Godfrey data (rows), but not detected in the original analysis. Pairs in samples BC_515_Section_1, BC_525 and LC_276 were also found significant by Moran's I (independence).

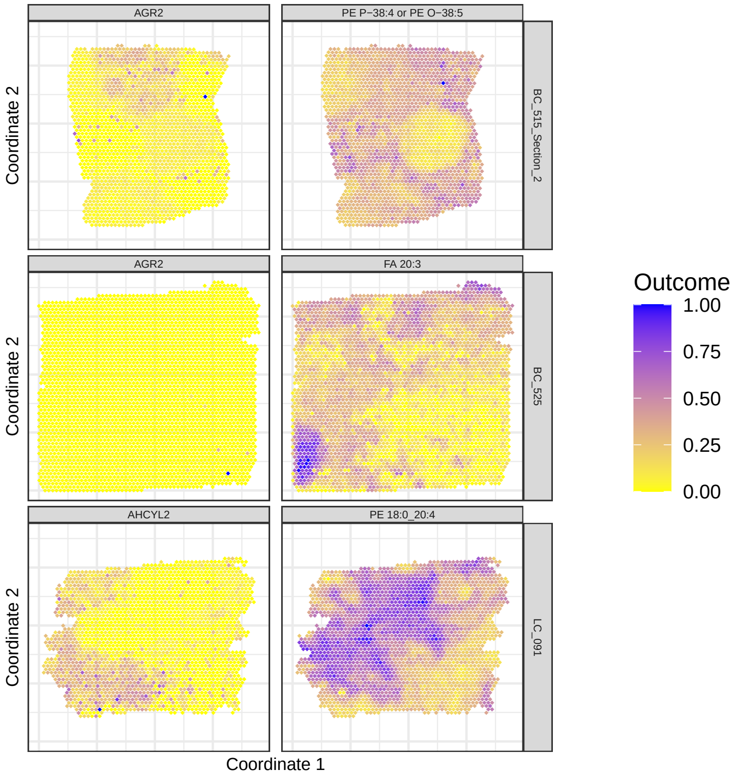

Fig S28: Gene-metabolite pairs with highest correlation found in the original analysis, but not declared significant by Moran's I (independence) or GAMs.

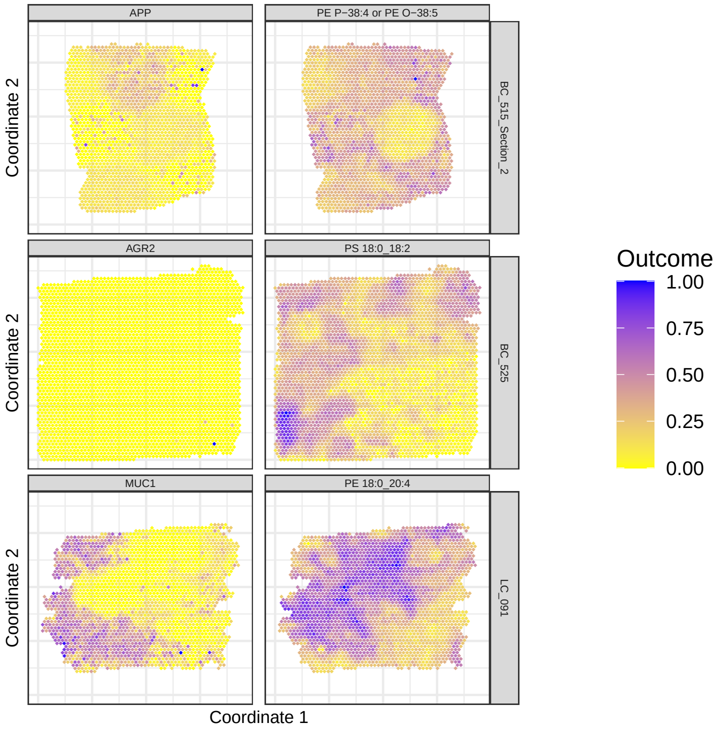

Fig S29: Gene-metabolite pairs with second highest significant correlation found in the original analysis, but not declared significant by Moran's I (independence) or GAMs.

###### 3.1.1.2 New findings

New findings by GAMs are shown in Figs S30-S31 and Table S1.

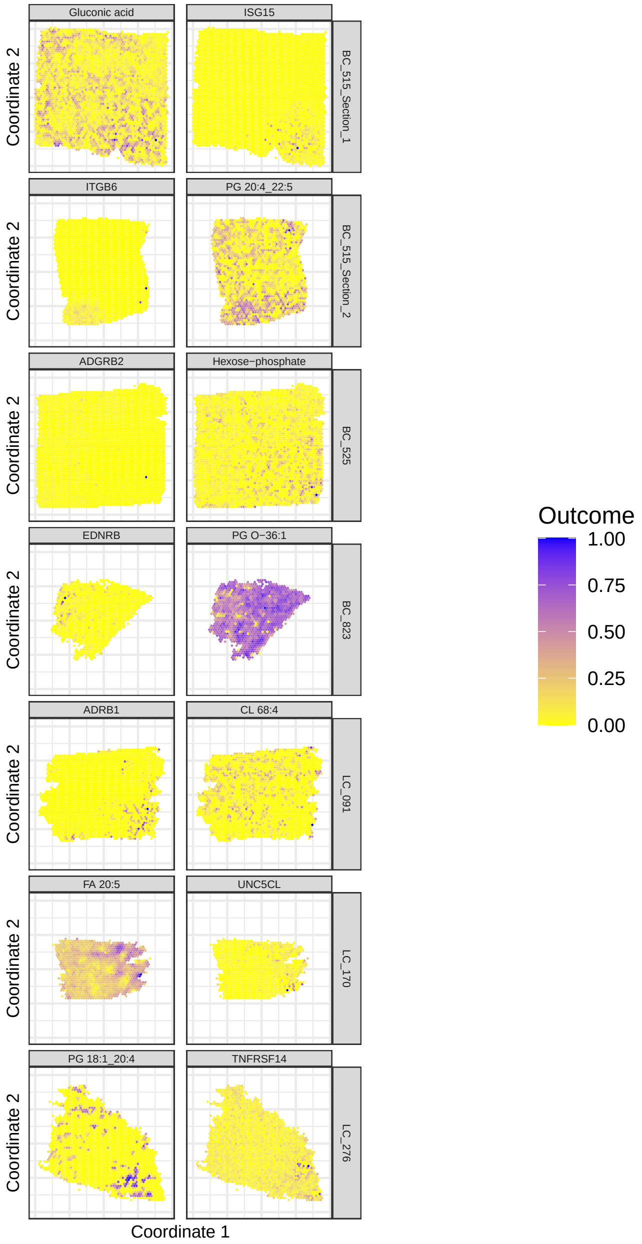

Fig S30: Gene and metabolite (columns) most significantly associated per section (rows) according to Moran's I (independence) in the Godfrey data. Colours reflect abundance, scaled to 0-1 for legibility.

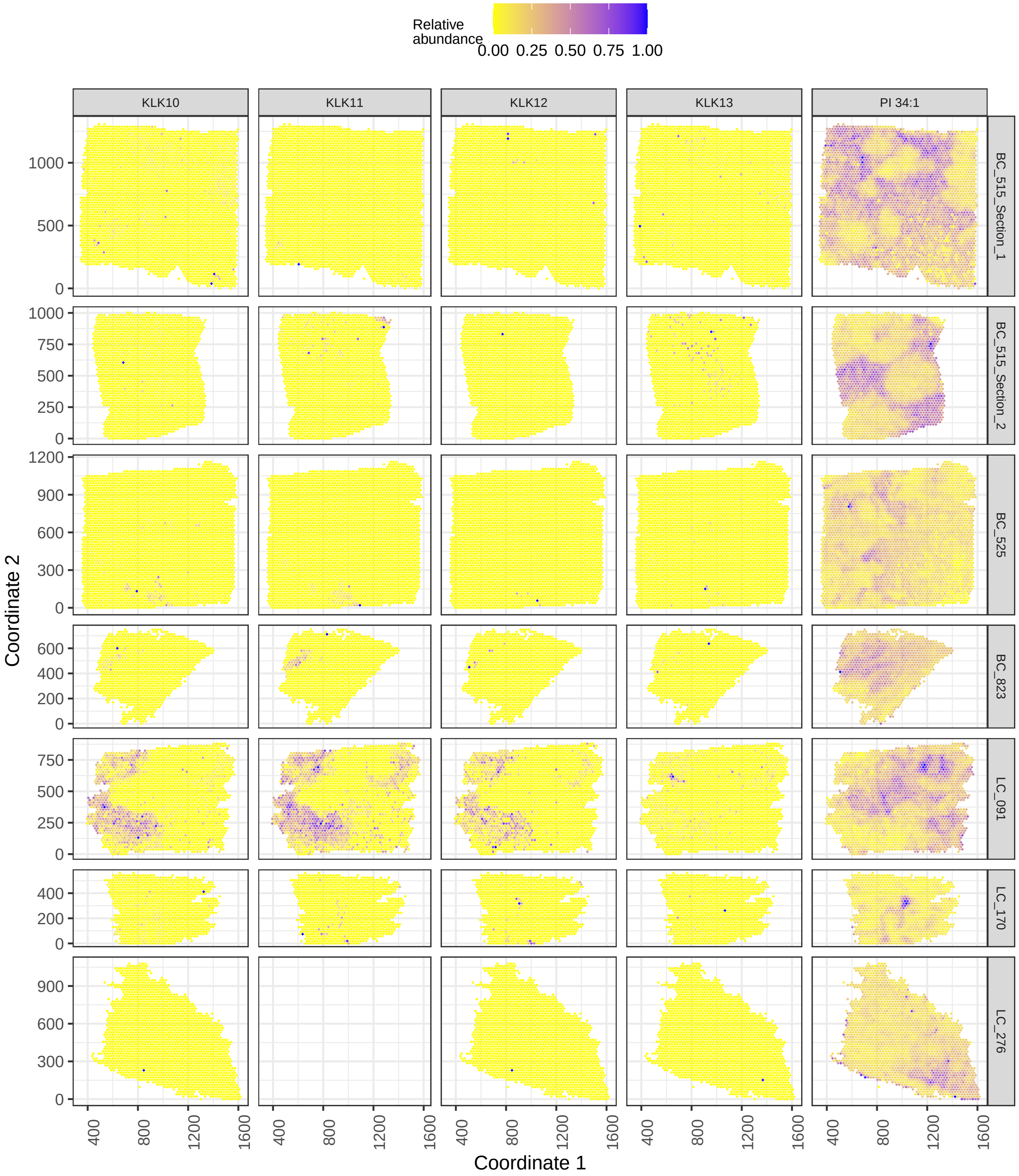

Fig S31: Relative abundance of PI 34:1 and the kallikrein transcripts most strongly associated with it (columns) in samples of the Godfrey data (rows). Colours reflect abundance, scaled to 0-1 for legibility. See Table S1 for full results.

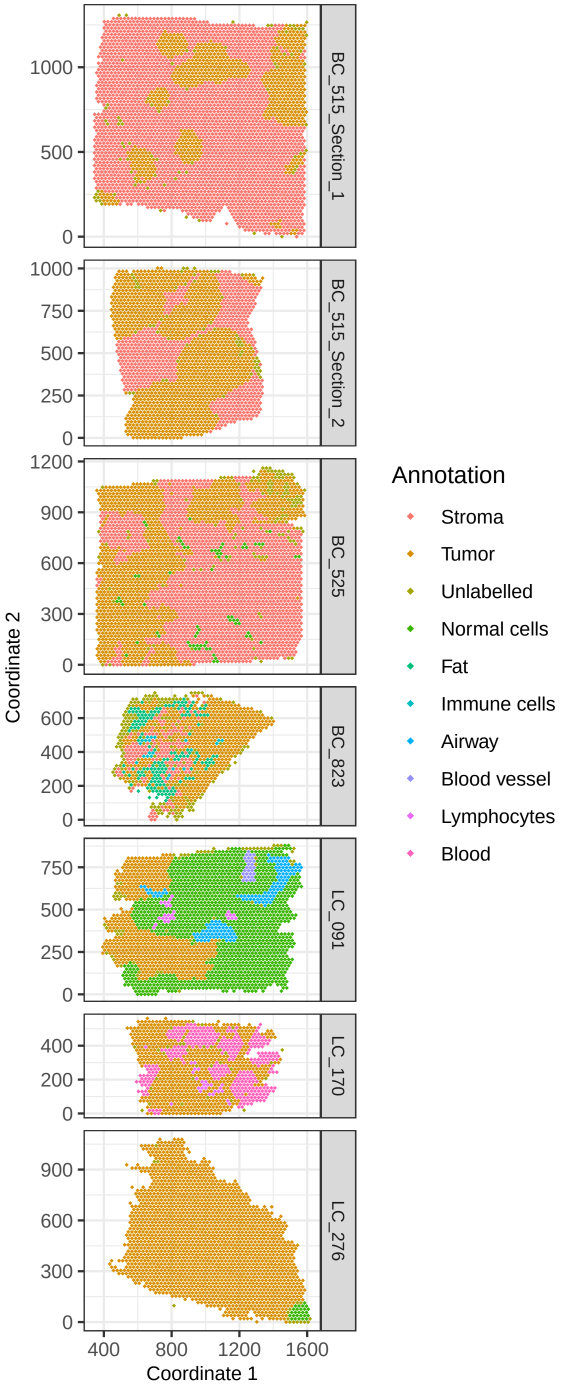

Fig S32: Godfrey data sections (rows) coloured by pathologists' annotation of the spots.

| Sample | Gene | Correlation | Adjusted.p.value |
|---|---|---|---|
| BC_515_Section_1 | KLK10 | -0.60 | 3.92e-08 |
| BC_515_Section_1 | KLK11 | -0.34 | 1.11e-04 |
| BC_515_Section_2 | KLK10 | -0.44 | 2.46e-07 |
| BC_515_Section_2 | KLK11 | -0.15 | 3.18e-07 |
| BC_515_Section_2 | KLK13 | -0.31 | 2.98e-02 |
| LC_091 | KLK10 | -0.58 | 4.45e-216 |
| LC_091 | KLK11 | -0.60 | 1.66e-192 |
| LC_091 | KLK12 | -0.57 | 3.24e-97 |
| LC_091 | KLK13 | -0.34 | 2.93e-21 |
| LC_170 | KLK10 | 0.25 | 4.88e-02 |

Table S1: Significant correlations of *KLK* genes with PI 34:1 estimated by GAMs with adjusted p-values for all Godfrey samples. Only KLK10 in BC_515_Section_1 and KLK13 in LC_091 are also significant according to Moran's I (independence).

##### 3.1.2 Multi-image analysis

In the original publication, the authors also look at which correlations are high in replicate sections of the same cancer type. They identify high correlations between *MUC1* and a series of phospholipids [7], but these are not significant after multiplicity correction in our analysis with GAMs (Fig S33).

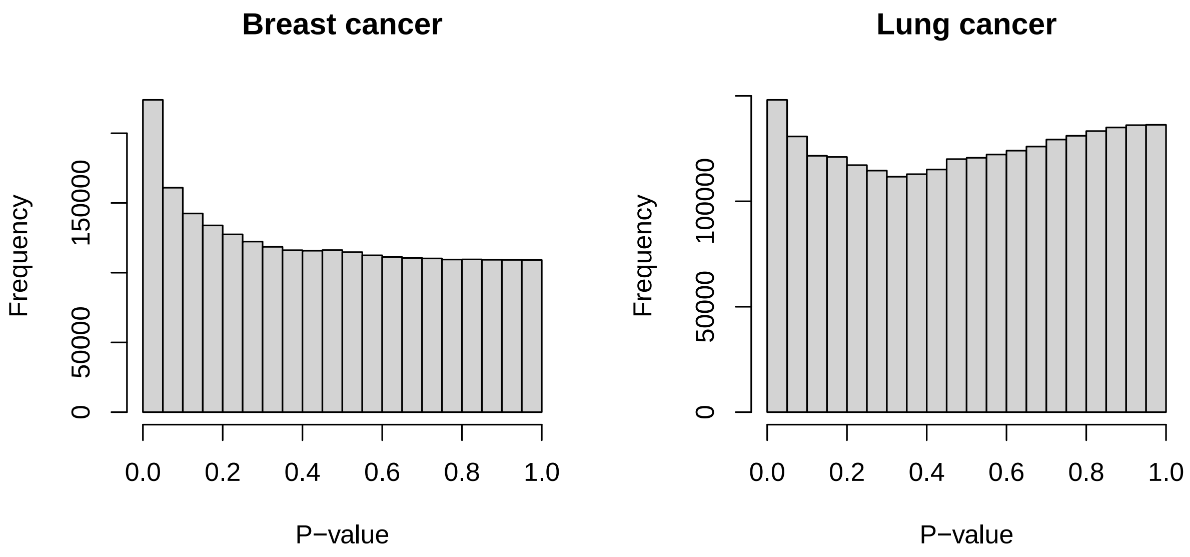

Fig S33: Histograms of p-values of GAMs of a replicated analysis per cancer type (plot titles) for the Godfrey data.

#### 3.2 Vicari et al. (2024)

This is a dataset on spatial transcriptomics and metabolomics on the same tissue sections from mouse and human brain [6]. Here we focus on the three mice were measured in triplicate. The first two triplets of mouse sections (V11L12-038 and V11L12-109) are from the striatum, the third (V11T16-085) is from the substantia nigra. The mice's brains were lesioned on one side through 6-hydroxydopamine (6-OHDA) injection to emulate Parkinson's disease. The authors aligned the images manually using the location information of the non-empty samples only with *STUtility* [28], but we used the *MAGPIE* package [29] to guide manual alignment using anchor points (available in supplementary material), the resulting alignment is shown in Fig S34. The spatial distributions of the spot-wise sums (library sizes and total ion counts) are shown in Fig S35, revealing spatial patterns for them.

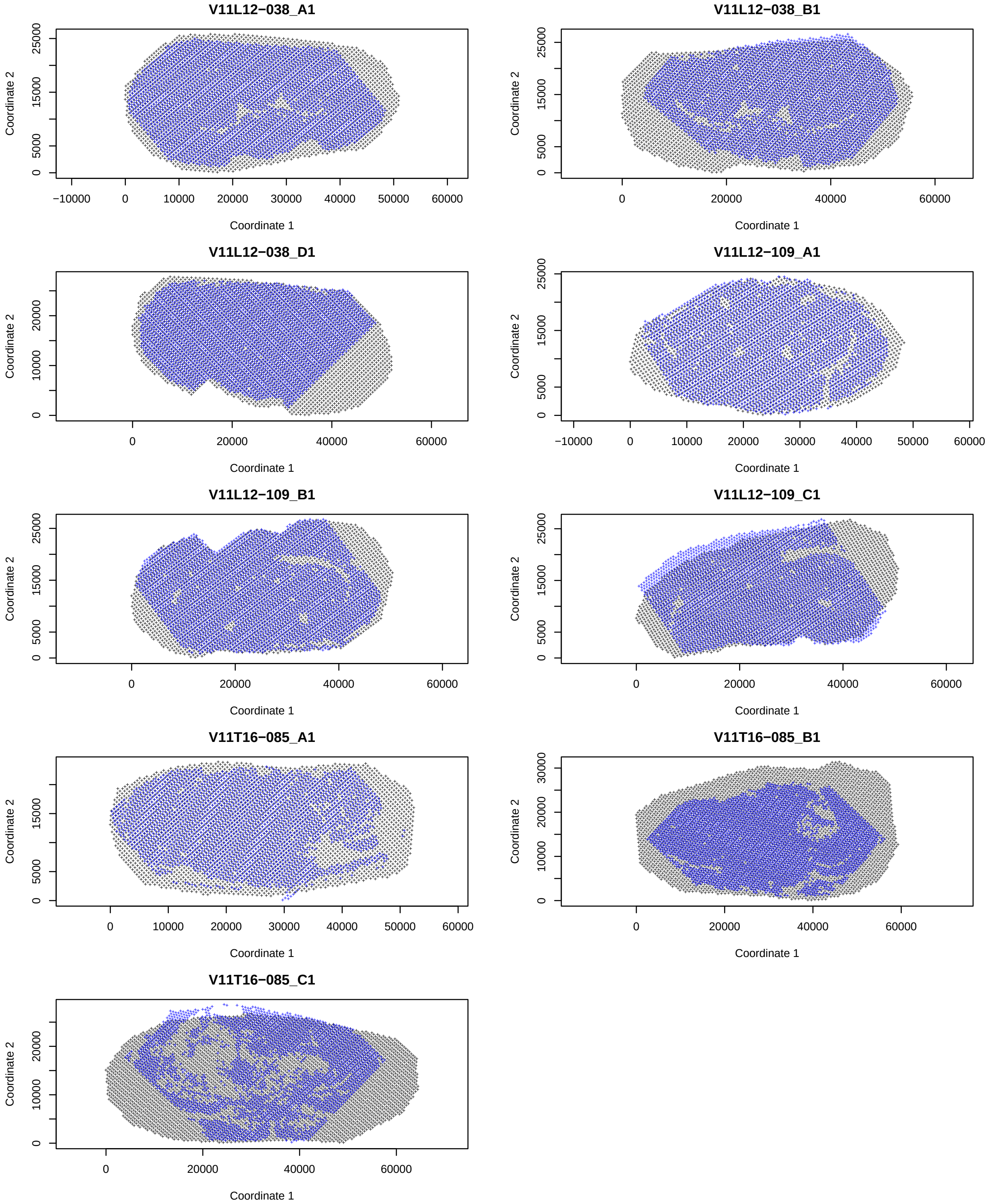

Fig S34: Images from Vicari et al. [6] aligned using *MAGPIE*. Metabolic spots are black, transcriptomic ones are blue.

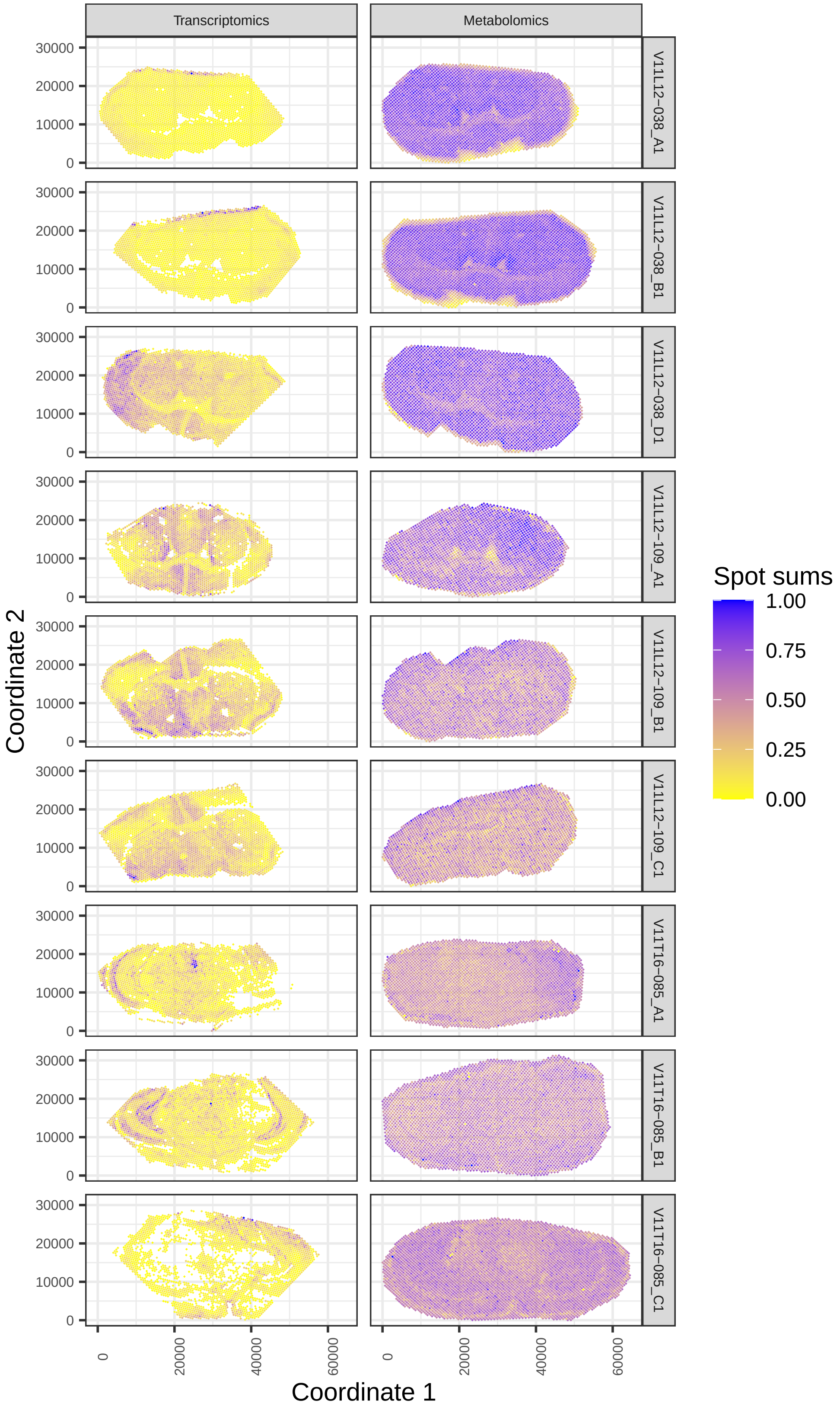

Fig S35: Spatial distribution of spot-wise sums (library sizes and total ion counts) from the transcriptomics respectively metabolomics modalities (columns) of the Vicari2024 data, for different sections (rows). All spot sums have been scaled to the [0,1] range per modality and omics type for legibility.

##### 3.2.1 Single-image analysis

**3.2.1.1 Comparison to authors' findings** In the original publication the authors used nearest neighbour distances to find a one-to-one mapping between metabolic and transcriptomic samples, for which then Pearson correlation is calculated [6]. The investigation focused on the metabolite dopamine, for which positive and negative correlations with a number of genes were found, but some were not confirmed by our analysis based on Moran's I (independence) or GAMs (Fig S36-S37).

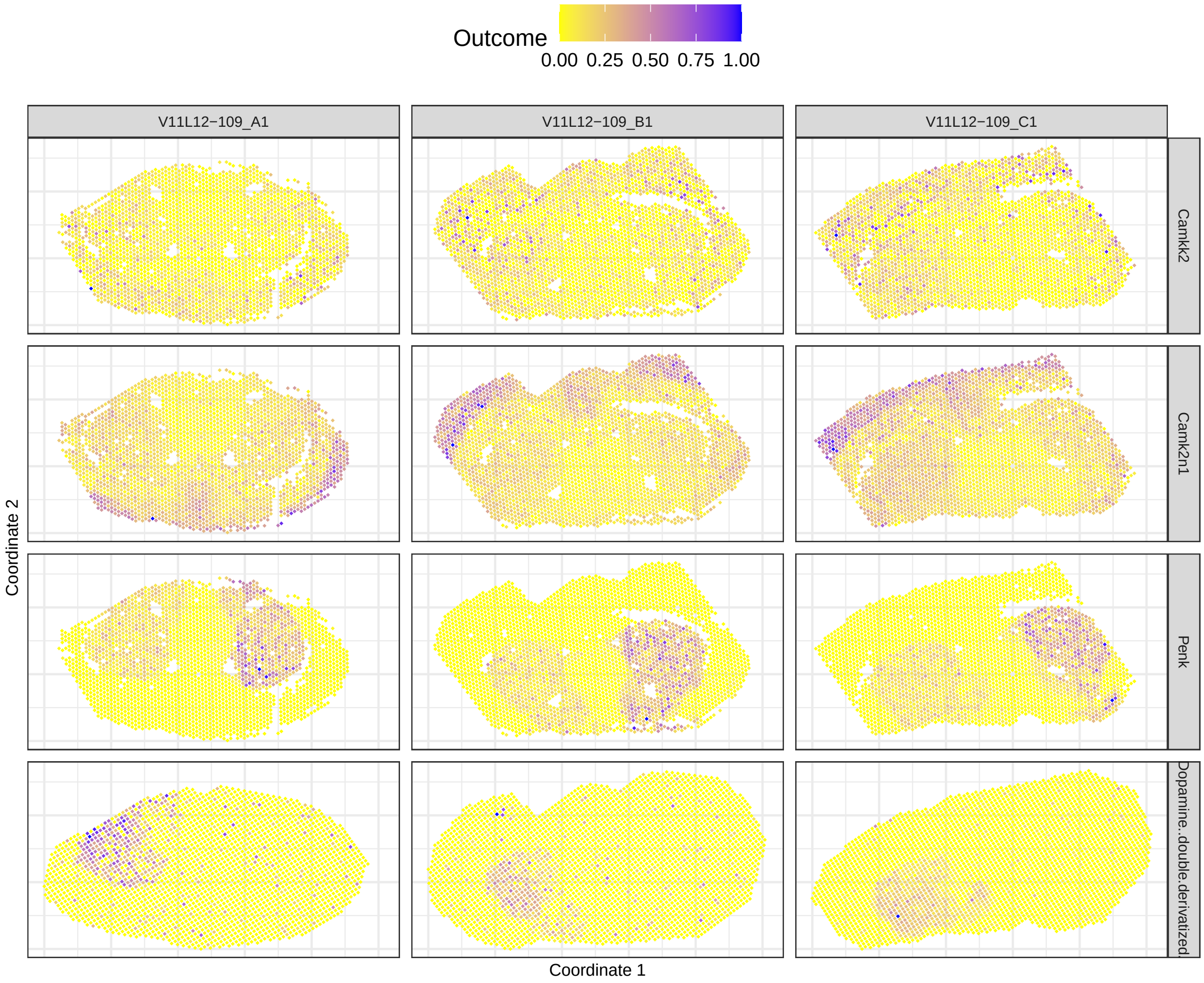

Fig S36: Genes (all but last row) found by the authors to be associated to dopamine (last row), but not significant for Moran's I (independence) or GAMs, in different sections of the striatum of the Vicari data where dopamine was detected (columns).

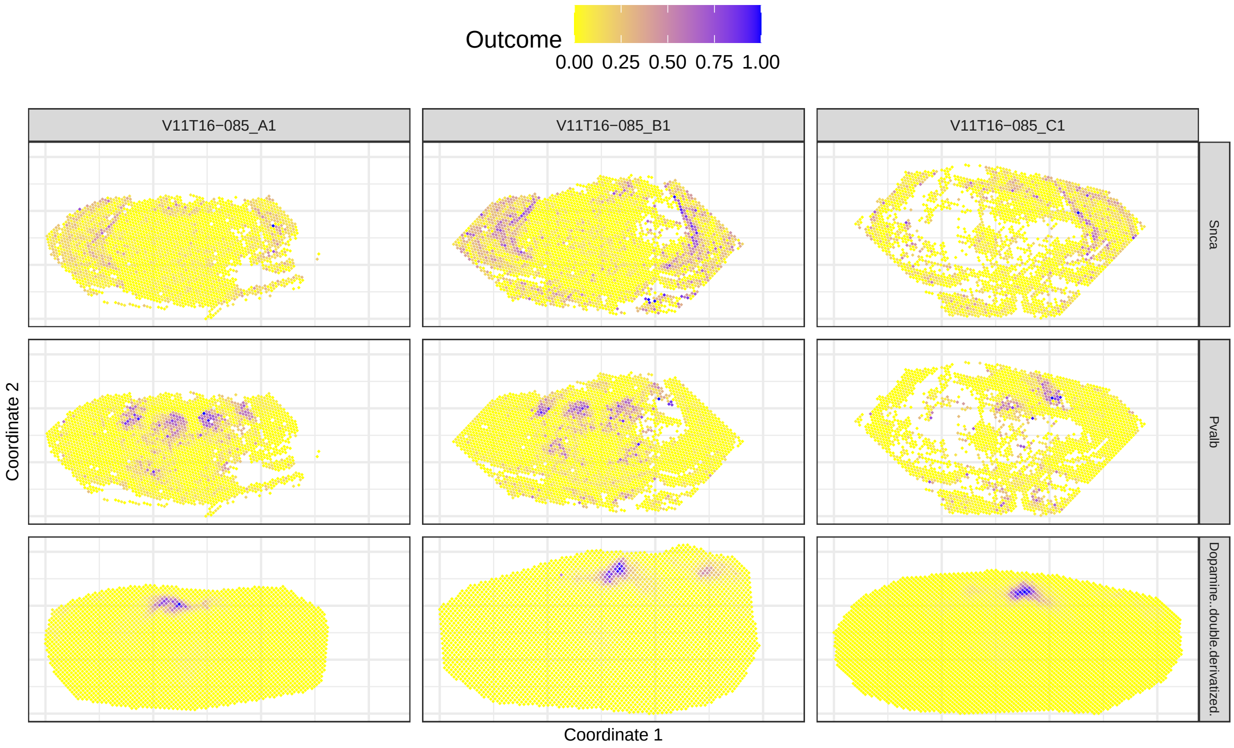

Fig S37: Genes (all but last row) for which both Moran's I (independence) and GAMs find a different association to dopamine (last row) than the authors, in different substantia nigra sections of the Vicari data (columns). *Snca* was found positively associated by the authors but negatively associated or not significant by GAMs or Moran's I (independence). *Pvalb* was found negatively associated by the authors but positively associated by GAMs and not significant by Moran's I (independence).

**3.2.1.2 New findings** The most significant findings are shown in S38-S39. The biologically interesting association between *Cspr1* and palmitoyl-L-carnitine is shown in Figure S40.

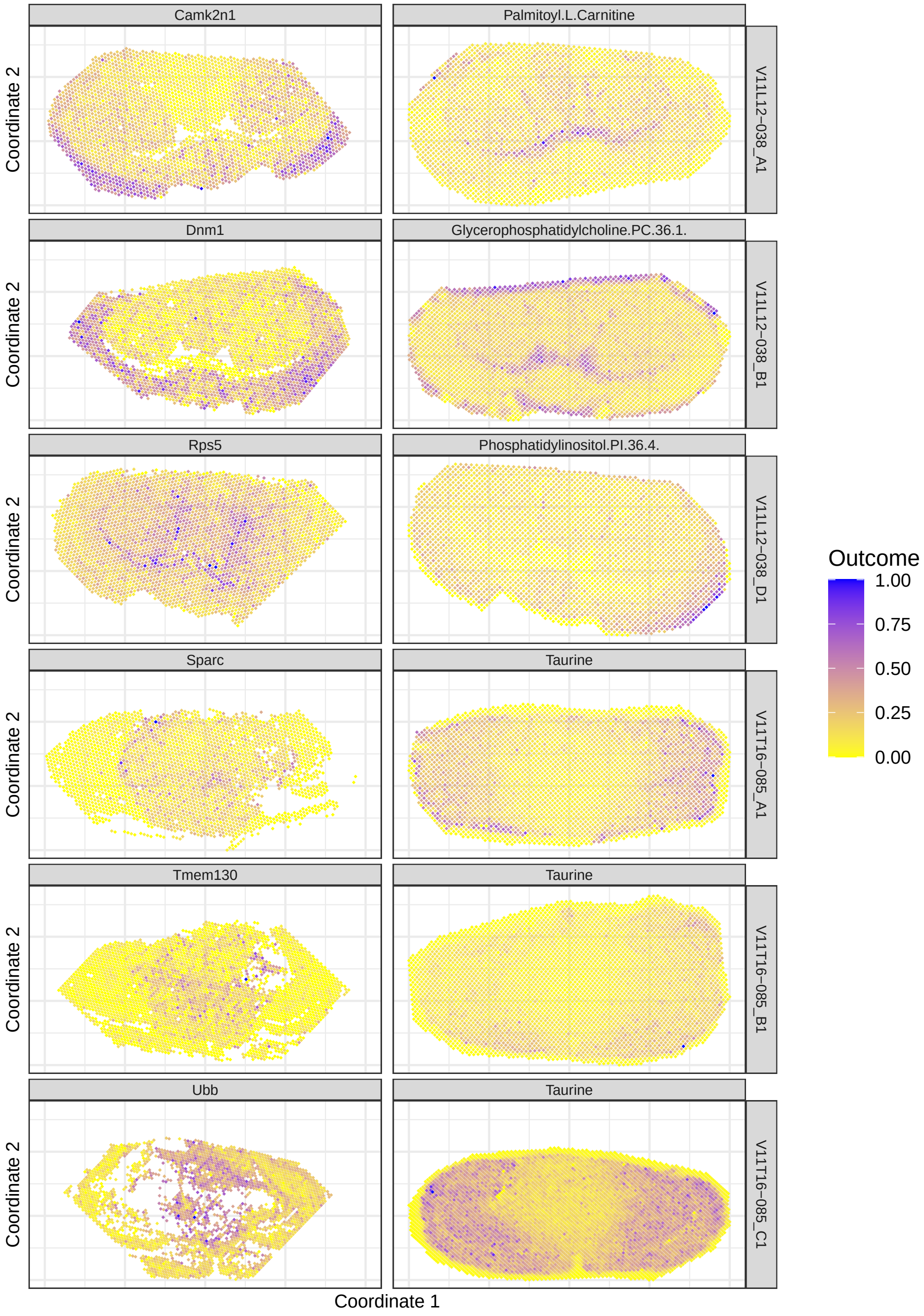

Fig S38: Gene-metabolite pairs with most significant negative correlation according to GAMs in all sections of the Vicari data that have significant negative correlations. None of these findings are significant according to Moran's I (independence).

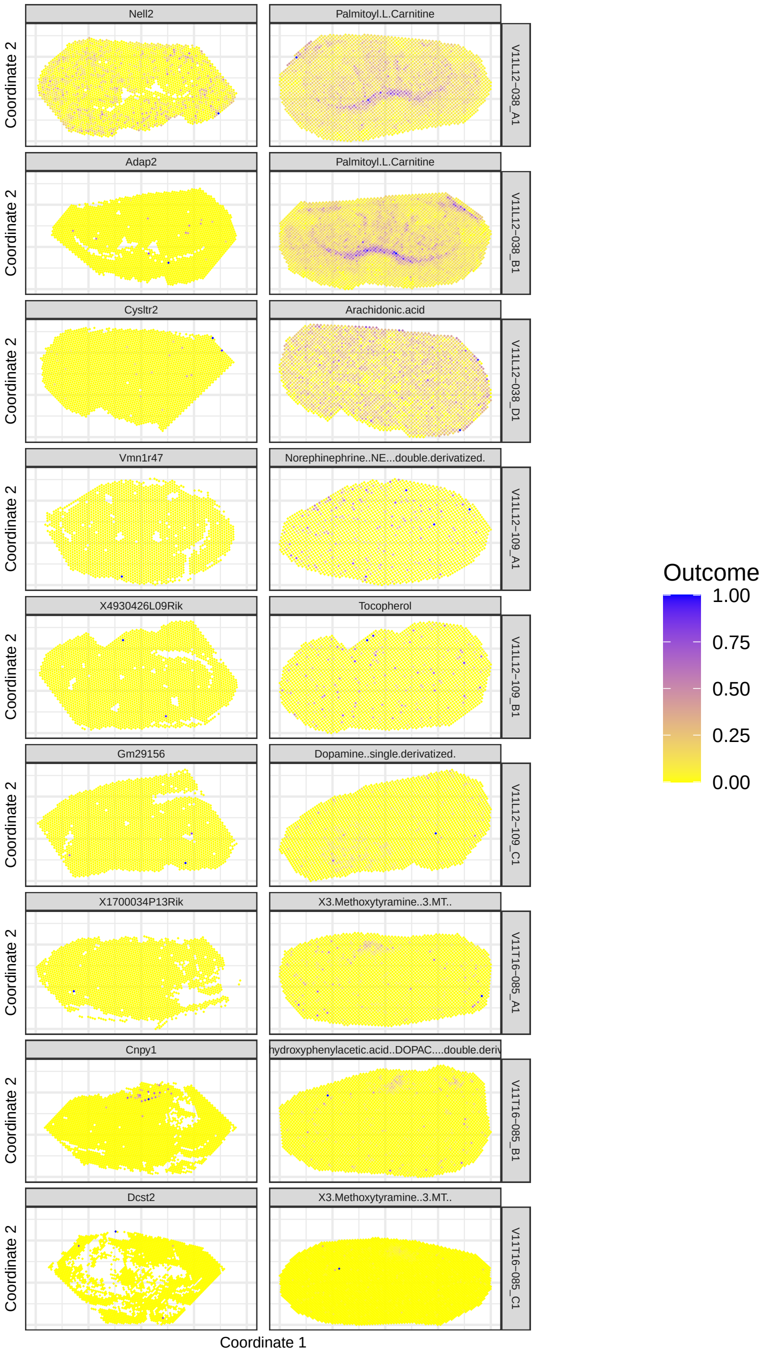

Fig S39: Gene-metabolite pairs most significant according to Moran's I in all sections of the Vicari data. Moran's I (independence) captures many very short-range associations of genes with metabolite hotspots, but this may also indicate lack of robustness to heteroskedasticity.

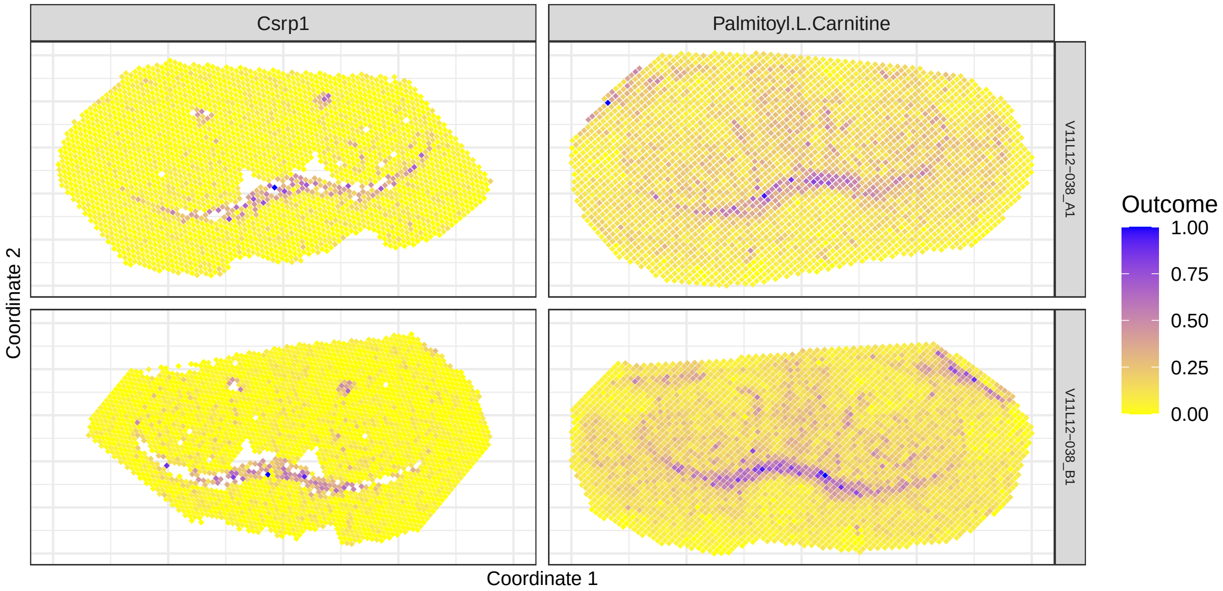

Fig S40: *Csrp1* transcript and palmitoyl-L-carnitine in striatum samples of the Vicari data (rows). This metabolite is only present in these two sections, where it is highly positively associated with Csrp1, with GAM correlation estimates and 0.67 and corresponding adjusted p-values and 5.8e-28 in samples V11L12-038_A1 and V11L12-038_B1, respectively. The associations are also significant according to Moran's I (independence).

##### 3.2.2 Multi-image analysis

We perform a multi-image analysis separately for the striatum and the substantia nigra sections, with a random effect for mouse for the former samples. The p-value distributions are shown in Fig S41.

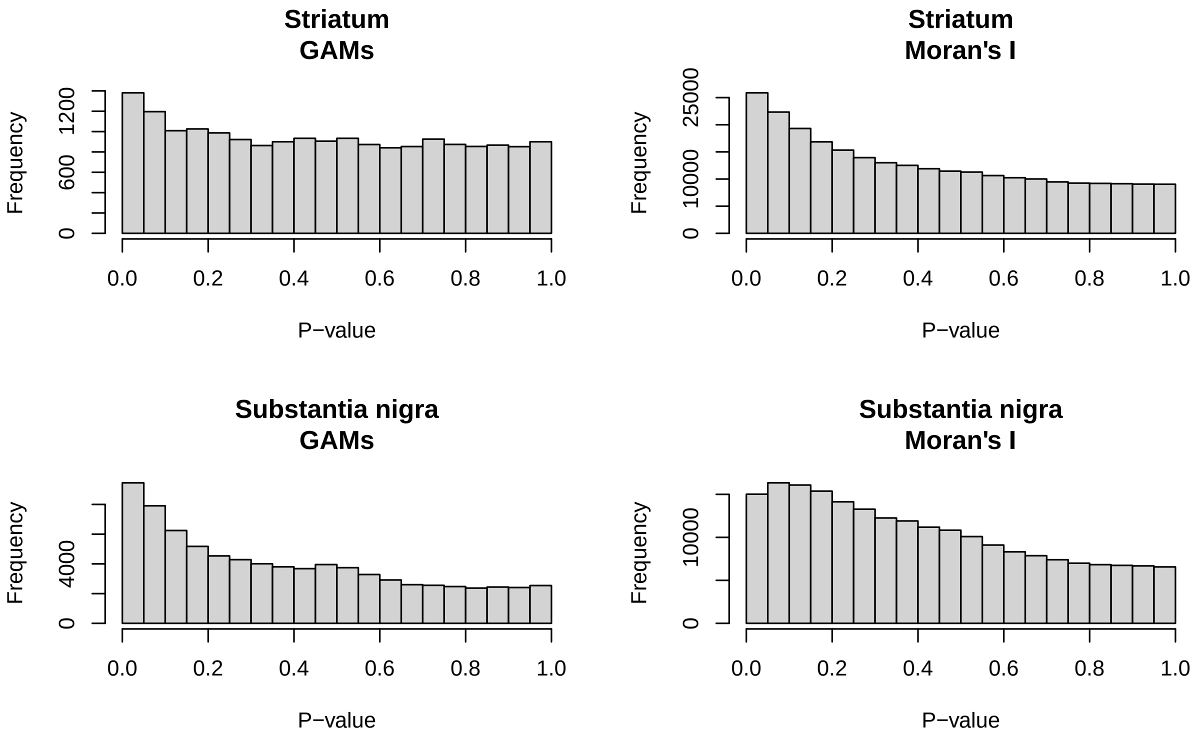

Fig S41: P-value distributions of replicated analysis of GAMs and Moran's I (independence) for striatum and substantia nigra samples (plot titles) in the Vicari data.

### 4 Recommendations

| Method | Strengths | Weaknesses | When to use |
|---|---|---|---|
| Methods with permutation or randomisation null distributions: *SpaGene*, *LIANA+*, Geary's C, Lee's L and some versions of bivariate Moran's I (*SpatialDM*, permutation, randomisation) | • Fast | • Geary's C and Lee's L require spot matching for differing resolutions<br>• Lack robustness to SAC, leading to spurious significances for bivariate associations | • Not recommended |
| Pearson correlation | • Fast<br>• Intuitive interpretation | • Requires spot matching for differing resolutions<br>• Yields spurious significances if SAC is present in both variables | • Not recommended |
| Modified t-test | • Intuitive interpretation: the quantity of interest is the Pearson correlation coefficient<br>• Robust to many types of SAC | • Requires spot matching for differing resolutions | • Data are homoskedastic or can be properly normalized<br>• Both modalities were measured at the same spots |
| Random shift test | • Applicable to any measure of spatial association<br>• Preserves SAC in permutation test | • Computationally demanding<br>• The p-values are discretized by the number of random shifts<br>• Limited power as some spatial association is retained in the random shifts | • Not recommended |
| Bivariate Moran's I (independence) [sbivar] | • Nonparametric<br>• Intuitive interpretation: closest equivalent of Pearson correlation under disjoint coordinates | • Weight matrix needs to be user defined, determining sensitivity to different types of spatial patterns<br>• Affected by heteroskedasticity | • Data are homoskedastic or can be properly normalized |
| Bivariate Gaussian processes (GPs) [sbivar] | • Robust to many types of SAC<br>• Amendable to other outcome distributions and link functions | • Very high computational demands<br>• Does not provide a measure of bivariate association strength, only a p-value<br>• Relies on parametric assumptions<br>• Complex mathematical derivation | • For single-image analysis<br>• A well-fitting outcome distribution is available or data can be properly normalized<br>• If computationally feasible |
| Generalized additive models (GAMs) [sbivar] | • Easily amendable to other outcome distributions and link functions<br>• Intuitive interpretation and visualisation<br>• Good power to detect moderate to long range differences | • Low power to detect short-range bivariate spatial association<br>• Relies on parametric assumptions<br>• Version accounting for residual SAC in a GAMM is very slow | • A well-fitting outcome distribution is available or data can be properly normalized<br>• If all spatial structure is captured by splines, use GAMS, otherwise GAMMs if computationally feasible |

Table S2: Methods to test for bivariate spatial association considered in this study, with strengths, weaknesses and our recommendation. [sbivar] indicates methods proposed or improved in this work and available in the *sbivar* R-package.

### 5 Software versions

```
## R version 4.6.1 (2026-06-24)
## Platform: x86_64-pc-linux-gnu
## Running under: Ubuntu 24.04.5 LTS
##
## Matrix products: default
## BLAS:   /usr/lib/x86_64-linux-gnu/openblas-pthread/libblas.so.3
## LAPACK: /usr/lib/x86_64-linux-gnu/openblas-pthread/libopenblasp-r0.3.26.so;  LAPACK version 3.12.0
##
## locale:
##  [1] LC_CTYPE=en_US.UTF-8       LC_NUMERIC=C
##  [3] LC_TIME=en_GB.UTF-8        LC_COLLATE=en_US.UTF-8
##  [5] LC_MONETARY=en_GB.UTF-8    LC_MESSAGES=en_US.UTF-8
##  [7] LC_PAPER=en_GB.UTF-8       LC_NAME=C
##  [9] LC_ADDRESS=C               LC_TELEPHONE=C
## [11] LC_MEASUREMENT=en_GB.UTF-8 LC_IDENTIFICATION=C
##
## time zone: Europe/Brussels
## tzcode source: system (glibc)
##
## attached base packages:
## [1] stats4    parallel  stats     graphics  grDevices utils     datasets
## [8] methods   base
##
## other attached packages:
##  [1] pizzarr_0.2.1          arrow_25.0.1           latticeExtra_0.6-31
##  [4] lattice_0.23-1         abind_1.4-8            lmerTest_3.2-1
##  [7] lme4_2.0-6             spatstat_3.6-2         spatstat.linnet_3.5-3
## [10] spatstat.model_3.7-2   rpart_4.1.27           spatstat.explore_3.8-3
## [13] spatstat.random_3.5-2  spatstat.geom_3.8-3    spatstat.univar_3.2-0
## [16] spatstat.data_3.1-9    jsonlite_2.0.0         bispdep_1.0-2
## [19] httr_1.4.9             nleqslv_3.3.7          fdrtool_1.2.18
## [22] smoppix_1.4.0          Seurat_5.5.1           SeuratObject_5.4.0
## [25] sp_2.2-3               sbivar_0.99.45         Cardinal_3.14.0
## [28] S4Vectors_0.50.3       ProtGenerics_1.44.0    BiocGenerics_0.58.1
## [31] generics_0.1.4         BiocParallel_1.46.0    anndata_0.8.0
## [34] rhdf5_2.56.1           SpaGene_0.1.0          gstat_2.1-6
## [37] peakRAM_1.0.2          NTSS_0.1.3             nlme_3.1-171
## [40] ggh4x_0.3.1            spdep_1.4-2            sf_1.1-3
## [43] spData_2.3.5           mvtnorm_1.4-2          stringr_1.6.0
## [46] SpatialPack_0.4-1      fastmatrix_0.6-6       RANN_2.6.3
## [49] Matrix_1.7-6           dplyr_1.2.1            patchwork_1.3.2
## [52] openxlsx_4.2.9         gplots_3.3.0           knitr_1.52
## [55] xtable_1.8-8           reshape2_1.4.5         ggplot2_4.0.3
##
## loaded via a namespace (and not attached):
##   [1] matrixStats_1.5.0           spatstat.sparse_3.2-0
##   [3] bitops_1.1-0                RColorBrewer_1.1-3
##   [5] numDeriv_2016.8-1.1         sctransform_0.4.3
##   [7] tools_4.6.1                 R6_2.6.1
##   [9] uwot_0.2.5                  mgcv_1.9-4
##  [11] rhdf5filters_1.24.1         withr_3.0.3
##  [13] splancs_2.01-45             gridExtra_2.3.1
```

```
##  [15] progressr_1.0.0              cli_3.6.6
##  [17] Biobase_2.72.0               fastDummies_1.7.6
##  [19] labeling_0.4.3               S7_0.2.2
##  [21] proxy_0.4-29                 ggridges_0.5.7
##  [23] pbapply_1.7-5                parallelly_1.48.0
##  [25] CardinalIO_1.10.0            FNN_1.1.4.1
##  [27] combinat_0.0-9               gtools_3.9.5
##  [29] ica_1.0-3                    zip_3.0.2
##  [31] interp_1.1-6                 lifecycle_1.0.5
##  [33] yaml_2.3.12                  SummarizedExperiment_1.42.0
##  [35] SparseArray_1.12.2           Rtsne_0.17
##  [37] grid_4.6.1                   promises_1.5.0
##  [39] miniUI_0.1.2                 cowplot_1.2.0
##  [41] magick_2.9.1                 pillar_1.11.1
##  [43] GenomicRanges_1.64.0         tcltk_4.6.1
##  [45] rjson_0.2.23                 boot_1.3-32
##  [47] spacetime_1.3-4              future.apply_1.20.2
##  [49] codetools_0.2-20             wk_0.9.5
##  [51] glue_1.8.1                   data.table_1.18.6.1
##  [53] MultiAssayExperiment_1.38.0 vctrs_0.7.3
##  [55] png_0.1-9                    spam_2.11-4
##  [57] Rdpack_2.6.6                 gtable_0.3.6
##  [59] assertthat_0.2.1             cachem_1.1.0
##  [61] zigg_0.0.2                   ks_1.15.3
##  [63] xfun_0.61                    rbibutils_2.4.1
##  [65] S4Arrays_1.12.0              mime_0.13
##  [67] Rfast_2.1.5.2                Seqinfo_1.2.0
##  [69] pracma_2.4.6                 reformulas_0.4.4
##  [71] survival_3.8-12              SingleCellExperiment_1.34.0
##  [73] units_1.0-1                  fitdistrplus_1.2-6
##  [75] ROCR_1.0-12                  xts_0.14.3
##  [77] bit64_4.8.6                  matter_2.14.0
##  [79] RcppAnnoy_0.0.23             geoR_1.9-6
##  [81] irlba_2.3.7                  KernSmooth_2.23-27
##  [83] otel_0.2.0                   DBI_1.3.0
##  [85] tidyselect_1.2.1             bit_4.6.0
##  [87] compiler_4.6.1               ontologyIndex_2.12
##  [89] DelayedArray_0.38.2          plotly_4.12.1
##  [91] stringfish_0.19.2            scales_1.4.0
##  [93] caTools_1.18.4               classInt_0.4-11
##  [95] lmtest_0.9-40                SpatialExperiment_1.22.0
##  [97] digest_0.6.39                goftest_1.2-3
##  [99] spatstat.utils_3.2-5         minqa_1.2.8
## [101] rmarkdown_2.32               XVector_0.52.0
## [103] jpeg_0.1-11                  htmltools_0.5.9
## [105] pkgconfig_2.0.3              MatrixGenerics_1.24.0
## [107] fastmap_1.2.0                rlang_1.3.0
## [109] htmlwidgets_1.6.4            shiny_1.14.0
## [111] farver_2.1.2                 zoo_1.9-0
## [113] mclust_6.1.3                 magrittr_2.0.5
## [115] s2_1.1.12                    dotCall64_1.2
## [117] Rhdf5lib_2.0.0               Rcpp_1.1.2
## [119] reticulate_1.47.0            stringi_1.8.9
## [121] MASS_7.3-66                  plyr_1.8.9
```

```
## [123] listenv_1.0.0              ggrepel_0.9.8
## [125] deldir_2.0-4               splines_4.6.1
## [127] tensor_1.5.1               GET_1.0-9
## [129] igraph_2.3.3               RcppHNSW_0.7.0
## [131] qs2_0.3.1                  evaluate_1.0.5
## [133] RcppParallel_6.2.1         nloptr_2.2.1
## [135] httpuv_1.6.17              tidyr_1.3.2
## [137] purrr_1.2.2                polyclip_1.10-7
## [139] scattermore_1.2            future_1.75.0
## [141] e1071_1.7-17               RSpectra_0.16-2
## [143] later_1.4.8                viridisLite_0.4.3
## [145] class_7.3-24               intervals_0.15.5
## [147] tibble_3.3.1               memoise_2.0.1
## [149] IRanges_2.46.0             cluster_2.1.8.2
## [151] globals_0.19.1             concaveman_1.2.0
```

### References

[1] R. L. Czaplewski. *Expected value and variance of Moran's bivariate spatial autocorrelation statistic for a permutation test.* Vol. 309. US Department of Agriculture, Forest Service, Rocky Mountain Forest and Range Experiment Station, 1993.

[2] Z. Li, T. Wang, P. Liu, and Y. Huang. "SpatialDM for rapid identification of spatially co-expressed ligand-receptor and revealing cell-cell communication patterns." In: *Nat. Commun.* 14.1 (2023), p. 3995.

[3] D. DeTomaso and N. Yosef. "Hotspot identifies informative gene modules across modalities of single-cell genomics". In: *Cell Syst.* 12.5 (2021), pp. 446 –456.

[4] B. F. Miller, D. Bambah-Mukku, C. Dulac, X. Zhuang, and J. Fan. "Characterizing spatial gene expression heterogeneity in spatially resolved single-cell transcriptomic data with nonuniform cellular densities." In: *Genome Res.* 31.10 (2021), pp. 1843 –1855.

[5] D. Wartenberg. "Multivariate Spatial Correlation: A Method for Exploratory Geographical Analysis". In: *Geographical Analysis* 17.4 (1985), pp. 263 –283.

[6] M. Vicari, R. Mirzazadeh, A. Nilsson, R. Shariatgorji, P. Bjärterot, L. Larsson, et al. "Spatial multimodal analysis of transcriptomes and metabolomes in tissues." In: *Nat. Biotechnol.* 42.7 (2024), pp. 1046 –1050.

[7] T. M. Godfrey, Y. Shanneik, W. Zhang, T. Tran, N. Verbeeck, N. H. Patterson, et al. "Integrating Ambient Ionization Mass Spectrometry Imaging and Spatial Transcriptomics on the Same Cancer Tissues to Identify RNA–Metabolite Correlations". In: *Angew. Chem. Int. Ed.* (2025).

[8] V. M. Ravi, P. Will, J. Kueckelhaus, N. Sun, K. Joseph, H. Salié, et al. "Spatially resolved multi-omics deciphers bidirectional tumor-host interdependence in glioblastoma." In: *Cancer Cell* 40.6 (2022), 639 –655 –e13.

[9] J. J. Lennon. "Red-Shifts and Red Herrings in Geographical Ecology". In: *Ecography* 23.1 (2000), pp. 101 –113.

[10] P. Clifford, S. Richardson, and D. Hemon. "Assessing the Significance of the Correlation between Two Spatial Processes". In: *Biometrics* 45.1 (1989), pp. 123 –134.

[11] P. Dutilleul, P. Clifford, S. Richardson, and D. Hemon. "Modifying the t test for assessing the correlation between two spatial processes". In: *Biometrics* (1993), pp. 305 –314.

[12] P. Dutilleul, B. Pelletier, and G. Alpargu. "Modified F tests for assessing the multiple correlation between one spatial process and several others". In: *J. Stat. Plan. Inference* 138.5 (2008), pp. 1402 –1415.

[13] Q. Liu, C.-Y. Hsu, and Y. Shyr. "Scalable and model-free detection of spatial patterns and colocalization." In: *Genome Res.* 32.9 (2022), pp. 1736 –1745.

[14] S. Seal and B. Neelon. "SpaceBF: Spatial coexpression analysis using Bayesian Fused approaches in spatial omics datasets". In: *Gigascience* 15 (2026).

[15] S.-I. Lee. "Developing a bivariate spatial association measure: An integration of Pearson's r and Moran's I". In: *Journal of Geographical Systems* 3.4 (2001), pp. 369 –385.

[16] R. S. Bivand and D. W. S. Wong. “Comparing implementations of global and local indicators of spatial association”. In: *Test* 27.3 (2018), pp. 716 –748.
[17] T. Mrkvička, J. Dvořák, J. A. González, and J. Mateu. “Revisiting the random shift approach for testing in spatial statistics”. In: *Spat. Stat.* 42 (2021), p. 100430.
[18] G. I. Ridder, O. J. Hardy, and O. Ovaskainen. “Generating spatially realistic environmental null models with the shift-and-rotate approach helps evaluate false positives in species distribution modelling”. In: *Methods in Ecology and Evolution* 15.12 (2024), pp. 2331 –2342.
[19] G. Matheron. “Principles of geostatistics”. In: *Economic Geology* 58.8 (1963), pp. 1246 –1266.
[20] E. J. Pebesma. “Multivariable geostatistics in S: The gstat package”. In: *Computers & Geosciences* 30.7 (2004), pp. 683 –691.
[21] H. H. Kelejian and I. R. Prucha. “On the asymptotic distribution of the Moran I test statistic with applications”. In: *J. Econom.* 104.2 (2001), pp. 219 –257.
[22] B. Phipson and G. K. Smyth. “Permutation P-values should never be zero: Calculating exact P-values when permutations are randomly drawn.” In: *Stat. Appl. Genet. Mol. Biol.* 9 (2010), Article39.
[23] D. Zhang and X. Lin. “Hypothesis testing in semiparametric additive mixed models”. In: *Biostatistics* 4.1 (2003), pp. 57 –74.
[24] I. Kats, R. Vento-Tormo, and O. Stegle. “SpatialDE2: Fast and localized variance component analysis of spatial transcriptomics”. In: *Biorxiv* (2021), pp. 2021 –2010.
[25] J. C. Pinheiro and D. M. Bates. *Mixed-Effects Models in S and S-PLUS*. Vol. 100. Springer, 2000, pp. 100 –461.
[26] J. Gardner, G. Pleiss, K. Q. Weinberger, D. Bindel, and A. G. Wilson. “Gpytorch: Blackbox matrix-matrix gaussian process inference with gpu acceleration”. In: *Advances in neural information processing systems* 31 (2018).
[27] D. Bates, M. Mächler, B. Bolker, and S. Walker. “Fitting Linear Mixed-Effects Models Using lme4”. In: *J. Stat. Softw.* 67.1 (2015), pp. 1 –48.
[28] J. Bergenstråhle, L. Larsson, and J. Lundeberg. “Seamless integration of image and molecular analysis for spatial transcriptomics workflows”. In: *BMC Genomics* 21.1 (2020), p. 482.
[29] E. C. Williams, L. Franzén, M. O. Lindvall, G. Hamm, S. Oag, M. M. Majumder, et al. “Spatially resolved integrative analysis of transcriptomic and metabolomic changes in tissue injury studies.” In: *Nat. Commun.* 17.1 (2026), p. 205.
